# Light-up nanostructures with allosteric DNA mimics of GFP

**DOI:** 10.1101/2025.07.10.664110

**Authors:** Tianqing Zhang, Xinmin Qian, Jiayi Zhang, Huangchen Cui, Tanxi Bai, Wen Wang, Yuxin Wang, Zhaoli Gao, Da Han, Wenwen Zeng, Bryan Wei

## Abstract

In this study, we present a systematic approach for the rational design of synthetic allosteric DNA aptamers. This methodology enables precise control over the allosteric ON-OFF transition in fluorescent DNA aptamers, allowing for the engineering of aptamers with highly tunable fluorescent properties. When combined with toehold-mediated strand displacement, we have developed a series of allosteric aptamers in which the target sequence functions as a specific allosteric modulator. Furthermore, these aptamers have been applied in synthetic DNA computing and in the construction of responsive nanostructures that light up upon activation.

## 1. Introduction

Throughout millions of years of evolution, allostery has become a highly efficient and adaptable mechanism for molecular stimulus response, regulating many essential life processes such as gene expression^[1]^ and signal transduction^[2]^. In allosteric regulation, the association between a biological macromolecule (typically a protein^[3-4]^ or RNA^[5]^) and its substrate is controlled by allosteric modulators, which act as activators^[3]^ or inhibitors^[6]^, resulting in either an up-regulated or down-regulated activity, respectively. Over the past two decades, allosteric regulation has garnered increasing interest across a broad research community, primarily due to its potential in developing allosteric drugs^[7-8]^ and allosteric antibodies^[9-10]^. These offer promising therapeutic advantages for some of the most difficult-to-treat human neurological and psychiatric disorders^[11]^. Due to the inherently limited programmability of protein dynamics, the ability to rationally design synthetic protein molecules with allosteric control capabilities remains constrained. In contrast, synthetic nucleic acid constructs, benefiting from their exceptional programmability^[12-16]^, have emerged as ideal molecules for rational allosteric design^[17-20]^. By applying well-established design principles of synthetic nucleic acids and the extensive toolkits of DNA nanotechnology^[21-25]^, engineered DNA constructs can be designed to recognize a variety of specific modulator—including biological^[26-27]^, chemical^[28-29]^, and even physical ones^[30-31]^—with high precision and functional control. Consequently, DNA-based allosteric systems represent a powerful strategy for various biomolecular detection tasks, offering levels of complexity and controllability that surpass those of their natural counterparts.

In this study, we successfully designed a series of allosteric nano-switches (Fig. 1) based on the recently reported fluorescent DNA aptamer—Lettuce^[32]^. Lettuce functions as a GFP mimic, producing green fluorescence upon binding to its ligand DFHBI-1T. DFHBI-1T is a derivative of the fluorophore HBI (4-hydroxybenzlidene imidazolinone) in GFP which is formed from an autocatalytic intramolecular cyclization of three residues in the nascent protein, Ser^65^-Tyr^66^-Gly^67^. The DNA lettuce aptamer comprises a core DFHBI-1T binding pocket, whose formation primarily depends on the stability of two flanking stems. Once in the correct conformation, the binding pocket can quickly bind and activate the fluorescence of DFHBI-1T fluorophore under appropriate excitation conditions. We designed and constructed split aptamers^[33-34]^ whose assembly is mediated by a stable constituent three-arm junction^[35]^, serving as the basic allosteric nano-switches. Prior to three-arm junction formation, the split aptamer adopts a loose conformation unable to form a complete binding pocket and bind to its ligand. Upon regulation by a target strand acting as an external allosteric modulator, the three-arm junction forms, leading to the reconstitution of the integral binding pocket and subsequent stable interaction with DFHBI-1T, thereby enabling fluorescence emission.

**Fig. 1.**
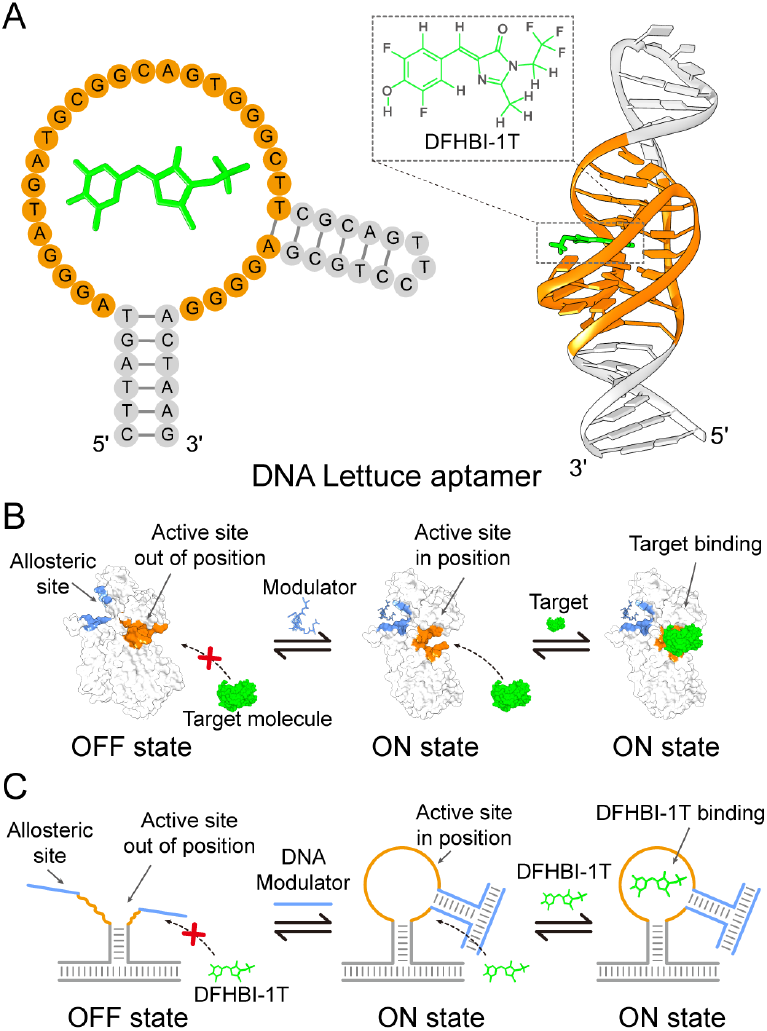
Allosteric nano-switches based on fluorescent DNA aptamer—Lettuce. A, strand diagram and 3D structure (PDB ID: 8FHX) of DNA Lettuce aptamer, sequence in orange depicts the binding pocket of aptamer ligand—DFHBI-1T. **B**, Schematics of allosteric regulation. The conformation switching from the blocked state (OFF state) to the open state (ON state) is triggered upon allosteric modulator binding. **C**, DNA-based allosteric nano-switches regulated by DNA modulators.

Through systematic screening of a set of synthetic DNA nano-switches, we specified a clear-cut fluorescent ON-OFF transition between well-defined conformational states. Subsequently, we enabled reversible regulation of fluorescent output by integrating toehold-mediated strand displacement into our design framework. By extending this strategy to incorporate multiple allosteric modulators, we successfully implemented a set of basic Boolean logic operations and their complex combinations. Finally, due to their excellent structure-dependent fluorescent properties, we integrated the allosteric DNA aptamers into light-up DNA nanostructures to facilitate characterization of nanostructure morphologies.

## 2. Results

### 2.1 Rational design and screening of allosteric fluorescent DNA nano-switches

We carefully specified ON and OFF allosteric states of fluorescent DNA nano-switch regulated by DNA modulators. As shown in Fig. 2A, the DNA Lettuce aptamer comprises three structural domains: two double-helical stems (P1 and P2, depicted in gray) and a ligand binding pocket (depicted in orange) positioned between P1 stem and P2 stem. With multiple guanines embedded in its sequence, the binding pocket forms a two-layer antiparallel G-quadruplex structure^[32]^. We hypothesize that stable formation of both P1 and P2 is essential for the proper conformation and function of the binding pocket. Specifically, stable P1 or P2 would reinforce the structural integrity of the binding pocket and enable fluorescence, whereas unstable P1 and P2 compromise fluorescence. To test the hypothesis, we designed nano-switches with extended P1 and P2 stems of varying lengths, regulated by a DNA modulator strand for allosteric control. A three-arm junction was incorporated into the nano-switch, where P1 (or P2) serves as one arm (Fig. 2B and D, Supplementary Figs. S1 and S4), with its stability dependent on the formation of the other two arms. The formation of the other two arms is regulated by a modulator strand. Without modulator strand, the P1 stem remains unstable, maintaining the binding pocket of the nano-switch in OFF state. After allosteric transition triggered by the modulator strand, stable P1 stem formation switches the nano-switch to ON state and induces fluorescence.

**Fig. 2.**
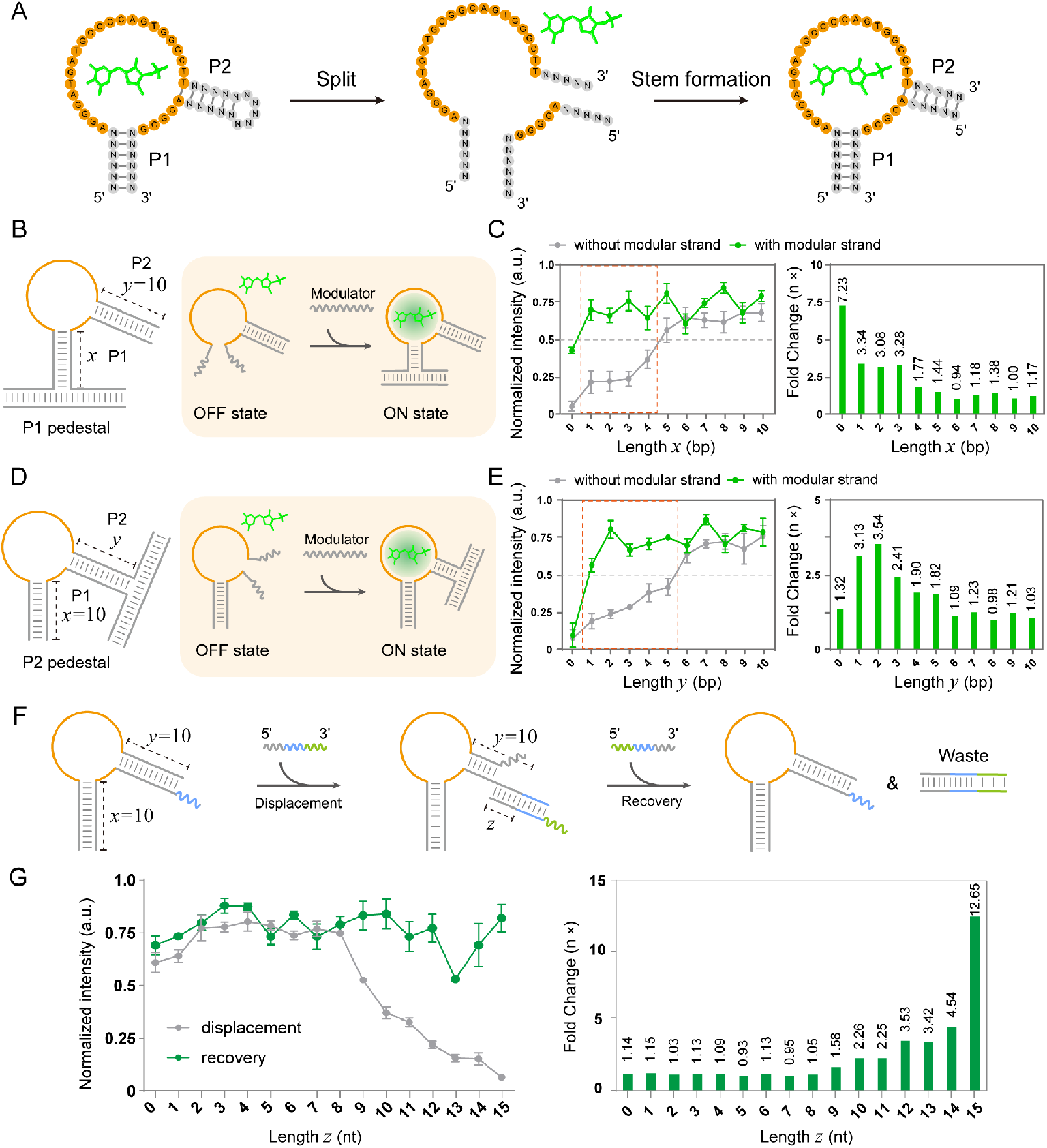
Design and screening of DNA allosteric aptamers. **A**, Strand diagram of a split DNA Lettuce aptamer. Sequence in orange depicts the binding pocket of aptamer ligand—DFHBI-1T. **B-C**. Design schematics (**B**) and screening results (**C**) of the allosteric nano-switch regulated by P1 formation. **D-E**. Design schematics (**D**) and screening results (**E**) of the allosteric nano-switch regulated by P2 formation. **F-G**. Design schematics (**F**) and screening results (**G**) of the reverse allosteric regulation. Errors bars, mean ± SEM.

We first designed two extended tails (15-bp) on P1 stem. These tails formed two arms upon collectively binding a 30-nt modulator strand, thereby forming a full three-arm junction with P1. With P2 fixed at 10 bp, we screened P1 variants of gradient lengths (0 to 10-bp) to evaluate the ON-OFF transition between allosteric states regulated by the same modulator strand. Systematic screening and quantitative analysis of fluorescence intensity measurements enabled the identification of optimal variants (Fig. 2C, Supplementary Table. S3).

While no transition was observed for seven negative variants, four positive variants (*x*=1, 2, 3, 4) exhibited successful ON-OFF fluorescence transition upon modulator binding. Initially, these four variants showed very weak fluorescence (normalized intensity below 0.5) without modulator. However, strong fluorescence (normalized intensity above 0.5) was detected upon modulator binding, marking a clear allosteric transition. In contrast, variants with P1 length ≥ 5 nt (*x*=5, 6, 7, 8, 9, 10) displayed a permanent ON state due to a stable P1 stem even without modulator binding (no three-arm junction). This ambiguous ON-OFF state rendered allosteric regulation unattainable. Finally, variants lacking P1 stem (*x*=0) maintained weak fluorescence intensity regardless of the presence of modulators, also preventing allosteric regulation. This absence of regulation might be attributed to disruption of the binding pocket conformation by the arm junction. The results are generally consistent with our thermodynamic simulation results (Supplementary Figs. S2-3 and Table S1), the high value of changes of standard free energy (ΔΔG ∼ 6 kcal/mol) and low switching equilibrium constant (Ks < E^-5^) of each successful allosteric design (*x*=1, 2, 3, 4) demonstrate the distinct transition between ON and OFF state.

With a similar rationale, we screened a set of P2 variants with gradient lengths (0 to 10-bp) to evaluate the modulator-regulated ON-OFF transition between allosteric states (Fig. 2E, Supplementary Table. S4). Five positive variants (*y*=1, 2, 3, 4, 5) were identified, showing successful fluorescence transition upon modulator binding. For these variants, only weak fluorescence (normalized intensity below 0.5) was detected without modulator, while strong fluorescence (normalized intensity above 0.5) was detected upon modulator binding, resulting in a prominent transition. Variants with P2 lengths ≥ 6 nt (*y*=6, 7, 8, 9, 10) displayed failed allosteric regulation, remaining in a permanent ON state due to sufficient P2 stem stability without modulator binding (no three-arm junction), resulting in strong fluorescence intensity independent of modulators. The variants without a P2 stem (*y*=0) showed ambiguous ON-OFF states, exhibiting weak fluorescence intensity even with modulators present. In conclusion, a clear ON-OFF transition was successfully achieved by designing and screening variants with pre-set P1 or P2 stems of different lengths. Given the comparable performance of P1-directed and P2-directed regulation, we selected P2-directed regulation for subsequent experiments. Similarly, the results are generally consistent with our thermodynamic simulation results (Supplementary Figs. S5-6 and Table. S2), the high value of changes of standard free energy (ΔΔG) and low switching equilibrium constant (Ks) of each successful allosteric design (*y*=1, 2, 3, 4, 5) demonstrate the distinct transition between ON and OFF state. Two successful designs (*y*= 3 and 5) are not fully consistent with the simulations results, which may be caused by the difficulties in accurately predicting the energy landscape of G-quadruplex related sequences.

For further exploring the allosteric reaction, we systematically tested six representative allosteric aptamer designs including three P1 variants (*x*=0, 1, 10) and three P2 variants (*y*=0, 2, 10). P1 (*x*=0) and P2 (*y*=0) variants represent false allosteric regulation due to insufficient ON state; P1 (*x*=1) and P2 (*y*=2) variants represent successful allosteric regulation with highest ON/OFF transition ratios (3.34× and 3.54×, respectively); P1 (*x*=10) and P2 (*y*=10) variants represent false allosteric regulation due to absence of OFF state. For P1 (*x*=0, 1) and P2 (*y*=0, 2), increasing the concentration of DFHBI-1T will not generate fluorescence without modulator, indicating the stable OFF state. After adding modulator, the fluorescence of P1 (*x*=1) and P2 (*y*=2) variants will increase with higher DFHBI-1T concentration until saturation, P1 (*x*=0) and P2 (*y*=0) variants displayed weaker or no increase in fluorescence with higher DFHBI-1T concentration. For P1 (*x*=10) and P2 (*y*=10), the fluorescence will increase with higher DFHBI-1T concentration until saturation both in the absence and presence of modulator, indicating the insufficient OFF state. Besides, P1 (*x*=1) and P2 (*y*=2) variants displayed a quick allosteric regulation speed (< 20 min) can be regulated by modulator strands with binding length (ω) over 9 nt (Supplementary Figs. S9-12).

To demonstrate the programmability of these allosteric DNA aptamer, we then incorporated toehold-mediated strand displacement into the design scheme to achieve a reversible allosteric regulation. Specifically, we designed an allosteric nano-switch where the P1 stem formed a stable 10-bp duplex, and the P2 stem formed a stable 10-bp duplex with an additional toehold region (Fig. 2F). This nano-switch features stable P1 and P2 stems with a well-defined binding pocket capable of ligand binding and strong fluorescence emission. Upon introducing a displacement strand that binds to the toehold and initiates strand displacement, the P2 stem dissociates and disrupts the binding pocket, resulting in decreased fluorescence activity (Fig. 2G, Supplementary Table. S5). Subsequent addition of an anti-displacement strand fully complementary to the displacement strand, restores the P2 stem as a stable duplex. This generates a functional binding pocket and consequently recovers fluorescence activity within 20 min (Supplementary Fig. S13).

### 2.2 Boolean logic operations based on allosteric DNA aptamer

With the successful implementation of toehold-mediated strand displacement in allosteric operations, our next goal was to use an allosteric DNA aptamer to achieve Boolean logic operations and complex nanocircuits, with the seamless incorporation of existing DNA nanotechnology design frameworks. Leveraging strand complementation and displacement, we implemented six basic logic gates (AND, OR, XOR, NAND, NOR and XNOR gates, Fig. 3). These logic gates fall into two categories: the first category (AND, OR, XOR) has an initial OFF state (output=0) when no inputs are present (input = [0, 0]). The second category (NAND, NOR, XNOR) has an initial ON state (output=1) when no inputs are present (input = [0, 0]).

**Fig. 3.**
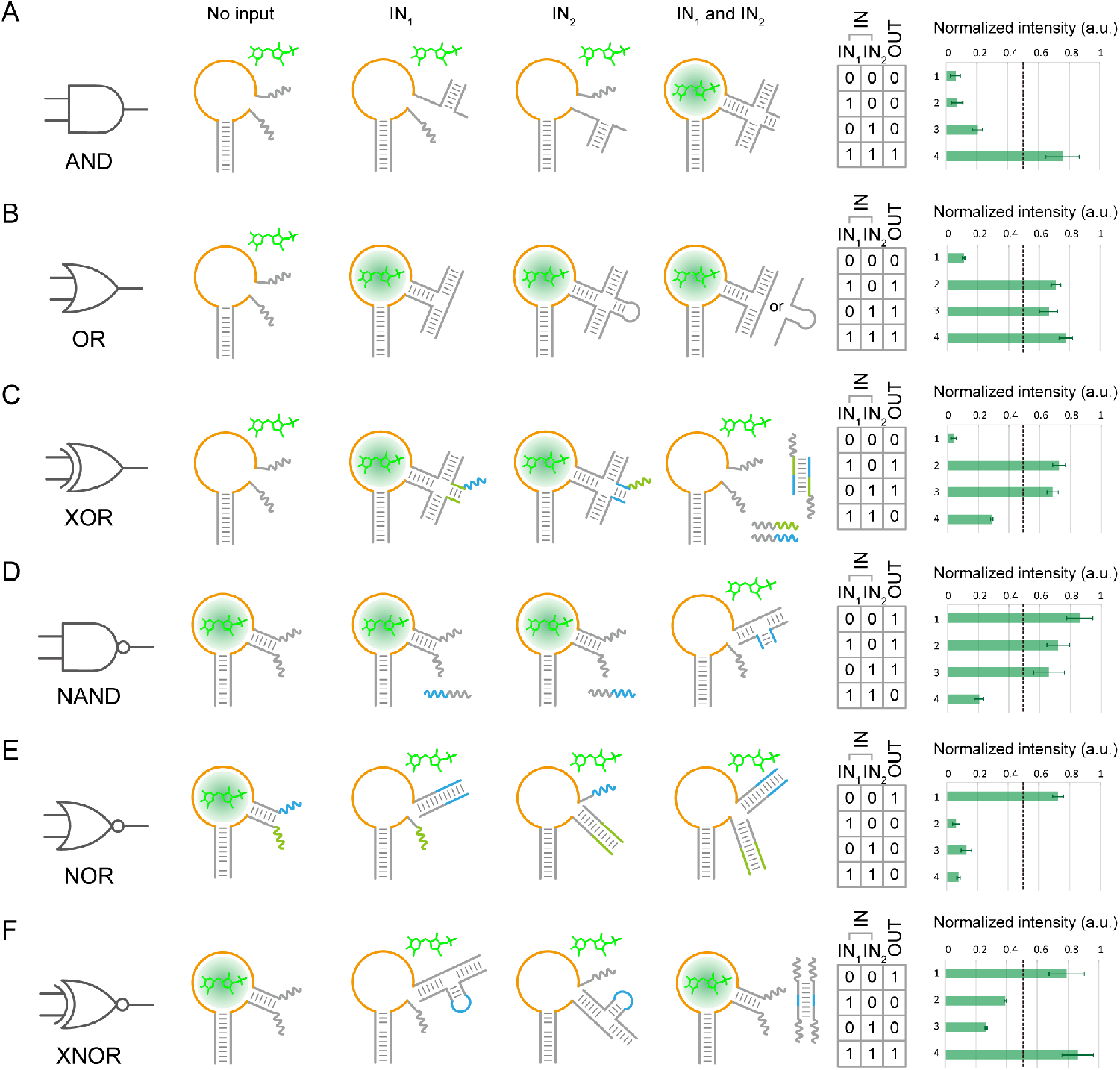
Six types of Boolean logic operations with allosteric nano-switches. **A**, AND gate. **B**, OR gate. **C**, XOR gate. **D**, NAND gate. **E**, NOR gate. **F**, XNOR gate. The logic gate was displayed in OFF or ON state without inputs, with IN1, IN2 or both inputs. Dashed lines: threshold of basic logic gates was defined as 0.5, if the normalized intensity is above 0.5, then generate a true value (output = 1); otherwise, generate a false value (output = 0). Left: design principle; right: truth table and fluorescence analysis. Errors bars, mean ± SEM.

The first category gates were constructed using an allosteric DNA aptamer design with stable P2 stems before adding modulator strands as inputs. When modulator strands were added, a conformational change stabilized the P2 stem, generating a TRUE value (output = 1). Conversely, the second category gates used an aptamer design with initially stable P2 stems. These stems became destabilized when one or both strands within the P2 stem were displaced by specific input strands. Upon addition of these displacement strands, the resulting deformation of the P2 stem and binding pocket led to fluorescence loss, generating a FALSE value (output = 0).

Taking the AND logic gate design as an example (Fig. 3A), an unstable P2 stem was configured as a 2-bp duplex segment flanked by two 15-nt unpaired single-stranded toeholds. In the absence of inputs (input = [0, 0]), the instability of P2 resulted in a deformed binding pocket, leaving the DNA aptamer in an OFF state with no fluorescent activity (output = 0). Two input strands were designed such that each bind to one toehold of the P2 stem, with the inputs themselves connected via a duplex binding region. When either input was present alone (input = [1, 0], [0, 1]), the P2 stem remained unstable, yielding no fluorescence (output = 0). Only when both inputs were available (input = [1, 1]) did a four-way junction form, stabilizing the P2 stem and creating a functional binding pocket. This switched the DNA aptamer to an ON state (output = 1). Outputs were recorded via fluorescence measurements and statistically analyzed. Similarly, OR, XOR, NAND, NOR, and XNOR gates were implemented by reconfiguring the nano-switch and strand displacement pathways (Fig. 3B-F, Supplementary Table. S6).

### 2.3 Combinatorial nanocircuits based on allosteric DNA aptamer

Building on the successful implementation of basic Boolean operations, we next aimed to construct a more complex allosteric circuit implementing a three-input combinatorial BUFFER function. We first tested the orthogonality of the allosteric regulation using 10 different pairs of nano-switches (Apt-1 ∼ Apt-10) with their corresponding modular strands (M-1 ∼ M-10). The ten modular strands had distinct sequences, ensuring each nano-switch could only be modulated by its specific modular strand (i.e., Apt-1 is modulated by M-1, Apt-2 is modulated by M-2). We measured the fluorescence intensity of each nano-switch in the presence of all ten modular strands and found no significant crosstalk (Fig. 4A, Supplementary Table. S7), demonstrating the orthogonality of this allosteric regulation strategy. We then designed a combinatorial BUFFER function with three modulators as inputs (Modulator-1 serves as IN1; Modulator-2 serves as IN2; Modulator-3 serves as IN3), as illustrated in Fig. 4B-D. In this circuit, each of the three BUFFER gates is controlled only by its target input and not by the other two inputs (IN1 for the 1st BUFFER gate; IN2 for the 2nd BUFFER gate; IN3 for the 3rd BUFFER gate). This combinatorial BUFFER function exhibited increasing fluorescence intensity levels with an increasing numbers of active inputs (Fig. 4E).

**Fig. 4.**
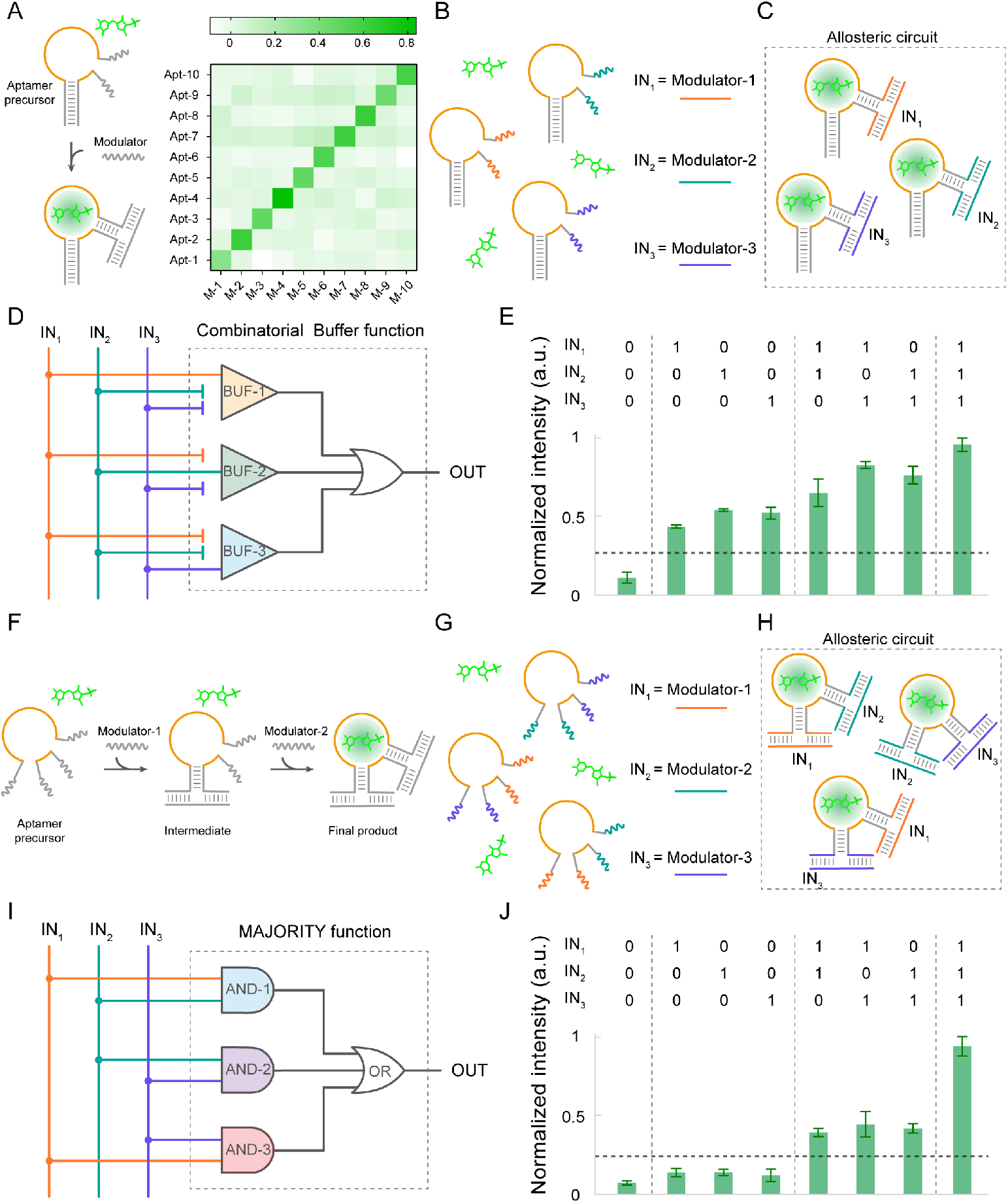
Programmable nanocircuits with allosteric nano-switches. **A**, Orthogonality test of ten different allosteric DNA aptamers with their corresponding modulator strands. **B-C**, Schematics of the combinatorial BUFFER nanocircuits, involving (**B**) three different allosteric aptamers and three different modular strands as inputs (Modulator-1 serves as IN1; Modulator-2 serves as IN2; Modulator-3 serves as IN3), and (**C**) allosteric circuit system (Aptamer-1 coupled with IN1; Aptamer-2 with IN2; Aptamer-3 with IN3). **D**, Schematics of the combinatorial BUFFER function. **E**, Statistical analysis of the allosteric nanocircuits in response to different combinations of inputs. The dash lines are set to distinguish four levels of inputs: no input, single input, double inputs and triple inputs. **F**, Schematics of the aptamers coregulated by two allosteric modulators. **G-H**, Schematics of the MAJORITY nanocircuits, involving (**G**) three different allosteric aptamers and three different modular strands as inputs, and (**H**) allosteric circuit system (Aptamer-1 coupled with IN1 and IN2; Aptamer-2 with IN2 and IN3; Aptamer-3 with IN3 and IN1). **I**, Schematics of the combinatorial MAJORITY function. **J**, Statistical analysis of the allosteric nanocircuits in response to different combinations of inputs. The dash lines are set to distinguish four levels of inputs: no input, single input, double inputs and triple inputs. Dashed lines: threshold of functions was defined as 0.25, if the normalized intensity is above 0.25, then generate a true value (output = 1); otherwise, generate a false value (output = 0). Errors bars, mean ± SEM.

We then constructed another complex allosteric circuit implementing the three-input MAJORITY function—a combinational circuit that generates TRUE value (or 1) when the majority of inputs (≥ 2) are added, otherwise the outputs are FALSE (or 0). To implement this, we integrated three orthogonal AND gates and one OR gate. Each AND gate was constructed using a co-allosteric regulation mechanism in which both the P1 and P2 stems of a nano-switch are controlled by two different modular strands. Each AND logic gate (representing one nano-switch) is regulated by two inputs, with every pair of gates sharing one common input, as illustrated in Fig. 4F-I. Specifically: INA and INB regulate the first AND gate; INB and INC regulate the second AND gate; INC and INA regulate the third AND gate. As expected, no fluorescence was detected in the absence of any inputs. When a majority of inputs (2 or 3 inputs) were present, most nano-switches underwent conformational changes, producing green fluorescence. Conversely, the system maintained its original conformation with any single input, and no fluorescence was detected (Fig. 4F-J, Supplementary Table. S8).

With an immense sequence space defined by a four-letter code (4^N^, where N is the length of modulator strand), we envision that many more distinct species of nano-switches could be created to implement advanced algorithms and molecular computing tasks.

### 2.4 Light-up DNA nanostructures with allosteric DNA aptamer

The programmable and controllable fluorescence properties prompt us to apply our allosteric aptamers as a reporter system for the fluorescent imaging of DNA nanostructures. Conventional fluorescent imaging of DNA nanostructures relies on chemically modified fluorophore-conjugated probes, where fluorescence intensity depends solely on fluorophore properties and is therefore non-adjustable. For example, the collective intensity of a nanostructure target associated with a probe or probe complex is independent of the probe configurations and clustering status. In contrast, our allosteric aptamer produces fluorescent ON/OFF signals in direct response to configurational transition, with the structural properties of DNA nanostructures serving as the foundation for its fluorescent functionality.

To validate this approach, we used a 10-helix bundle (10-HB) DNA origami as a representative DNA nanostructure and evaluated its fluorescent performance when modified with allosteric DNA aptamers. We incorporated allosteric DNA aptamers into 16 pairs of staples along a constituent helix (Helix-9) of the 10-HB origami (Supplementary Fig. S14 and S19), creating a modified design—10-HB-apt. In a well-folded state where all staples bind to scaffolds as designed, the allosteric aptamer adopts a proper binding pocket configuration, enabling the nanostructure to exhibit high fluorescent intensity. When the morphological state of nanostructure is compromised (e.g., due to degradation or denaturation), misfolded staples disrupt the aptamer binding pocket, resulting in fluorescence loss. For comparison, we appended the same 16 modification sites with FAM-labeled DNA probes in a 10-HB-pro design as a control group. To test whether the structural integrity of 10-HB affects fluorescence, we measured the fluorescent intensity of both designs after treatment with DNase I-induced degradation and heat-induced denaturation, respectively (Supplementary Figs. S17-S18).

DNase I is an endonuclease that degrades both single- and double-stranded DNA into dinucleotide, trinucleotide, and oligonucleotide products^[36-37]^, which can significantly compromise the structural integrity of DNA nanostructures^[38-39]^. Agarose gel electrophoresis revealed dissociated structures after degradation with DNase I (at a final concentration of 20 Unit/mL). Consistently, fluorescence measurement showed signal loss in 10-HB-apt after DNase I treatment, while the signal remained constant for the 10-HB-pro control despite structural decay. Similarly, heating (40-80°C) produced analogous structure-dependent effects. The fluorescence signal of 10-HB-apt was significantly more sensitive to denaturation by heating than that of 10-HB-pro (Supplementary Table. S9). These degradation and denaturation tests demonstrated that the fluorescent functionality of allosteric DNA nanostructures depends on proper structure configuration.

Next, we implemented our allosteric strategy to achieve inducible lighting-up of DNA nanostructures. To achieve higher fluorescence intensity under fluorescence microscopy, we incorporated allosteric DNA aptamers into 32 pairs of staples within two constituent helices of the 10-HB origami (16 pairs along Helix-9 and 16 pairs along Helix-0, Fig. 5A and Supplementary Fig. S14). All 32 sites contain the same aptamer regulatory sequence, which binds to identical modulator oligos. In the absence of the modulator, the allosteric aptamer lacks a stable binding pocket and emits no fluorescence. After introducing the modulator into the system, its binding to the two tails of the P2 stem induced the aptamer binding pocket into an ON state, triggering fluorescence emission from the nanostructures.

**Fig. 5.**
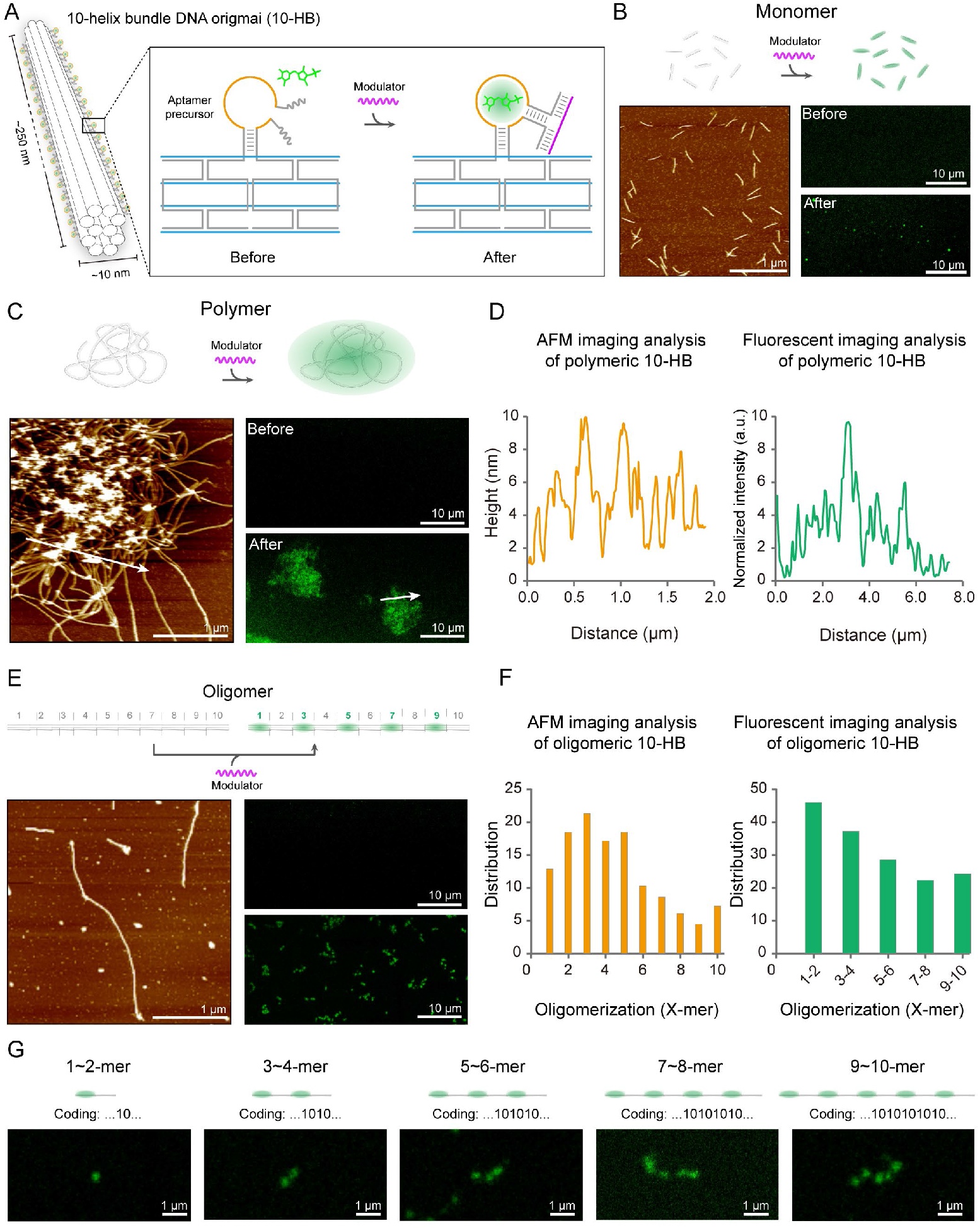
Light-up nanostructures with allosteric DNA aptamer. **A**, Schematics of the 10-HB origami appended with allosteric aptamers at 32 sites along two helices. **B, C, E**, Schematics (top) depicting the inducible lighting-up of monomeric (**B**), polymeric (**C**) and oligomeric (**E**) 10-HB origami. Bottom-left: AFM imaging results; Bottom-right: CLSM imaging results. Scale bars for AFM images: 1 μm; scale bars for CLSM images: 10 μm. **D, F**, AFM imaging analysis (left) and fluorescent imaging analysis of polymeric (**D**) and oligomeric (**F**) 10-HB origami. For statistical analysis in (**F**), N is ∼200. **G**, Schematics and CLSM imaging results of representative 10-HB origami oligomers with different binary codes. Scale bars: 1 μm.

The 10-HB origami was engineered to adopt three distinct clustering states—monomer, polymer, and oligomer—by programming intermolecular edge-edge interactions using complementary sticky ends between neighboring origami units (Supplementary Figs. S15-S16). Monomeric 10-HB origami displayed dispersed rod-like structures under atomic force microscopy imaging (AFM) and consistently appeared as dispersed bright dots (considering the diffraction limit in optics is 200 nm, the 10-HB origami with a similar length (∼250 nm) can only present a round spots under optical microscopy) under confocal fluorescence microscopy (CLSM) after allosteric regulation by modulators (Fig. 5B, Supplementary Figs. S20-S21). Polymeric 10-HB origami formed micrometer-scale clusters observable under both AFM and CLSM (Fig. 5C, Supplementary Figs. S22-S23), with similar density distributions (height distribution under AFM and intensity distribution under CLSM). For the oligomeric design, ten variants of the 10-HB origami monomers (O1∼O10) were designed. These origami shared the same structural core but featured ten distinct groups of connection edge staples, enabling specific end-to-end assembly into long linear structures. To generate a fluorescent dotted-line pattern, we modified O1/O3/O5/O7/O9 with allosteric aptamer which can generate fluorescence while leaving O2/O4/O6/O8/O10 unmodified without fluorescence generation. As a result, the oligomer will present an intermittent fluorescent dot pattern (…101010…, “1” means fluorescence dots and “0” means darkness) under fluorescence microscopy (Fig. 5G). During a one-pot assembly of these 10-HB variants, we observed a distribution of oligomer lengths (Fig. 5E, Supplementary Figs. S24-S25), indicating varying degrees of oligomerization. The consistent structural distributions revealed by both AFM and CLSM validate the feasibility of using allosteric aptamer to characterize DNA nanostructures.

After the successfully demonstration of inducible lighting-up of DNA nanostructures using allosteric aptamers, we next proceeded to use mRNAs as modulators to light up DNA nanostructures. In this part, 16 pairs of staples along Helix-9 of the 10-HB origami (Fig. 6A) were modified with allosteric aptamers targeting 16 distinct binding sequences. These aptamers on a single 10-HB origami were designed to recognize and bind in tandem to 16 target sites (20 nt per site) on a specific mRNA molecule, enabling 1:1 target recognition. We tested the performance of this allosteric aptamer-functionalized DNA origami using three in vitro transcribed mRNA targets (designated mRNA-1, mRNA-2, and mRNA-3). The functionalized 10-HB origami initially presented no obvious fluorescence signal in the absence of target mRNA and generated a positive signal upon target binding, though the fluorescent intensity remained limited (Fig. 6B, Supplementary Fig. S25A-B). We hypothesized that this limited intensity resulted from intrinsic mRNA secondary structures compromising the accessibility of target mRNA sites for allosteric aptamers. To address this, we treated mRNA targets with a denaturing condition (heating at 80°C followed by rapid cooling on ice) to disrupt secondary structures and minimize refolding before hybridization with the allosteric aptamers. Consequently, denatured mRNA-1 and mRNA-2 targets exhibited significant fluorescence enhancement (Fig. 6C). Moreover, mRNA-1 induced higher intensity than mRNA-2 did, likely due to its weaker secondary structure.

**Fig. 6.**
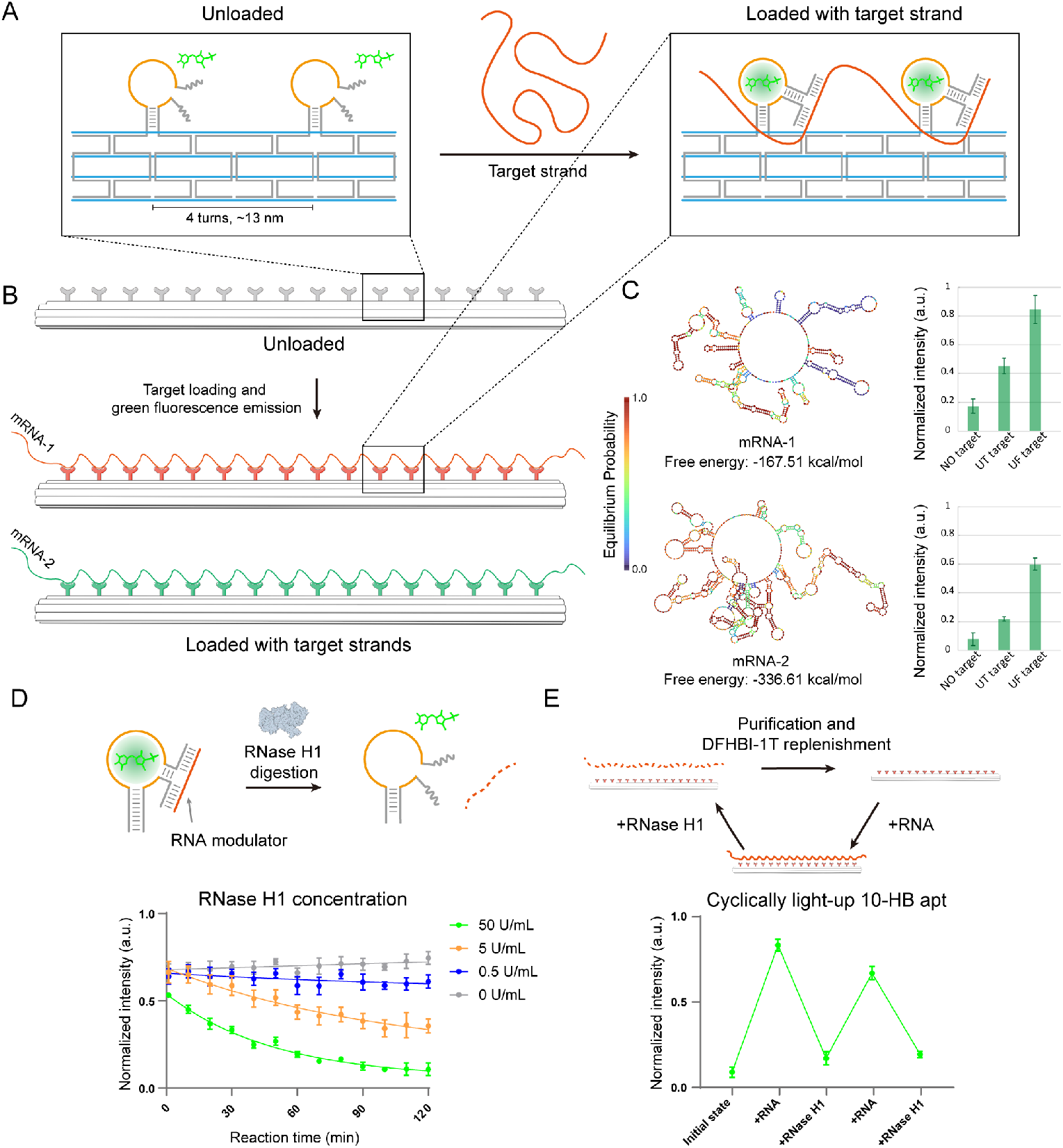
Light-up DNA nanostructures with RNA. **A-B**, Schematics of the 10-HB origami appended with allosteric aptamers at 16 sites along one helix that can recognize and bind to target mRNA. **C**, Left, schematics of the secondary and energy landscape of mRNA-1 and mRNA-2. Right, statistical analysis of fluorescence intensity of 10-HB origami before and after binding to target mRNA. UT target, untreated mRNA target; UF, unfolded mRNA targets after heating and crash-cooling on ice. **D**, Schematics and digestion results using different concentration of RNase H1. **E**, Schematics and fluorescence analysis showing two cycles of 10-HB origami light-up. Errors bars, mean ± SEM.

However, the binding of denatured mRNA-3 did not yield a significant enhancement in fluorescence intensity, despite its likely minimal secondary structure (Supplementary Fig. S25C). Given that mRNA-3 is only ∼500-nt long, a tightly stretched configuration with 10-nt linkers between binding segments is necessary for it to fully bind all 16 aptamers on the 10-HB origami. Consequently, skipping certain binding segments was deemed inevitable, resulting in reduced fluorescent intensity for mRNA-3. Increasing the concentration ratio of mRNA to 10-HB origami counteracted the effect of omitted aptamer binding, leading to a compensatory increase in fluorescence intensity (Supplementary Fig. S25D and Table. S10). Therefore, designing an appropriate aptamer density to match a specific mRNA length is preferable.

After demonstrating the possibility of using RNA to light-up DNA nanostructures, we took advantage of the degradable nature of RNA and proceed to build a cyclical nanostructure light-up platform under the assistance of RNase H1, which can degrade RNA strands from DNA-RNA hybrids. Through a screening on a simple allosteric aptamer, we demonstrated RNase H1 can fully degrade all RNA modulators at a minimum concentration of 50 U/mL (Fig. 6D). For cyclically lighting-up DNA nanostructures, mRNA-1 was first used to light up 10-HB origami equipped with allosteric aptamers. The 10-HB origami was then quenched due to mRNA-1 degradation via RNase H1 treatment in order to finish the first cycle. Following the first cycle, the 10-HB origami was purified with centrifugal filter to remove RNase H1 and then replenished with DFHBI-1T to the original concentration. The second cycle was finished with the same mRNA-1 binding, RNase H1 digestion and purification procedure. Fluorescence analysis showed successful implementation of such a cyclical nanostructure light-up platform (Fig. 6E and Supplementary Table. S11). We believe our system will provide a very efficient and cost-friendly method to retrieve target nanostructures in the DNA nanostructure-based information repository, which can be used for repetitive information reading and writing in the future.

## 3. Conclusion

Building on programmable allosteric regulation, we created a series of synthetic nucleic acid nano-switches with precisely controlled fluorescence ON-OFF transition. By employing a single-stranded DNA modulator targeting the allosteric site, we enabled adjustable fluorescent readouts in these custom-designed DNA nano-switches. This robust system allows for diverse allosteric regulatory functions, including designated allosteric transitions, reverse allosteric regulation, and complex logic operations.

The study here offers a highly adaptable design framework for nucleic acid nanotechnology, highlighting its extensibility and vast application potential. By modular integration of diverse functional DNA sequences responsive to environmental cues (expanded allosteric modulators, including proteins, ions, pH, light, etc.), the core allosteric switching mechanism could become a universal platform. Such a universal platform can allow researchers to “plug in” specific sensors tailored to desired biological, chemical and physical signals, transforming allosteric aptamers into versatile fluorescent detectors or actuators, pointing directly toward sophisticated applications in bioimaging and diagnostics. Beyond the fluorescence-based readouts demonstrated in this study, the framework is readily adaptable for allosterically regulating a wide range of biochemical activities by substituting the fluorescent aptamer core with other functional DNAs or RNAs. This flexibility opens new avenues for engineering modular riboswitches, ribozymes, and RNA logic circuits with customized responses.

By synthetically building these allosteric devices from scratch, the platform also offers an approach for researchers to investigate how structure dictates function and dynamics in nucleic acids and other biomolecular systems. This synthetic biology approach provides a simplified system to dissect allostery principles that might be obscured in more complex natural systems. Attempting to rationally engineer or re-engineer natural biomolecules (proteins, RNAs) or complexes using similar principles can lead to deeper mechanistic understanding of their function and uncover previously unrealized application potential in both biology (e.g., designing novel enzymes, metabolic pathways) and material science (e.g., creating programmable biomaterials, self-assembling nanostructures).

In sum, the study here offers an extensible synthetic framework that enables the creation of sophisticated DNA devices for potential applications, while simultaneously providing fundamental insights and design principles. These principles, in turn, empower the rational engineering of both synthetic and naturally occurring systems, advancing both basic scientific understanding and technological innovation across disciplines.

## 4. Experimental Section

### Oligonucleotides

DNA sequences were designed by Uniquimer^[40]^ and tested by NUPACK^[41]^ to prevent unspecific complementation. All unlabeled DNA strands (with PAGE purification) were purchased from Sangon and GENEWIZ. Fluorophore labeled DNA strands (with HPLC purification) were purchased from Sangon Biotech Co. Ltd. Details about DNA sequences were available in **Sequences** part in Supplementary Information.

### dsDNA templates

In vitro transcription DNA templates were purchased from GENEWIZ and amplified in 50 μL PCR mix with 1× PrimeSTAR® Max DNA Polymerase Premix (Takara) and two terminal primers. The PCR amplification program consisted of 35 cycles: 98°C for 10 s, 57°C for 5 s and 72°C for ∼ 10−20 s (10 s/kb). PCR products were analyzed with 1% agarose gel electrophoresis and then extracted with a gel purification kit (Mei5bio). Detailed DNA template sequences were available in **Sequences** part in Supplementary Information.

### Chemicals

Reagent-grade chemicals (Tris base, Acetic acid, Magnesium chloride, EDTA, Boric acid, Sodium chloride, Potassium chloride, SYBR gold, DFHBI-1T, etc.) commercially available were used without further purification.

#### Buffers

1. Allosteric aptamer assembly buffer: **Buffer 1** (10×): 400 mM tris base, 200 mM acetic acid, 10 mM EDTA and 125 mM magnesium chloride, 1500 mM Potassium chloride, pH ∼8.0;
2. DNA origami assembly buffer: **Buffer 2** (10×): 400 mM tris base, 200 mM acetic acid, 10 mM EDTA and 125 mM magnesium chloride, pH ∼8.0;
3. AFM scanning buffer: **Buffer 3** (1×): 20 mM tris base, 0.5 mM EDTA, 10 mM magnesium chloride, pH ∼8.0;
4. DFHBI-1T staining buffer: **Buffer 4** (1×): 40 mM tris base, 20 mM acetic acid, 150 mM potassium chloride, 12.5 mM magnesium chloride, pH ∼8.0;

### Preparation of allosteric aptamer

For all DNA aptamers, final concentration of each DNA strand in a 60 μL mixture solution (10×**Buffer 1** diluted to 1×) was 500 nM unless otherwise specified. The DNA mixture solutions were then thermally annealed: 95 °C for 2 min, followed by a gradient from 85 °C to 22°C at a rate of 1°C/30 s, then hold at 22 °C.

### ON-OFF regulation and logic operation

DNA modulators or inputs were then added into the annealed solution with their stoichiometric molar ratio (1:1 in ON-OFF tests/logic gates operations/combinatorial Buffer function, 2:1 in Majority function) and incubated overnight at 22 °C.

### NUPACK simulation

Thermodynamic simulation was performed in NUPACK with each sequence and their target secondary structure (Dot-Bracket notation) as inputs. All simulations were carried out under the following conditions: DNA concentration is 1 μM, temperature is 25 °C, monovalent cation concentration is 0.15 M, bivalent cation concentration is 0.0125 M.

### DNA origami monomer/polymer preparation

For the assembly of 10-HB DNA origami monomers and polymers, 10 nM p8064 ssDNA scaffold and 50 nM staples (including core staple strands and edge staple strands) were mixed in 1× **Buffer 2** solution and annealed using the following protocol: 85 °C for 5 min followed by a gradient from 85 to 25 °C by −0.5 °C/1 min and 25 to 4 °C by −3 °C/1 min. Transfer the assembly product into a 50 kDa Amicon Ultra Centrifugal Filter (Millipore), add 400 μL 1×**Buffer 2** and centrifuge at 13,000×g for 2 min. Discard the centrifugate and repeat for 4 times to remove excess staples. We supposed there was no origami loss during the filtration, thus the final concentration of 10-HB origami can be calculated according to the total amount of origami in initial solution and the volume of final filter residue. Modular strands or FAM probes were then added into the filter residue with a stoichiometric molar ratio (16:1 or 32:1) and incubated overnight at 22 °C.

### DNA origami oligomer preparation

For the assembly of 10-HB DNA origami oligomers, each of the 10 components was first assembled independently with their respective set of edge staples. Then transfer all components into a 50 kDa Amicon Ultra Centrifugal Filter (Millipore) for a one-pot purification, add 450 μL 1×**Buffer 2** and centrifuge at 13,000×g for 2 min. Discard the centrifugate and repeat for 4 times to remove excess staples. The purified product was incubated at 42 °C for 8 hours to form oligomer. Modular strands were then added with a molar ratio of 32:1 and incubated overnight at 22 °C.

### DNase I-induced degradation

0.5 or 1 μL DNase I enzyme (Sangon) was added into 50 μL of pre-incubated 10-HB-apt or 10-HB-pro origami (10 nM) solutions, and incubated at 37 °C for 10 min. The product was quickly loaded into agarose gels for agarose gel electrophoresis.

### Heating-induced denaturation

50 μL of pre-incubated 10-HB-apt or 10-HB-pro origami (10 nM) solutions were heated at 40∼80 °C for 5 min and followed by an incubation at 22 °C for 20 min.

### Native agarose gel electrophoresis (AGE)

1% native agarose gel was prepared with 0.5×TBE (10 mM Mg^2+^) as the gel running buffer. Gels were run at 90 V (constant voltage, Bio-Rad) for 120 min in an ice-water bath. After electrophoresis, gel was imaged in Amersham ImageQuant 800 (Cytiva) at designative channel.

### In vitro transcription

dsDNA templates were transcribed with HiScribe™ T7 Quick High Yield RNA Synthesis Kit (NEB) following the manufacturer’s instructions. The transcription products were purified with DEPC-treated water (Sangon) in a 30 kDa Amicon Ultra Centrifugal Filter (Millipore) for 4 times to remove excess rNTPs and ions.

### Light-up 10-HB origami with RNA

RNA was thermally denatured with the following protocol: heating at 80°C for 3 min followed by crash-cooling on ice for 5 min. Denatured RNA were quickly mixed with pre-annealed 10-HB origami with a molar ratio of 1:1 at 4 °C overnight. To prevent RNA degradation, Rnase inhibitor (Sangon) was also added into the solution following the manufacturer’s instructions. For cyclically lighting-up DNA nanostructure, RNase H1 (Beyotime Biotech) was added to a final concentration of 100 U/mL and incubated for 1 h. The digestion product was the purified with 50 kDa Amicon Ultra Centrifugal Filter (Millipore) to remove RNase H1. The purified 10-HB origami was then quantified by Nanodrop. Because DFHBI-1T will also lost during the purification, new DFHBI-1T was then added to the original concentration.

### Fluorescence intensity measurement

Fluorescence measurements for DFHBI-1T (MedChemExpress) binding were performed in the following protocol: aptamer-DFHBI-1T complexes were prepared by incubation of annealed DNA nanodevices or origami in 1×**Buffer 4** at a stoichiometric molar ratio (aptamer: DFHBI-1T = 1:2) at 22 °C for 30 min. Fluorescence intensity was recorded in 96-well plate using a Varioskan Flash Spectral Scanning Multimode Reader (Thermo Fisher Scientific) or Spark High Performance Multimode Plate Reader (Tecan) at excitation and emission wavelengths of 465 nm and 507 nm, respectively. Fluorescence intensity of FAM probe-labeled origami was recorded at excitation and emission wavelengths of 490 nm and 520 nm, respectively.

Fluorescence intensity values were normalized using the following equation:

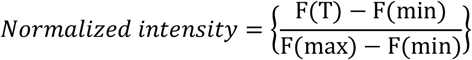

where F(T) is the fluorescence signal of target allosteric aptamers obtained under fluorescence reader, F(max) is the designated maximum fluorescence signal, F(min) is the designated minimum fluorescence signal which is also the background signal in the absence of DNA. Binding curves were obtained using the following equation:

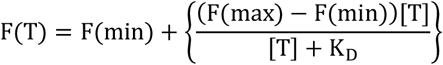

where [T] is the concentration of DFHBI-1T, K_D_ is the dissociation constant.

### AFM imaging

After annealing, 2 μL of diluted origami sample (1∼5 nM) was loaded onto freshly cleaved mica. After incubation for ∼1 min, 40 μL of **Buffer 3** were reloaded onto mica. AFM images were obtained using a Multimode-8 microscope with Nanoscope V controller (Bruker). Samples were imaged under liquid ScanAsyst mode with C-type triangle tips (resonant frequency, f0 = 40∼75 kHz; spring constant, k = 0.24 N m^-1^) from the SNL-10 silicon nitride cantilever chip (Bruker).

### Confocal laser-scanning microscopy (CLSM) imaging

5 μL of diluted origami sample (0.1 nM) was loaded to the microscope slide and the coverslip was placed on top of the sample by applying gentle pressure. Samples were imaged using a confocal laser scanning microscope (Nikon A1 HD25) equipped with a 488 nm laser. Images were analyzed with the same adjustment to substrate background fluorescence using Nikon Elements Analysis software.

## Supporting Information

Supporting Information is available from the author.

## Acknowledgements

This work is supported by: National Key R&D Program of China (grant No. 2021YFF1200200), National Natural Science Foundation of China (grant No. 31770926,), Tsinghua University Spring Breeze Fund, and Tsinghua University-Peking University Joint Center for Life Sciences (to B.W.); National Key R&D Program of China (grant No. 2023YFC2306300), National Natural Science Foundation of China (grant No. 32225019, 92357304, 32394003), Beijing Natural Science Foundation (grant No. 5222010), Tsinghua University-Peking University Joint Center for Life Sciences, the Institute for Immunology, and School of Basic Medical Sciences at Tsinghua University (to W.Z.); National Natural Science Foundation of China (grant No. 22225402, 32341017), Leading Innovative and Entrepreneur Team Introduction Program of Zhejiang Province (grant No. 2024R01005), Hangzhou Institute of Medicine, Chinese Academy of Sciences (to D.H.); Research Grant Council of Hong Kong (grant No. 14207421), Department of Biomedical Engineering, Chinese University of Hong Kong (to Z.G.).

## Data availability

All the data presented in the article and Supplementary Information are available from the corresponding authors upon reasonable request.

## Author contributions

T.Z. and X.Q. conceived the study and conducted the experiments. J.Z. and H.C. assisted with the experiment design. T.B. and W.W. designed the 10HB origami. Y.W. assisted with sample preparation. T.Z., B.W., W.Z., D.H., Z.G. supervised the project. All authors analyzed the data and wrote the manuscript.

