## supplemental information for "Light-up nanostructures with allosteric DNA mimics of GFP"

### This PDF file includes:

Supplementary Figures S1 to S25

Supplementary Tables S1 to S11

Sequences

#### Supplementary Figures

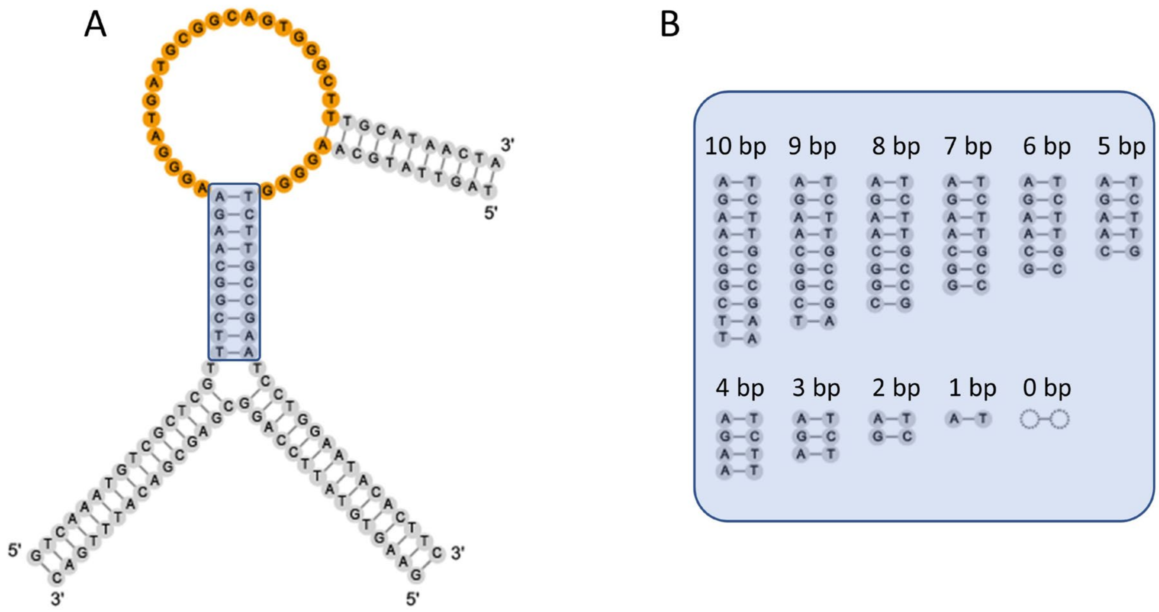

**Figure S1.** Design schematics of the allosteric nano-switch regulated by P1 formation. **A**, strand diagram of the nano-switch with a 10-bp P1 stem. **B**, strand diagram depicting varying P1 lengths (*x*= 0~10).

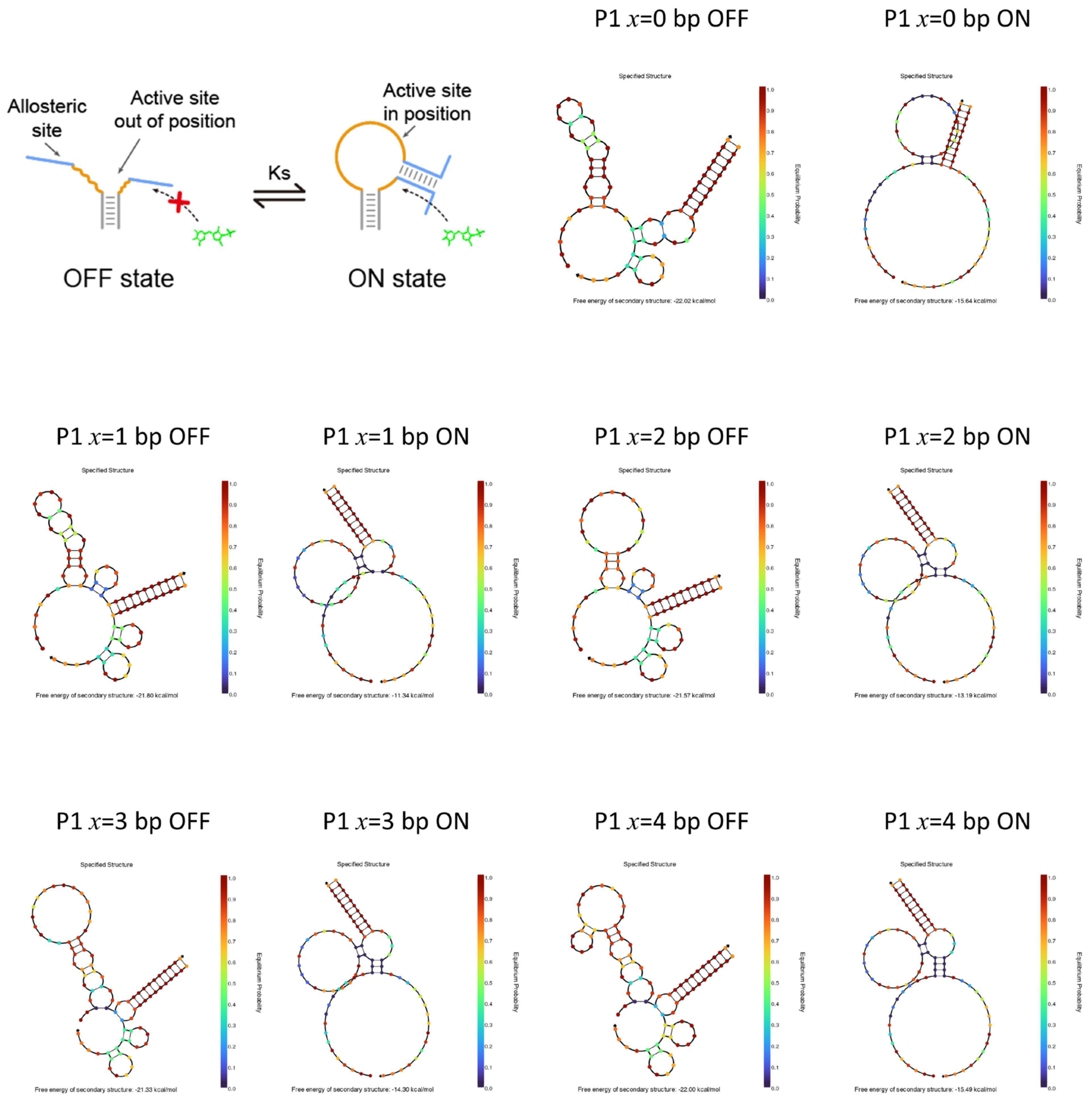

**Figure S2.** NUPACK[1, 2] simulation of the minimum free energy (MFE) of nano-switches of varying P1 lengths ($x$=1~4) in the ON and OFF states.

It's worth noting that there are still no reliable tools to accurately predict and simulate G-quadruplex structures which is exactly the core structure in Lettuce aptamer for DFHBI-1T binding and fluorescence generation. Thus, in the NUPACK platform, we neglect the thermodynamic parameters of G-quadruplex structures and used the default simulation results for further calculation.

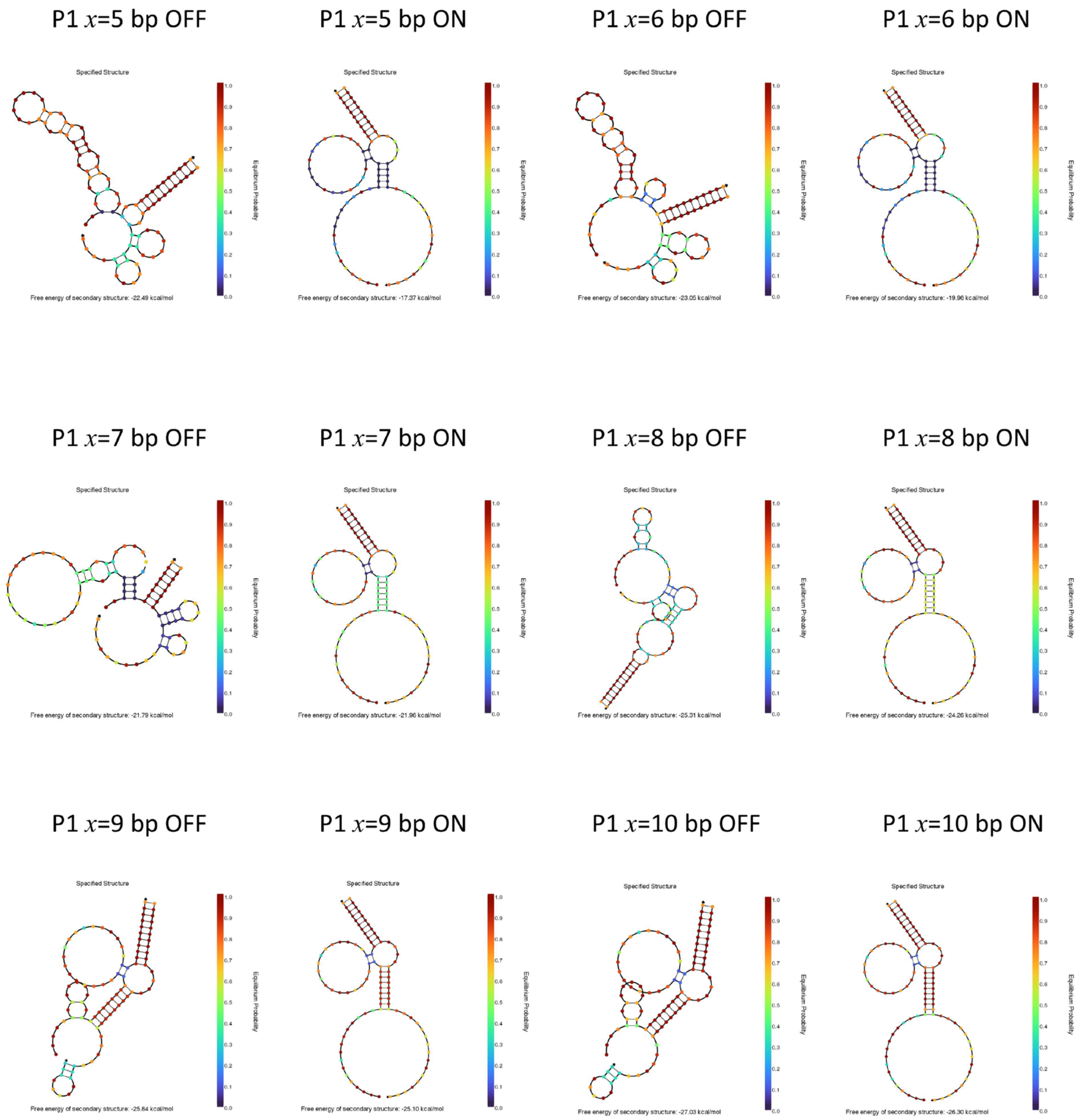

**Figure S3.** NUPACK simulation of the minimum free energy (MFE) of nano-switches of varying P1 lengths (*x*=5~10) in the ON and OFF states.

P1 $x$= 0~10bp

| | 0 bp | 1 bp | 2 bp | 3 bp | 4 bp | 5 bp | 6 bp | 7 bp | 8 bp | 9 bp | 10 bp |
|---|---|---|---|---|---|---|---|---|---|---|---|
| kcal/mol $\Delta G_{OFF\text{-}state}$ | -22.02 | -21.8 | -21.57 | -21.33 | -22 | -22.49 | -23.05 | -21.79 | -25.31 | -25.84 | -27.03 |
| kcal/mol $\Delta G_{ON\text{-}state}$ | -15.64 | -11.34 | -13.19 | -14.3 | -15.49 | -17.37 | -19.96 | -21.96 | -24.26 | -25.1 | -26.3 |
| $\Delta\Delta G$ | 6.38 | 10.46 | 8.38 | 7.03 | 6.51 | 5.12 | 3.09 | -0.17 | 1.05 | 0.74 | 0.73 |
| $K_S$ | 1.9E-05 | 1.8E-08 | 6.2E-07 | 6.2E-06 | 1.5E-05 | 1.6E-04 | 5.1E-03 | 1.3E+00 | 1.7E-01 | 2.8E-01 | 2.9E-01 |

**Table S1.** The Gibbs free energies and switching equilibrium constant Ks of each aptamer calculated from NUPACK simulation results.

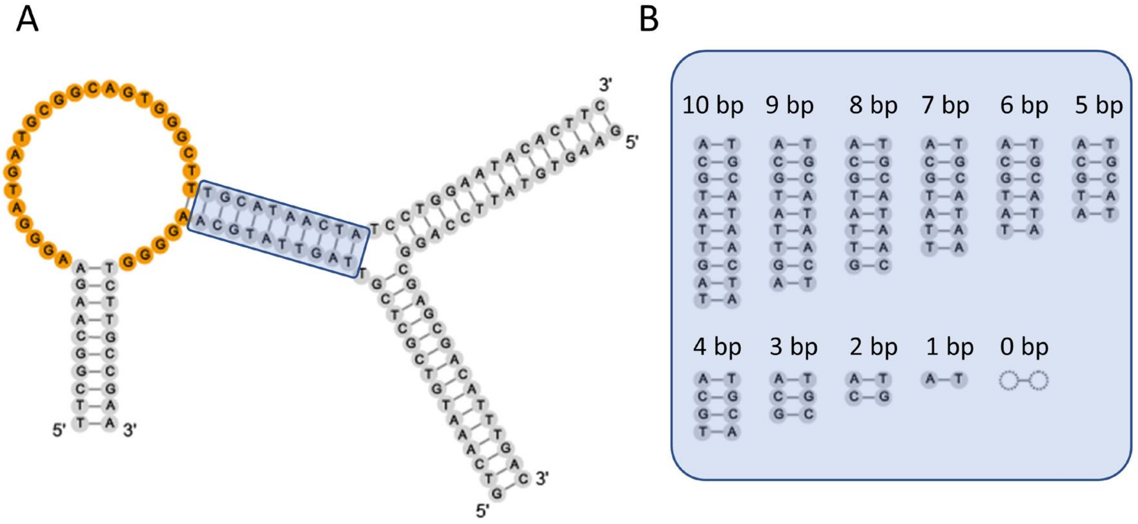

**Figure S4.** Design schematics of the allosteric aptamer regulated by P2 formation. **A**, strand diagram of the nano-switch with a 10-bp P2 stem. **B**, strand diagram depicting varying P2 lengths (*y*= 0~10).

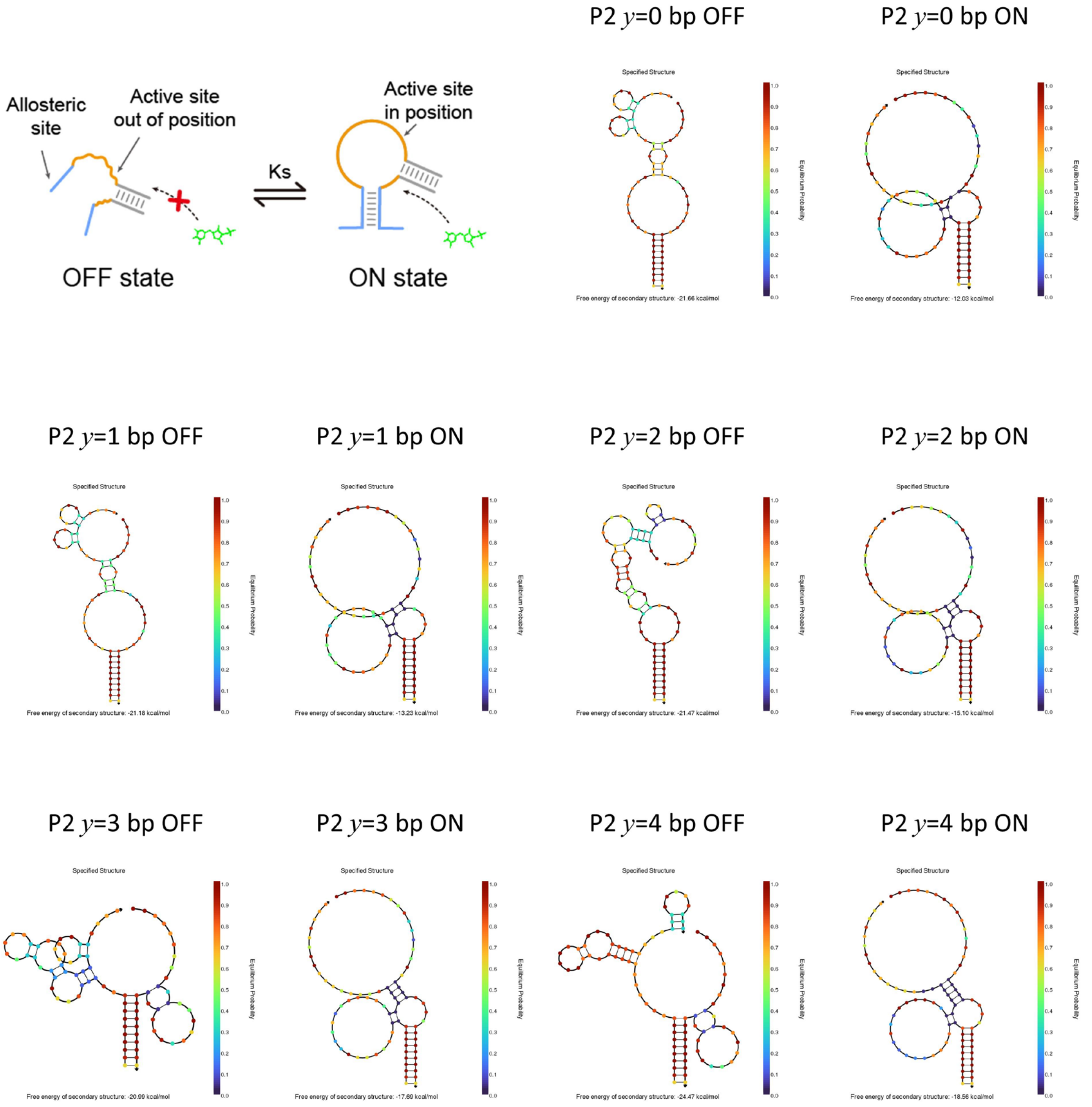

**Figure S5.** Simulated minimum free energy (MFE) of allosteric aptamer of varying P2 lengths (*y*=1~4) in the ON and OFF states.

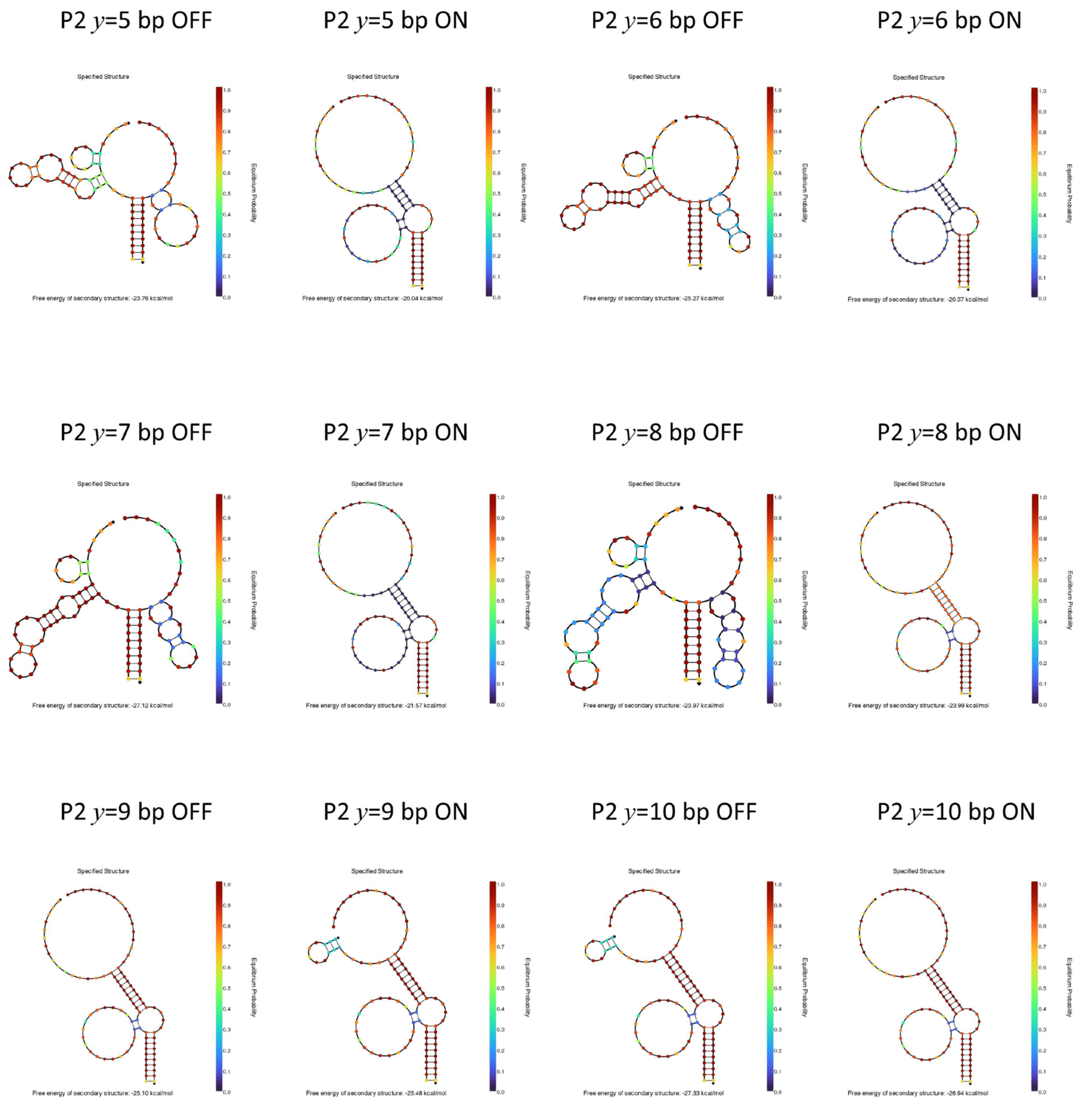

**Figure S6.** Simulated minimum free energy (MFE) of allosteric aptamer of varying P2 lengths ($y$=5~10) in the ON and OFF states.

P2 $y$= 0~10bp

| | | 0 bp | 1 bp | 2 bp | 3 bp | 4 bp | 5 bp | 6 bp | 7 bp | 8 bp | 9 bp | 10 bp |
|---|---|---|---|---|---|---|---|---|---|---|---|---|
| kcal/mol | $\Delta G_{OFF\text{-}state}$ | -21.66 | -21.18 | -21.47 | -20.99 | -24.47 | -23.76 | -25.27 | -27.12 | -23.97 | -25.48 | -27.33 |
| | $\Delta G_{ON\text{-}state}$ | -12.03 | -13.23 | -15.1 | -17.69 | -18.56 | -20.04 | -20.37 | -21.57 | -23.99 | -25.1 | -26.94 |
| | $\Delta\Delta G$ | 9.63 | 7.95 | 6.37 | 3.3 | 5.91 | 3.72 | 4.9 | 5.55 | -0.02 | 0.38 | 0.39 |
| | $K_S$ | 7.3E-08 | 1.3E-06 | 1.9E-05 | 3.6E-03 | 4.2E-05 | 1.8E-03 | 2.3E-04 | 7.7E-05 | 1.0E+00 | 5.2E-01 | 5.1E-01 |

**Table S2.** The Gibbs free energies and switching equilibrium constant Ks of each aptamer calculated from NUPACK simulation results.

Normalized intensity of P1 variants (3 replicates)

| X (bp) | 0 bp | 1 bp | 2 bp | 3 bp | 4 bp | 5 bp | 6 bp | 7 bp | 8 bp | 9 bp | 10 bp |
|---|---|---|---|---|---|---|---|---|---|---|---|
| With Modular | 0.3852 | 0.8082 | 0.5917 | 0.8138 | 0.4818 | 0.7544 | 0.5073 | 0.7610 | 0.7888 | 0.6793 | 0.7944 |
| | 0.4591 | 0.7066 | 0.7484 | 0.8239 | 0.7385 | 0.9337 | 0.7323 | 0.7773 | 0.8247 | 0.5525 | 0.7239 |
| | 0.4165 | 0.5636 | 0.6249 | 0.6221 | 0.6949 | 0.7231 | 0.5626 | 0.6806 | 0.9132 | 0.7896 | 0.8433 |
| Without Modular | 0.0216 | 0.1269 | 0.1454 | 0.2287 | 0.4188 | 0.6196 | 0.6855 | 0.5812 | 0.6130 | 0.6830 | 0.6269 |
| | 0.0479 | 0.2790 | 0.2786 | 0.1812 | 0.3697 | 0.4645 | 0.5630 | 0.6137 | 0.5365 | 0.7146 | 0.7435 |
| | 0.1048 | 0.2164 | 0.2146 | 0.2789 | 0.2956 | 0.5862 | 0.6753 | 0.6797 | 0.6812 | 0.6279 | 0.6521 |

**Table S3.** Normalized fluorescence intensity of allosteric aptamer with varying P1 lengths.

Normalized intensity of P2 variants (3 replicates)

| y (bp) | 0 bp | 1 bp | 2 bp | 3 bp | 4 bp | 5 bp | 6 bp | 7 bp | 8 bp | 9 bp | 10 bp |
|---|---|---|---|---|---|---|---|---|---|---|---|
| With Modular | -0.0664 | 0.5889 | 0.8969 | 0.6609 | 0.7537 | 0.7661 | 0.7482 | 0.8037 | 0.7959 | 0.7742 | 0.6987 |
| | 0.1060 | 0.4580 | 0.7769 | 0.5765 | 0.6236 | 0.7116 | 0.5933 | 0.8376 | 0.6111 | 0.7717 | 0.6573 |
| | 0.2065 | 0.6033 | 0.6943 | 0.7170 | 0.7021 | 0.7280 | 0.6993 | 0.9187 | 0.6647 | 0.8461 | 0.9557 |
| Without Modular | 0.0653 | 0.1164 | 0.2548 | 0.2787 | 0.3145 | 0.4635 | 0.6315 | 0.7164 | 0.6627 | 0.6794 | 0.6664 |
| | 0.0030 | 0.1970 | 0.1997 | 0.2830 | 0.3479 | 0.3982 | 0.6812 | 0.6627 | 0.7635 | 0.7479 | 0.7935 |
| | 0.1185 | 0.2136 | 0.2145 | 0.2479 | 0.4330 | 0.3479 | 0.5646 | 0.6987 | 0.6971 | 0.5485 | 0.7789 |

**Table S4.** Normalized fluorescence intensity of allosteric aptamer with varying P2 lengths.

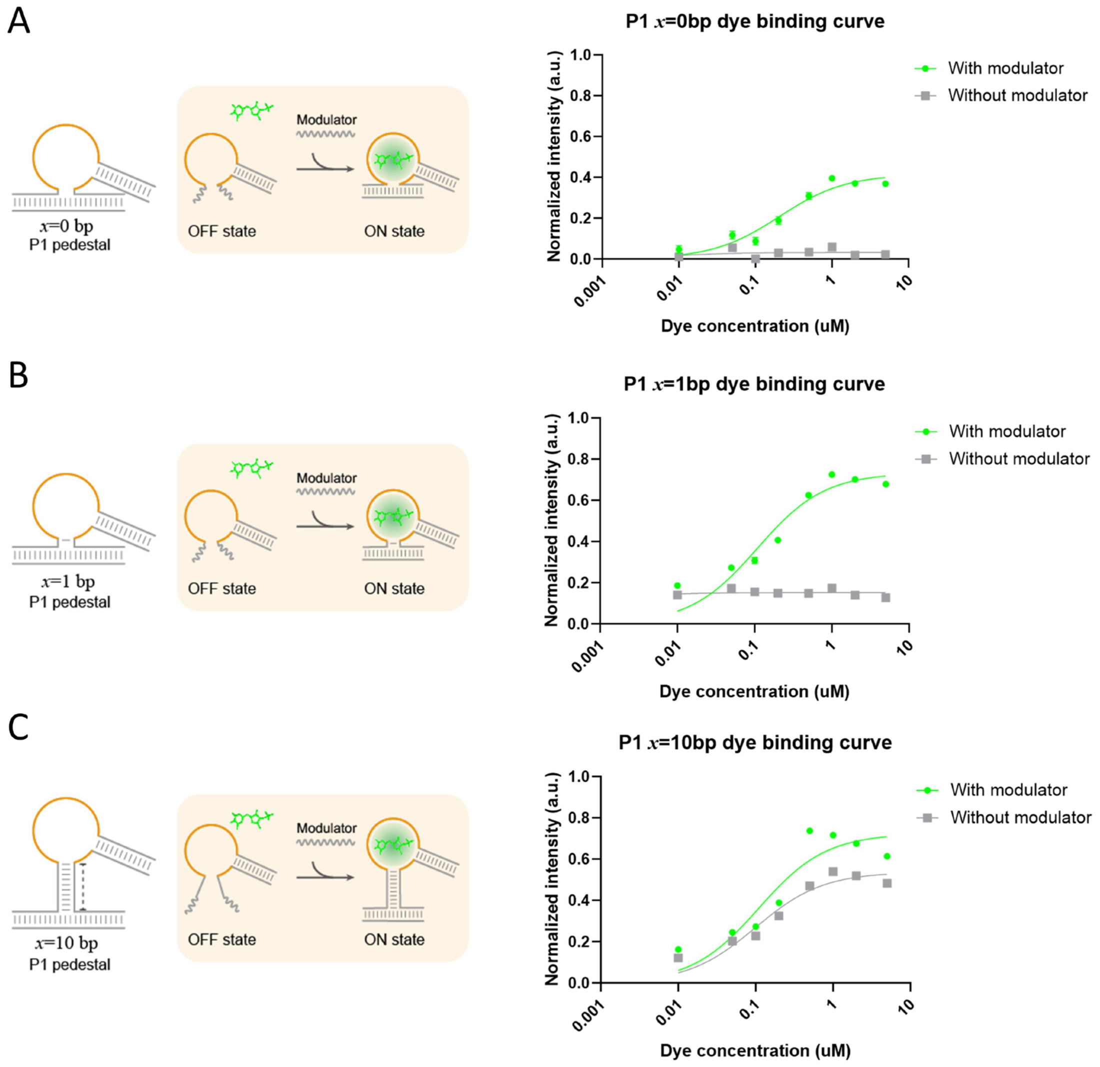

**Figure S7.** Schematics and DFHBI-1T binding curves of three representative allosteric aptamers (**A**) P1 $x$= 0 bp, (**B**) P1 $x$= 1 bp, and (**C**) P1 $x$= 10 bp in the absence (grey) and presence (green) of the modulator strand.

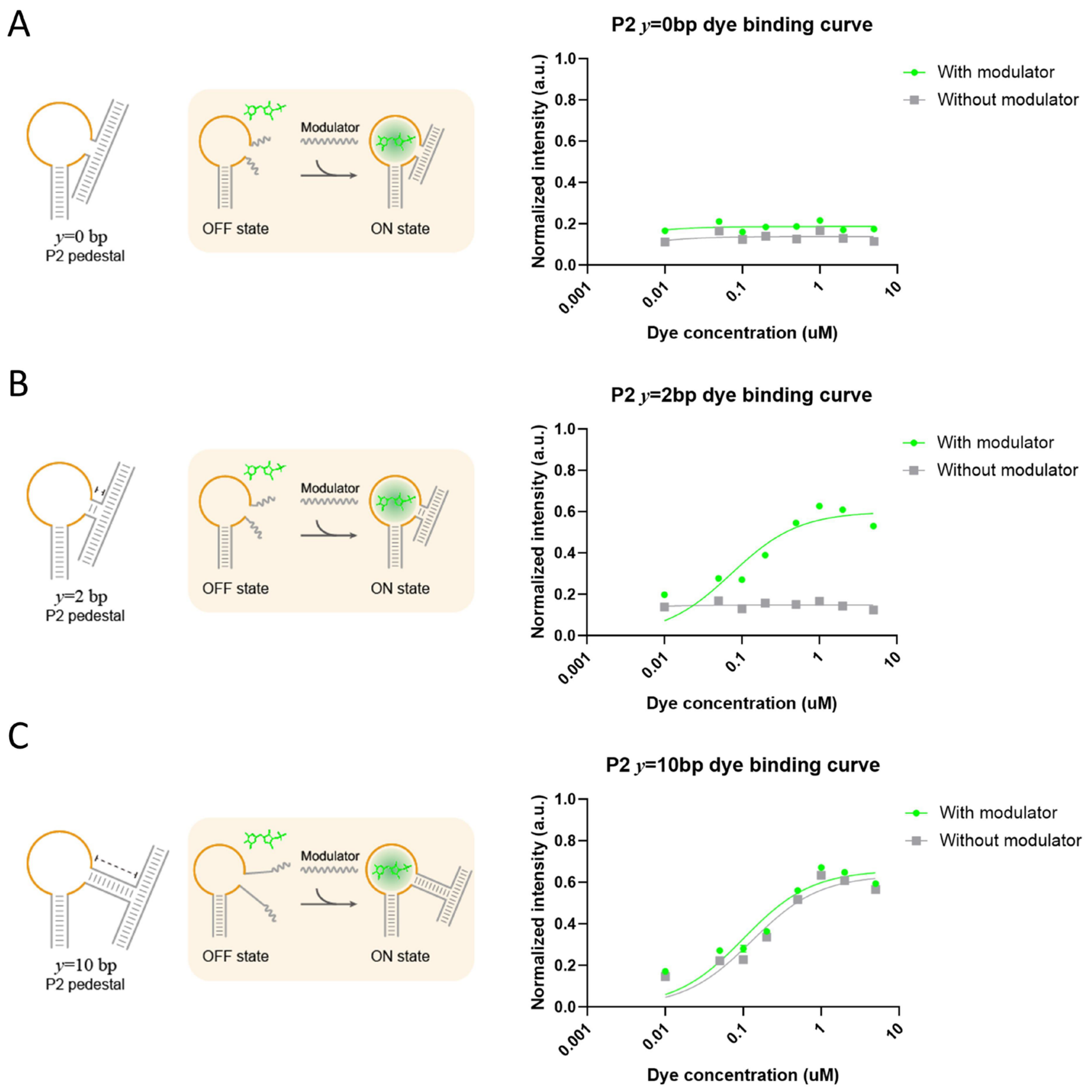

**Figure S8.** Schematics and DFHBI-1T binding curves of three representative allosteric aptamers (**A**) P2 *y*= 0 bp, (**B**) P2 *y*= 2 bp, and (**C**) P2 *y*= 10 bp in the absence (grey) and presence (green) of the modulator strand.

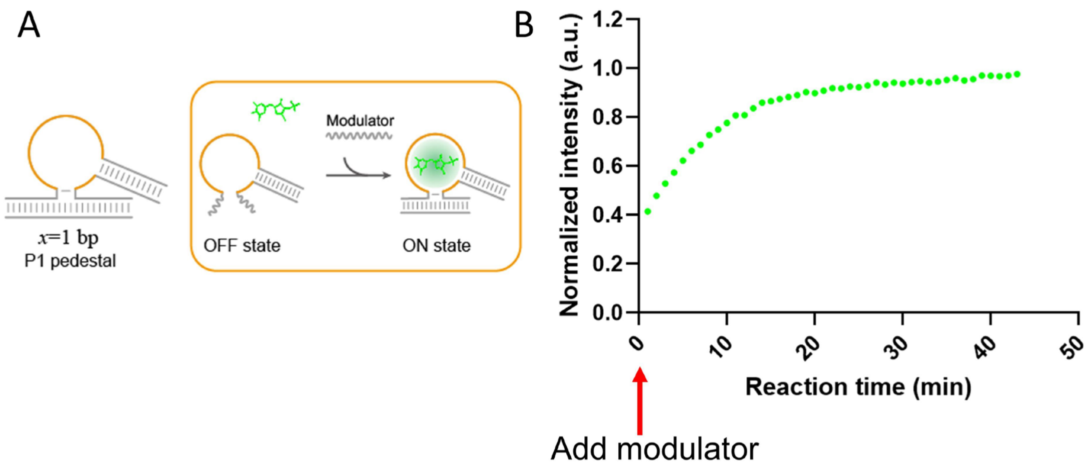

**Figure S9.** Schematics (**A**) and kinetic traces (**B**) showing the signal increasement upon adding modulator strand into allosteric aptamer system (P1 *x*= 1 bp).

It's worth noting that the binding of modulator to allosteric aptamer is very quick, once modulator is added the transition from OFF state to ON state will immediately initiate. It will take some time to add modulator into the solution and mix them thoroughly and put the plate in fluorescence reader, which will collectively result in several minutes delay of fluorescence recording, so the real initiation of reaction is actually earlier than "0" min.

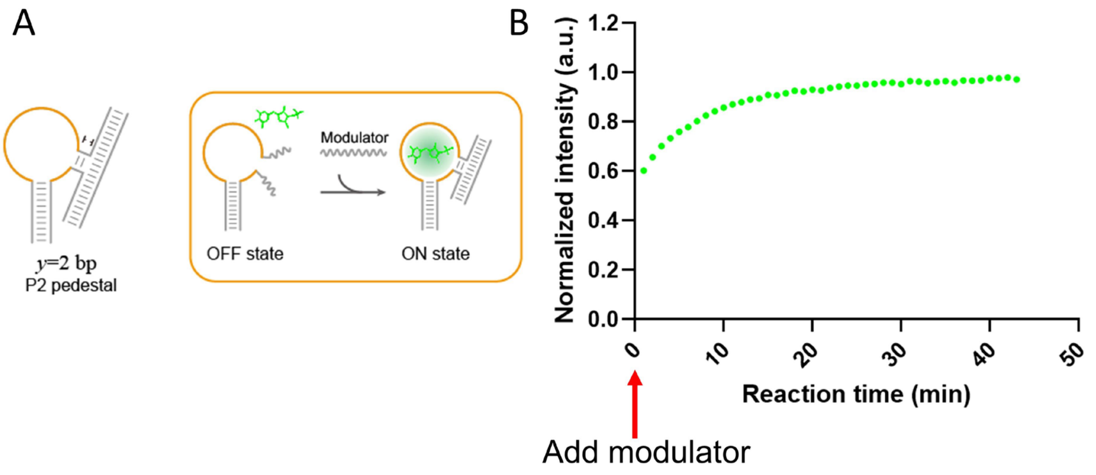

**Figure S10.** Schematics (**A**) and kinetic traces (**B**) showing the signal increasement upon adding modulator strand into allosteric aptamer system (P2 *y*= 2 bp).

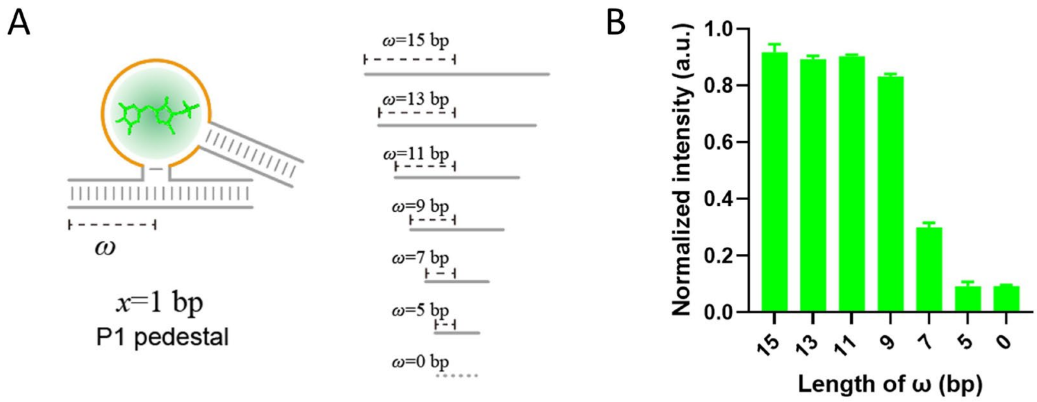

**Figure S11.** Schematics (**A**) and statistical analysis (**B**) showing the fluorescence signal plotted against the length of modulator binding region (ω) in the allosteric aptamer system (P1 *x*= 1 bp).

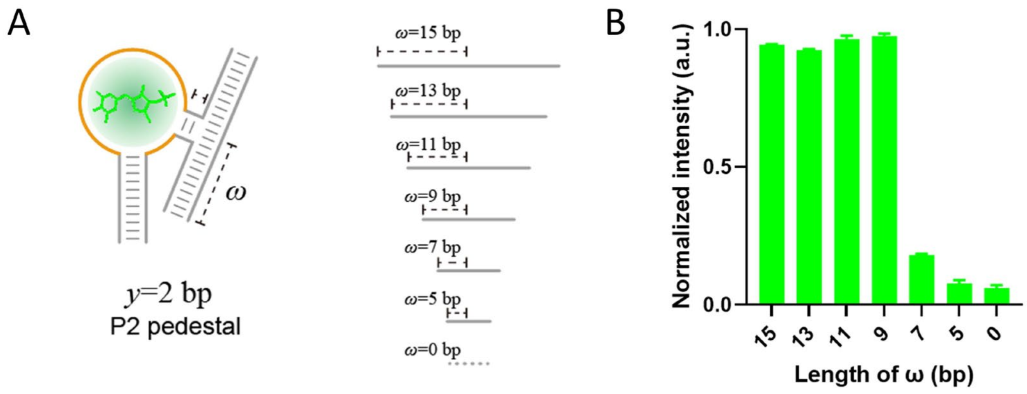

**Figure S12.** Schematics (**A**) and statistical analysis (**B**) showing the fluorescence signal plotted against the length of modulator binding region (ω) in the allosteric aptamer system (P2 *y*= 2 bp).

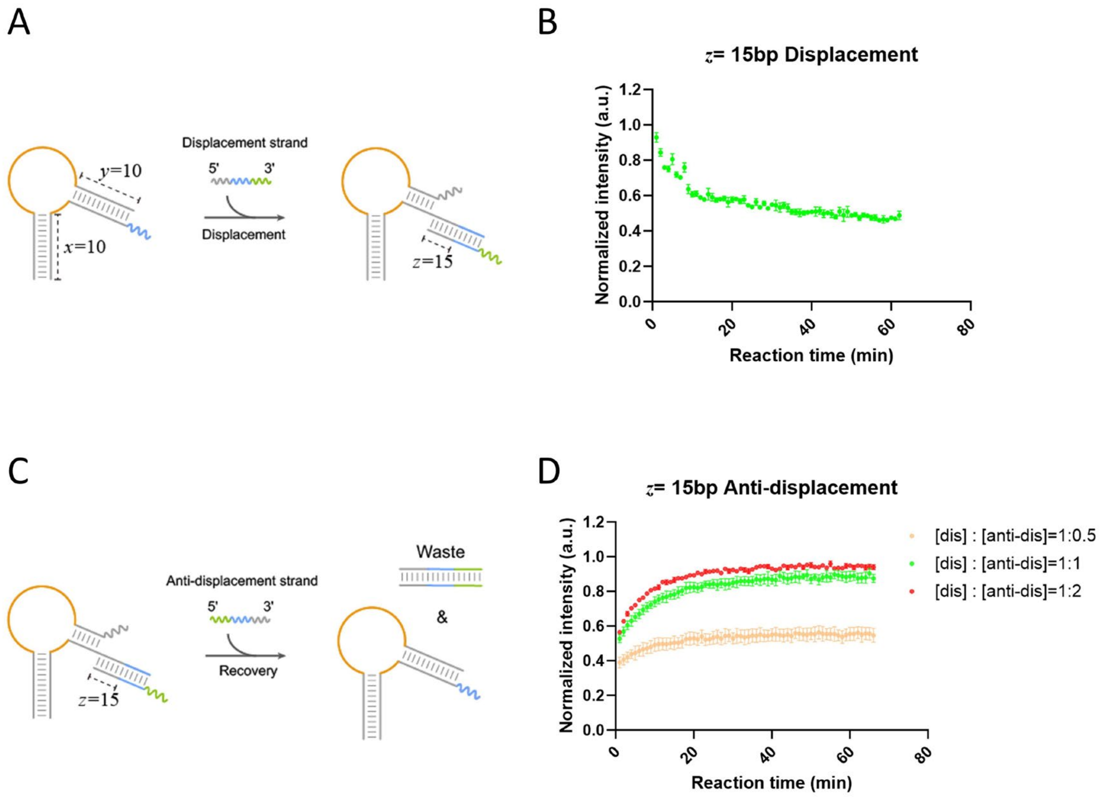

**Figure S13.** (**A-B**) Schematics and kinetic traces showing the signal change upon adding displacement strand into allosteric aptamer system (z= 15 bp). (**C-D**) Schematics and kinetic traces showing the signal change upon adding anti-displacement strand to recover the function of allosteric aptamer.

Normalized intensity of reverse allosteric variants (3 replicates)

| z (bp) | 0 bp | 1 bp | 2 bp | 3 bp | 4 bp | 5 bp | 6 bp | 7 bp | 8 bp | 9 bp | 10 bp | 11 bp | 12 bp | 13 bp | 14 bp | 15 bp |
|---|---|---|---|---|---|---|---|---|---|---|---|---|---|---|---|---|
| displacement | 0.6455 | 0.6888 | 0.8728 | 0.7834 | 0.8747 | 0.7299 | 0.7474 | 0.8256 | 0.7348 | 0.5296 | 0.3235 | 0.3111 | 0.2016 | 0.1282 | 0.1572 | 0.0653 |
| | 0.5050 | 0.6006 | 0.7366 | 0.7945 | 0.7322 | 0.8051 | 0.6839 | 0.7106 | 0.7312 | 0.5316 | 0.3542 | 0.2904 | 0.1958 | 0.1441 | 0.0975 | 0.0384 |
| | 0.6461 | 0.5987 | 0.6691 | 0.7186 | 0.7621 | 0.7803 | 0.7453 | 0.7331 | 0.7440 | 0.4926 | 0.4181 | 0.3583 | 0.2485 | 0.1851 | 0.1939 | 0.0875 |
| recovery | 0.7401 | 0.7089 | 0.8383 | 0.8303 | 0.8702 | 0.7753 | 0.8203 | 0.6041 | 0.7037 | 0.9072 | 0.8762 | 0.7317 | 0.7245 | 0.5116 | 0.7845 | 0.9177 |
| | 0.5924 | 0.7136 | 0.7977 | 0.8292 | 0.8825 | 0.6473 | 0.7895 | 0.7967 | 0.7889 | 0.8638 | 0.9138 | 0.5936 | 0.8843 | 0.5250 | 0.7765 | 0.8017 |
| | 0.7074 | 0.7435 | 0.7187 | 0.9333 | 0.8273 | 0.7379 | 0.8527 | 0.7597 | 0.8345 | 0.6867 | 0.6883 | 0.8359 | 0.6694 | 0.5285 | 0.4776 | 0.6983 |

**Table S5.** Normalized fluorescence intensity of allosteric aptamer designs with reverse regulation.

Normalized intensity of Boolean logic gates (3 replicates)

| Input | [0, 0] | [1, 0] | [0, 1] | [1, 1] |
|---|---|---|---|---|
| AND | 0.0282 | 0.0127 | 0.1324 | 0.5179 |
| | 0.1226 | 0.1346 | 0.2267 | 0.8449 |
| | 0.0269 | 0.0669 | 0.2330 | 0.8062 |
| OR | 0.0827 | 0.7236 | 0.5370 | 0.7186 |
| | 0.0761 | 0.6308 | 0.5979 | 0.8018 |
| | 0.1280 | 0.6397 | 0.7254 | 0.6591 |
| XOR | 0.0162 | 0.7117 | 0.6277 | 0.2679 |
| | 0.0405 | 0.7534 | 0.6127 | 0.2931 |
| | 0.0750 | 0.6174 | 0.7227 | 0.2719 |
| NAND | 0.6675 | 0.6118 | 0.5787 | 0.1455 |
| | 0.8903 | 0.6395 | 0.5048 | 0.2494 |
| | 0.9373 | 0.8369 | 0.8285 | 0.1921 |
| NOR | 0.6887 | 0.0595 | 0.0630 | 0.0874 |
| | 0.6280 | 0.0194 | 0.1258 | 0.0760 |
| | 0.7483 | 0.1003 | 0.1765 | 0.0628 |
| XNOR | 0.6358 | 0.3912 | 0.2465 | 0.6535 |
| | 0.7142 | 0.4153 | 0.2605 | 0.9484 |
| | 1.0099 | 0.3535 | 0.2842 | 0.9723 |

**Table S6.** Normalized fluorescence intensity of allosteric aptamer for implementing six kinds of Boolean logic gates.

Normalized intensity of orthogonality designs (3 replicates)

| | M-1 | M-2 | M-3 | M-4 | M-5 | M-6 | M-7 | M-8 | M-9 | M-10 |
|---|---|---|---|---|---|---|---|---|---|---|
| Apt-1 | 0.3521 | -0.0395 | -0.0258 | -0.0593 | -0.0166 | -0.0357 | -0.0210 | 0.0027 | -0.0409 | 0.0130 |
| | 0.2597 | 0.0610 | 0.0136 | -0.0230 | 0.0428 | -0.0085 | 0.0088 | 0.0180 | -0.0354 | 0.0009 |
| | 0.3881 | 0.0103 | -0.2573 | -0.0572 | -0.0117 | 0.0172 | 0.0513 | 0.0155 | -0.0260 | -0.0069 |
| Apt-2 | -0.0075 | 0.4421 | 0.0025 | -0.0213 | 0.0038 | -0.0534 | -0.0117 | -0.1165 | -0.0661 | 0.1340 |
| | -0.0377 | 0.6450 | -0.0013 | 0.0866 | 0.0373 | -0.0722 | -0.0861 | -0.0261 | 0.0345 | -0.0244 |
| | 0.0984 | 0.7666 | 0.0004 | 0.0191 | 0.0263 | -0.0467 | 0.0826 | 0.1180 | 0.0742 | 0.0500 |
| Apt-3 | -0.0190 | 0.0063 | 0.4346 | -0.0892 | 0.0577 | -0.0177 | -0.0030 | 0.0163 | -0.1637 | -0.0459 |
| | -0.1549 | 0.0194 | 0.4817 | 0.0272 | -0.0216 | 0.0094 | 0.0400 | 0.0146 | 0.0585 | 0.0002 |
| | 0.0217 | -0.0271 | 0.5218 | 0.0309 | 0.0188 | -0.0037 | 0.0112 | 0.1155 | 0.0076 | 0.0022 |
| Apt-4 | 0.0043 | -0.0361 | 0.0153 | 0.8448 | 0.0097 | 0.1116 | 0.0353 | -0.0416 | 0.0644 | 0.0454 |
| | 0.0216 | 0.0392 | 0.1243 | 0.7440 | 0.0063 | 0.0434 | 0.0207 | -0.0638 | 0.0124 | 0.0701 |
| | 0.0160 | 0.0063 | -0.0604 | 0.9233 | 0.0911 | 0.0136 | 0.0277 | -0.0087 | 0.1556 | 0.0286 |
| Apt-5 | 0.0071 | 0.0423 | -0.0299 | -0.0256 | 0.5134 | -0.0201 | -0.0089 | 0.0049 | 0.0078 | -0.0110 |
| | -0.0679 | 0.0297 | 0.0406 | -0.0216 | 0.5036 | -0.0577 | -0.0061 | 0.1626 | -0.0145 | -0.0096 |
| | 0.0445 | 0.0059 | 0.0348 | 0.0089 | 0.4196 | 0.0390 | 0.0433 | -0.0228 | 0.0219 | 0.0029 |
| Apt-6 | -0.0186 | -0.0329 | -0.0721 | 0.0112 | -0.0550 | 0.5632 | 0.0193 | -0.0406 | -0.0072 | 0.0186 |
| | -0.0022 | -0.0397 | 0.0441 | -0.0388 | -0.0252 | 0.4953 | -0.0530 | 0.0120 | -0.0033 | -0.3320 |
| | 0.0001 | -0.0442 | -0.0311 | -0.0507 | -0.0148 | 0.5728 | 0.0057 | -0.1177 | -0.0315 | 0.0308 |
| Apt-7 | 0.0466 | -0.0069 | 0.0215 | 0.0850 | 0.1979 | 0.1085 | 0.5443 | 0.0966 | 0.0804 | 0.0315 |
| | 0.0660 | 0.1161 | 0.0814 | 0.0589 | 0.0905 | 0.1562 | 0.6003 | 0.0928 | 0.0867 | -0.0094 |
| | 0.0497 | 0.0600 | 0.0344 | 0.0092 | 0.0107 | 0.1506 | 0.6766 | -0.0259 | 0.0218 | 0.0477 |
| Apt-8 | 0.0057 | -0.0323 | -0.0339 | -0.0798 | -0.0203 | -0.0338 | -0.0232 | 0.5251 | 0.0301 | 0.0069 |
| | -0.0672 | 0.0038 | -0.0307 | -0.0833 | 0.0241 | 0.0046 | 0.0296 | 0.6595 | 0.0171 | -0.0422 |
| | 0.2812 | -0.0159 | -0.0135 | 0.0178 | -0.0282 | 0.0438 | -0.0206 | 0.6962 | 0.0535 | -0.0338 |
| Apt-9 | -0.0137 | 0.0443 | 0.0006 | 0.0221 | 0.0039 | 0.0967 | 0.1325 | 0.0360 | 0.3804 | 0.0222 |
| | -0.0528 | 0.1093 | 0.0544 | 0.0262 | 0.0365 | 0.0590 | 0.1013 | -0.0176 | 0.5407 | 0.0576 |
| | 0.0009 | 0.0844 | -0.0161 | 0.0964 | 0.1226 | 0.1207 | 0.0203 | 0.0486 | 0.3830 | 0.0565 |
| Apt-10 | -0.0286 | -0.0280 | -0.0149 | 0.1075 | -0.0620 | -0.0787 | 0.0646 | 0.0097 | 0.0791 | 0.5674 |
| | -0.0257 | -0.0363 | 0.0349 | -0.0313 | -0.0378 | 0.0120 | -0.0522 | -0.0461 | 0.0293 | 0.4648 |
| | 0.0138 | 0.0202 | 0.0000 | 0.0140 | -0.0580 | 0.1009 | 0.0567 | 0.0042 | 0.0547 | 0.7233 |

**Table S7.** Normalized fluorescence intensity of allosteric aptamer for orthogonality analysis.

Normalized intensity of circuits (3 replicates)

| Input | [0, 0, 0] | [1, 0, 0] | [0, 1, 0] | [0, 0, 1] | [1, 1, 0] | [1, 0, 1] | [0, 1, 1] | [1, 1, 1] |
|---|---|---|---|---|---|---|---|---|
| BUFFER | 0.1580 | 0.4002 | 0.5239 | 0.5714 | 0.6816 | 0.7807 | 0.8258 | 0.9367 |
| | 0.0498 | 0.4201 | 0.5021 | 0.4506 | 0.4577 | 0.7886 | 0.6294 | 0.8100 |
| | 0.1141 | 0.4304 | 0.5219 | 0.4800 | 0.7262 | 0.8332 | 0.7397 | 0.9105 |
| MAJORITY | 0.0534 | 0.1251 | 0.1369 | 0.0694 | 0.2936 | 0.3071 | 0.3136 | 0.7191 |
| | 0.0888 | 0.0653 | 0.1060 | 0.0547 | 0.2905 | 0.4811 | 0.3035 | 0.6865 |
| | 0.0486 | 0.1352 | 0.0848 | 0.1564 | 0.3529 | 0.2723 | 0.3825 | 0.8449 |

**Table S8.** Normalized fluorescence intensity of allosteric aptamer for implementing combinatorial BUFFER function and MAJRITY function.

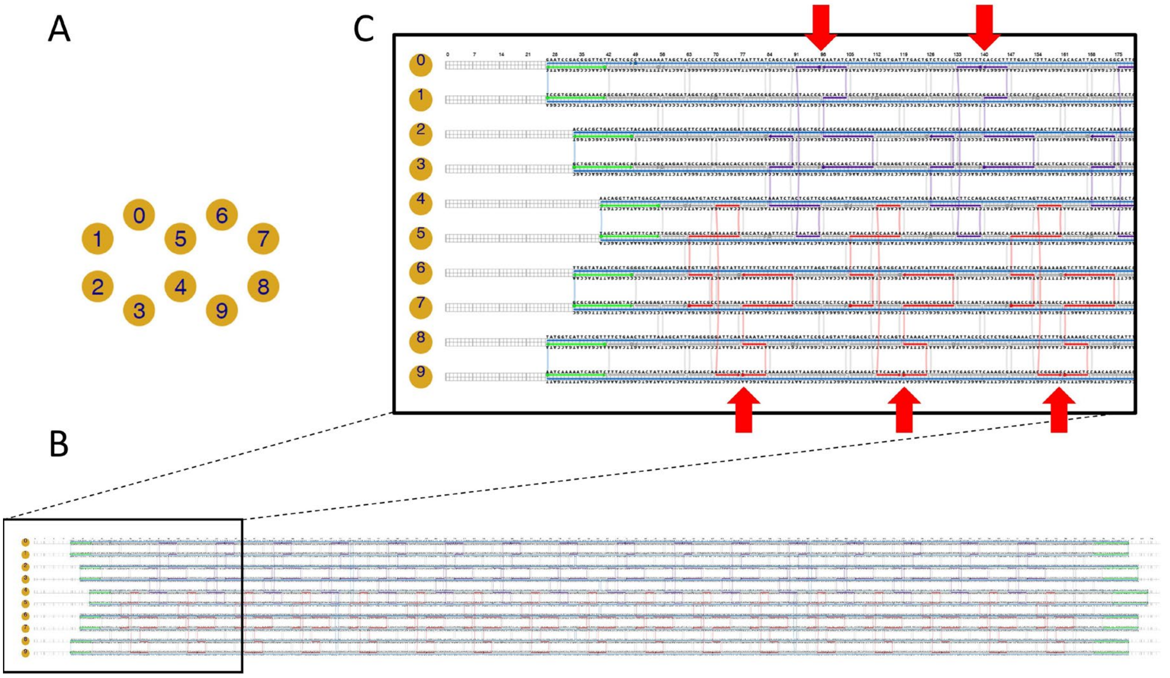

**Figure S14.** Schematics of 10-HB origami design. **A**, arrangement of ten helices in the formation of 10-HB origami. **B**, caDNAno[3, 4] design of 10-HB origami. **C**, zoom-in view of the staple arrangement and sequence. Red arrows: representative sites on helix-9 and helix-0 for incorporating allosteric aptamer or FAM-labeled probes.

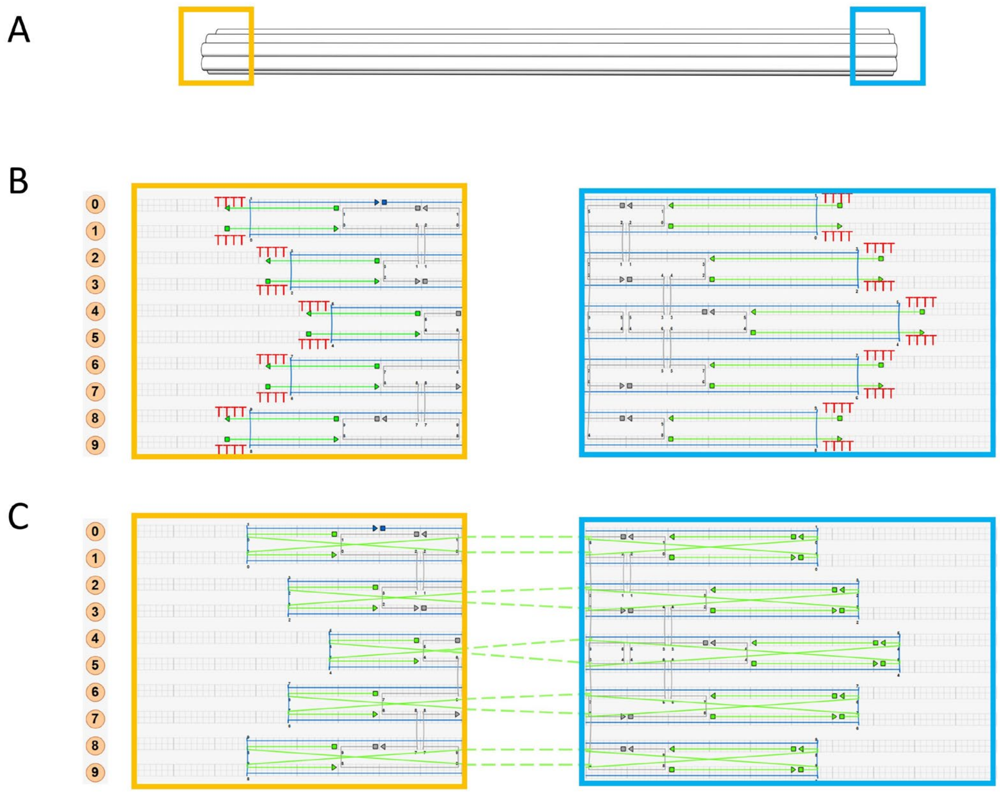

**Figure S15.** Schematics of 10-HB origami monomer and polymer design. **A-B**, zoom-in views showing left edge (orange) and right edge (blue) of the monomer 10-HB origami. Each edge staple has a “TTTT” tail to prevent edge-edge stacking. **C**, zoom-in views showing left edge (orange) and right edge (blue) of the individual 10-HB origami for polymeric design. Each staple at the left edge has a 4-nt overhang to seamlessly bind 4-nt vacancy at the right edge of the neighbor origami.

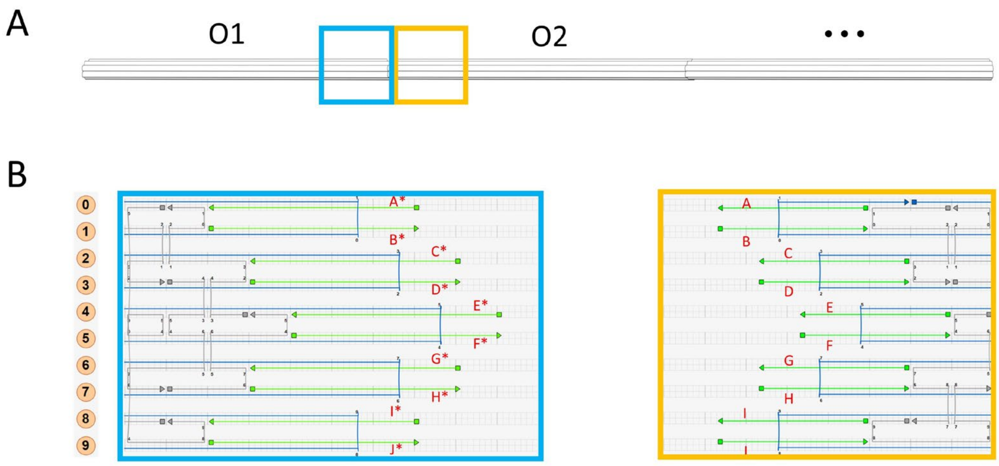

**Figure S16.** Schematics of 10-HB origami oligomer design. **A-B**, zoom-in views showing the design of left edge (orange) of O2 and right edge (blue) of O1 in 10-HB origami oligomers. Each staple at the left edge of O2 has a 10-nt sticky end to complementarily bind 10-nt sticky end at the right edge staples of the O1.

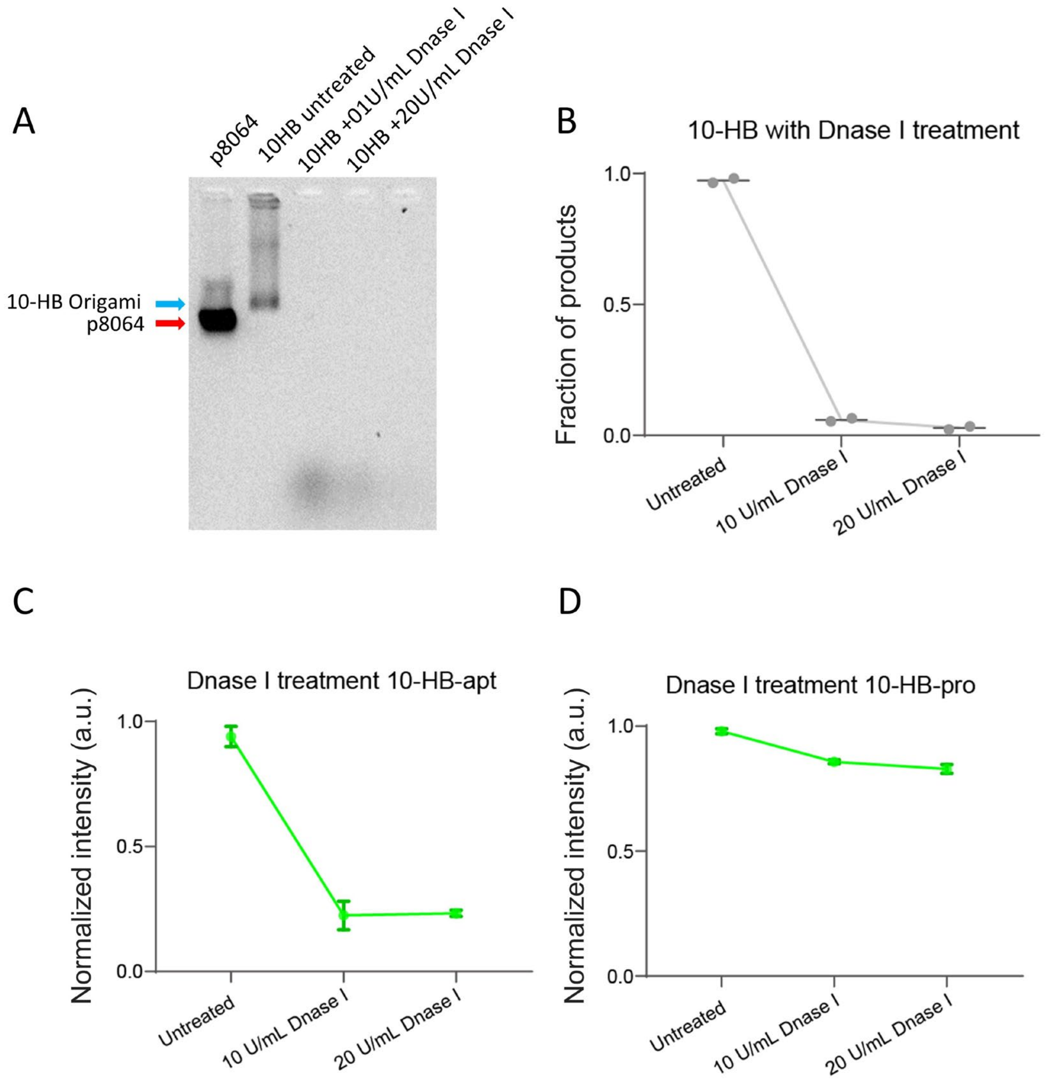

**Figure S17. A-B**, native agarose gel electrophoresis and statistical analysis of 10-HB origami treated with Dnase I. **C-D**, fluorescence analysis of 10-HB-apt and 10-HB-pro treated with Dnase I.

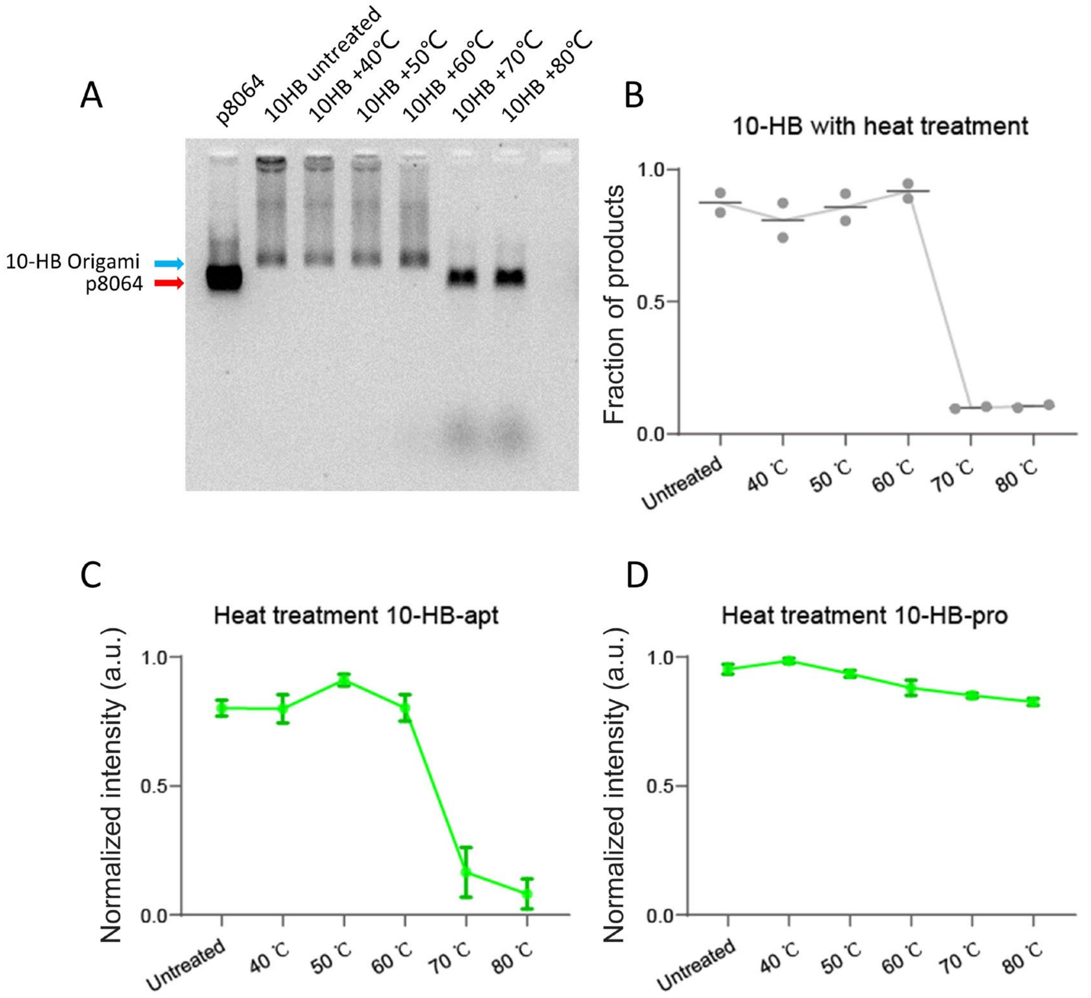

**Figure S18. A-B**, native agarose gel electrophoresis and statistical analysis of 10-HB origami with heat treatment. **C-D**, fluorescence analysis of 10-HB-apt and 10-HB-pro with heat treatment.

A

Normalized intensity of 10-HB-apt and 10-HB-pro
with Dnase I treatment (3 replicates)

| | untreated | 10 U/mL Dnase I | 20 U/mL Dnase I |
|---|---|---|---|
| 10HB-apt | 0.8613 | 0.2606 | 0.2081 |
| | 0.9631 | 0.1125 | 0.2444 |
| | 0.9956 | 0.3000 | 0.2469 |
| 10HB-pro | 0.9654 | 0.8443 | 0.8275 |
| | 0.9735 | 0.8563 | 0.7990 |
| | 0.9987 | 0.8716 | 0.8590 |

B

Normalized intensity of 10-HB-apt and 10-HB-pro
with heat treatment (3 replicates)

| Input | untreated | 40 ℃ | 50 ℃ | 60 ℃ | 70 ℃ | 80 ℃ |
|---|---|---|---|---|---|---|
| 10HB-apt | 0.7875 | 0.6913 | 0.8681 | 0.8312 | 0.1219 | 0.0194 |
| | 0.7569 | 0.8694 | 0.9431 | 0.7438 | 0.0981 | 0.089 |
| | 0.8613 | 0.8369 | 0.9225 | 0.8344 | 0.2750 | 0.1350 |
| 10HB-pro | 0.9164 | 0.9669 | 0.9101 | 0.8518 | 0.8616 | 0.8411 |
| | 0.9686 | 0.9988 | 0.9434 | 0.9115 | 0.8467 | 0.8215 |
| | 0.9750 | 0.9909 | 0.9527 | 0.8790 | 0.8445 | 0.8164 |

**Table S9.** Normalized fluorescence intensity of 10-HB-apt and 10-HB-pro with Dnase I treatment (**A**) and heat treatment (**B**).

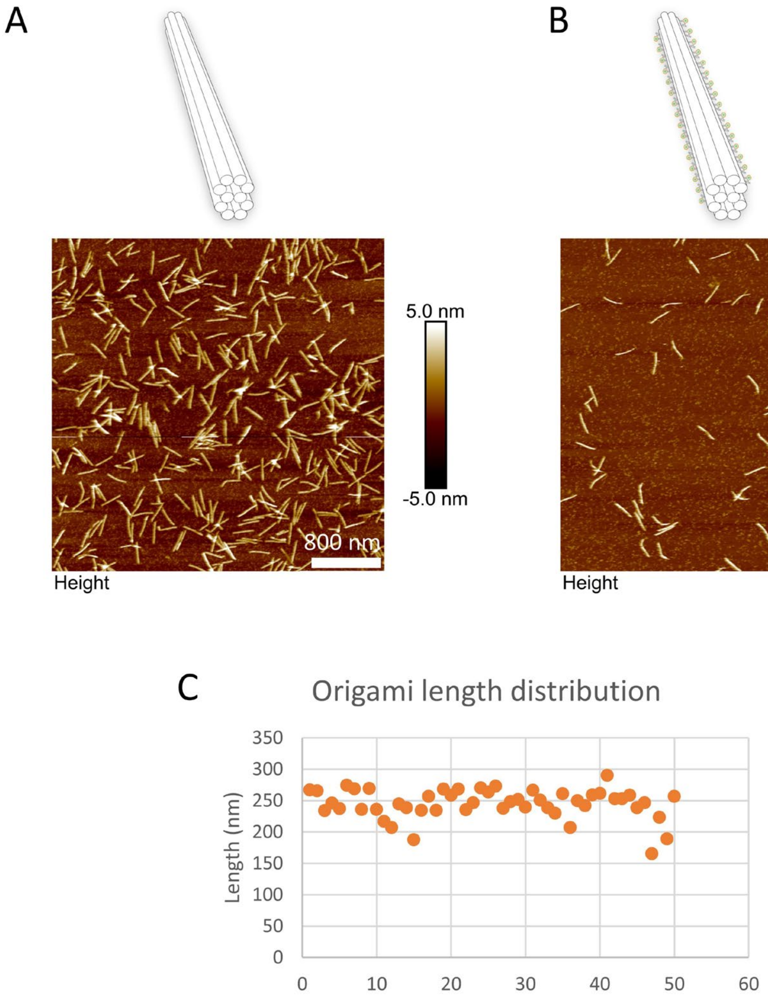

**Figure S19. A-B**, Representative atomic force microscopy (AFM) images of 10-HB origami (**A**) and 10-HB-apt origami (**B**) monomers. **C**, statistical analysis of the 10-HB origami length. The result shows that 10-HB-apt origami is ~250 nm long as expected. Scale bars: 800 nm.

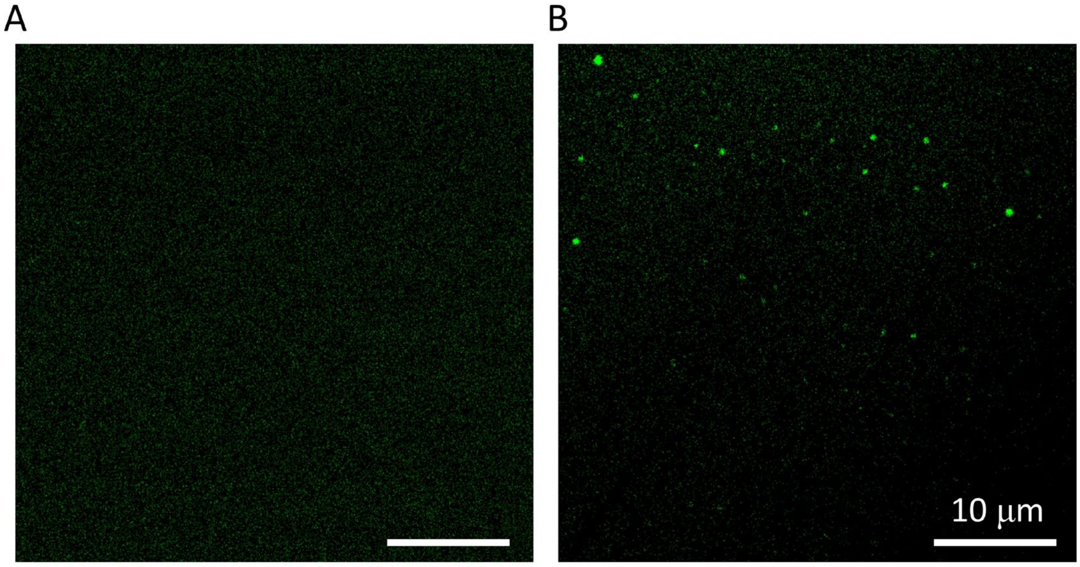

**Figure S20. A-B**, Representative fluorescence microscopy images of monomeric 10-HB origami before (**A**) and after (**B**) modulator binding. Scale bars: 10 μm.

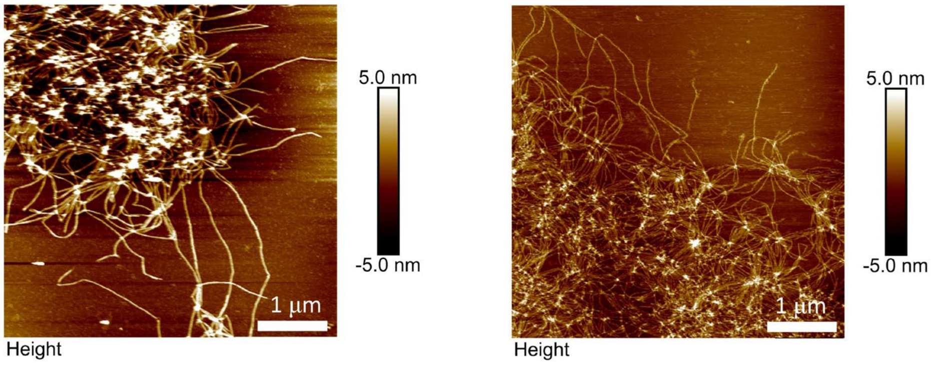

**Figure S21.** Representative atomic force microscopy (AFM) images of polymeric 10-HB-apt origami. Scale bars: 1 μm.

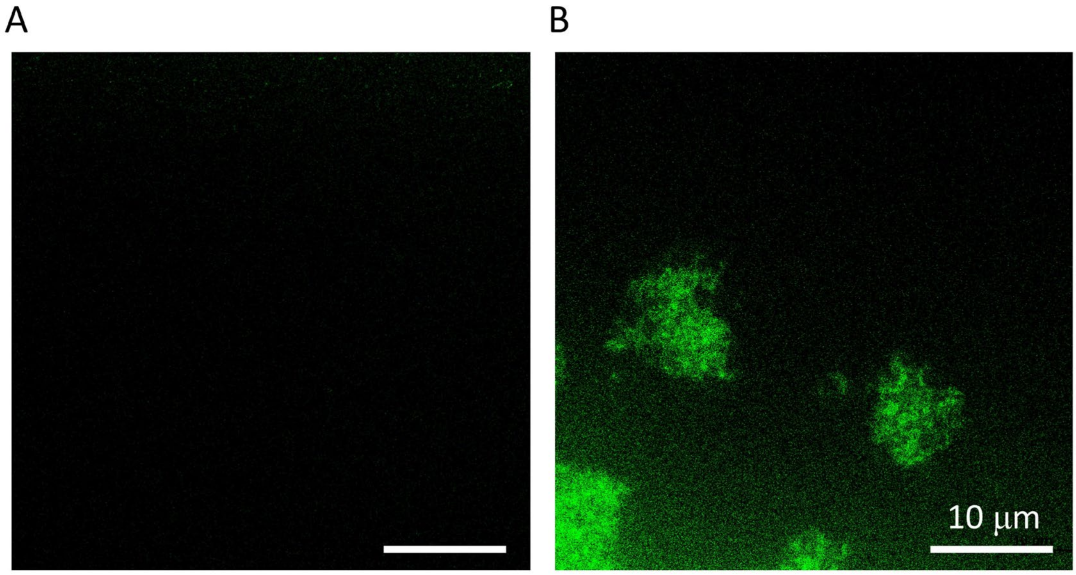

**Figure S22. A-B**, Representative fluorescence microscopy images of polymeric 10-HB origami before (**A**) and after (**B**) modulator binding. Scale bars: 10 μm.

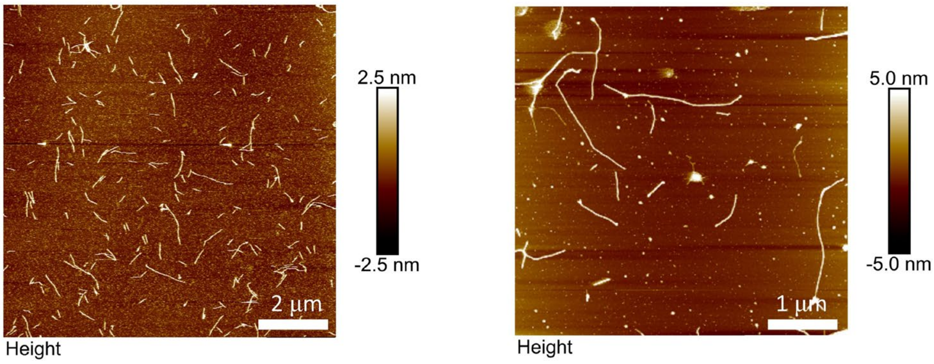

**Figure S23.** Representative atomic force microscopy (AFM) images of oligomeric 10-HB-apt origami. Scale bars: 1 µm (left) and 1 µm (right).

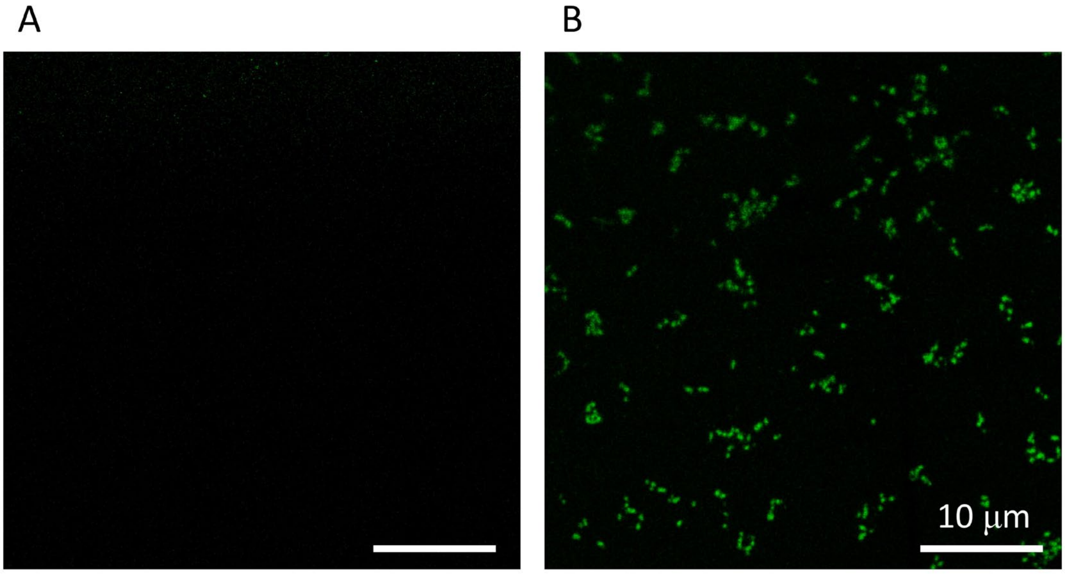

**Figure S24. A-B**, Representative fluorescence microscopy images of oligomeric 10-HB origami before (**A**) and after (**B**) modulator binding. Scale bars: 10 μm.

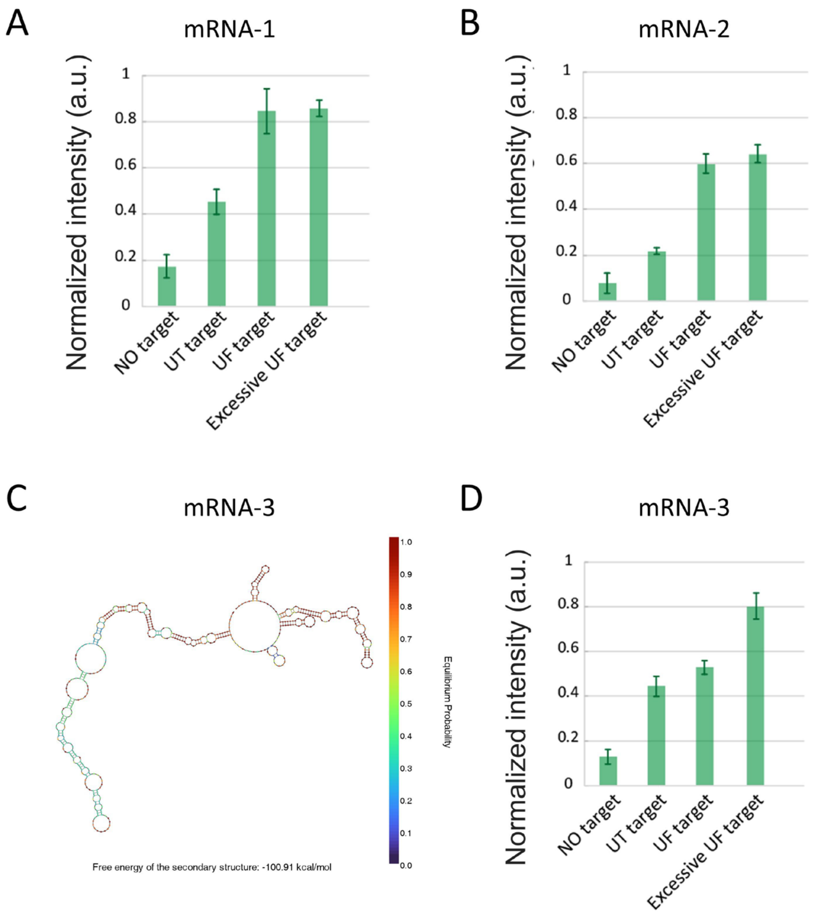

**Figure S25. A-B**, statistical analysis of fluorescence intensity of 10-HB origami before and after binding to target mRNA-1 (**A**) and mRNA-2 (**B**). **C**, schematics of the secondary and energy landscape of mRNA-3. **D**, statistical analysis of fluorescence intensity of 10-HB origami before and after binding to target mRNA-3. UT target, untreated mRNA target; UF, unfolded mRNA targets after heating and then crash-cooling on ice. Errors bars, mean ± S.E.M..

Normalized intensity of 10-HB targeting RNA (3 replicates)

| Input | No target | UT target | UF target | Excessive UF target |
|---|---|---|---|---|
| mRNA-1 | 0.1302 | 0.3921 | 0.8680 | 0.8446 |
| | 0.1628 | 0.4799 | 0.7407 | 0.8333 |
| | 0.2304 | 0.4844 | 0.9308 | 0.8982 |
| mRNA-2 | 0.1214 | 0.2403 | 0.6470 | 0.6928 |
| | 0.0883 | 0.2094 | 0.5654 | 0.6258 |
| | 0.0347 | 0.2131 | 0.5956 | 0.6251 |
| mRNA-3 | 0.0987 | 0.4077 | 0.5236 | 0.8148 |
| | 0.1593 | 0.4955 | 0.5062 | 0.7453 |
| | 0.1502 | 0.4458 | 0.5685 | 0.8605 |

**Table S10.** Normalized fluorescence intensity of 10-HB origami targeting mRNA.

Normalized intensity of 10-HB light-up cycles (3 replicates)

| Input | Initial state | 1st cycle +RNA | 1st cycle +RNase H1 | 2nd cycle +RNA | 2nd cycle +RNase H1 |
|---|---|---|---|---|---|
| 10-HB apt | 0.0832 | 0.8728 | 0.1443 | 0.6838 | 0.1769 |
| | 0.1222 | 0.8202 | 0.1537 | 0.6274 | 0.1920 |
| | 0.0634 | 0.8088 | 0.2139 | 0.6988 | 0.2134 |

**Table S11.** Normalized fluorescence intensity of 10-HB-apt that lights-up cyclically.

### Sequences (5' to 3')

#### Allosteric aptamer sequences:

| Strand name | Sequence (5' to 3') |
|---|---|
| AlloApt-P1-modulator | GAAGTGTATTCCAGGCGAGCGACATTTGAC |
| AlloApt-P1-10nt-1 | GTCAAATGTCGCTCGtTTCGGCAAGAAGGGATGATGCGGCAGTGGGCTTTGCATAACTC |
| AlloApt-P1-10nt-2 | GAGTTATGCAAGGGGTCTTGCCGAAtCCTGGAATACACTTC |
| AlloApt-P1-9nt-1 | GTCAAATGTCGCTCGtTCGGCAAGAAGGGATGATGCGGCAGTGGGCTTTGCATAACTC |
| AlloApt-P1-9nt-2 | GAGTTATGCAAGGGGTCTTGCCGAtCCTGGAATACACTTC |
| AlloApt-P1-8nt-1 | GTCAAATGTCGCTCGtCGGCAAGAAGGGATGATGCGGCAGTGGGCTTTGCATAACTC |
| AlloApt-P1-8nt-2 | GAGTTATGCAAGGGGTCTTGCCGtCCTGGAATACACTTC |
| AlloApt-P1-7nt-1 | GTCAAATGTCGCTCGtGGCAAGAAGGGATGATGCGGCAGTGGGCTTTGCATAACTC |
| AlloApt-P1-7nt-2 | GAGTTATGCAAGGGGTCTTGCCtCCTGGAATACACTTC |
| AlloApt-P1-6nt-1 | GTCAAATGTCGCTCGtGCAAGAAGGGATGATGCGGCAGTGGGCTTTGCATAACTC |
| AlloApt-P1-6nt-2 | GAGTTATGCAAGGGGTCTTGCtCCTGGAATACACTTC |
| AlloApt-P1-5nt-1 | GTCAAATGTCGCTCGtCAAGAAGGGATGATGCGGCAGTGGGCTTTGCATAACTC |
| AlloApt-P1-5nt-2 | GAGTTATGCAAGGGGTCTTGtCCTGGAATACACTTC |
| AlloApt-P1-4nt-1 | GTCAAATGTCGCTCGtAAGAAGGGATGATGCGGCAGTGGGCTTTGCATAACTC |
| AlloApt-P1-4nt-2 | GAGTTATGCAAGGGGTCTTtCCTGGAATACACTTC |
| AlloApt-P1-3nt-1 | GTCAAATGTCGCTCGtAGAAGGGATGATGCGGCAGTGGGCTTTGCATAACTC |
| AlloApt-P1-3nt-2 | GAGTTATGCAAGGGGTCTtCCTGGAATACACTTC |
| AlloApt-P1-2nt-1 | GTCAAATGTCGCTCGtGAAGGGATGATGCGGCAGTGGGCTTTGCATAACTC |
| AlloApt-P1-2nt-2 | GAGTTATGCAAGGGGTCtCCTGGAATACACTTC |

| | |
|---|---|
| AlloApt-P1-1nt-1 | GTCAAATGTCGCTCGtAAGGGATGATGCGGCAGTGGGCTTTGCATAACTC |
| AlloApt-P1-1nt-2 | GAGTTATGCAAGGGGTtCCTGGAATACACTTC |
| AlloApt-P1-0nt-1 | GTCAAATGTCGCTCGtAGGGATGATGCGGCAGTGGGCTTTGCATAACTC |
| AlloApt-P1-0nt-2 | GAGTTATGCAAGGGGtCCTGGAATACACTTC |
| AlloApt-P2-modulator | GAAGTGTATTCCAGGCGAGCGACATTTGAC |
| AlloApt-P2-10nt-1 | TTCGGCAAGAAGGGATGATGCGGCAGTGGGCTTTGCATAACTCtCCTGGAATACACTTC |
| AlloApt-P2-10nt-2 | GTCAAATGTCGCTCGtGAGTTATGCAAGGGGTCTTGCCGAA |
| AlloApt-P2-9nt-1 | TTCGGCAAGAAGGGATGATGCGGCAGTGGGCTTTGCATAACTtCCTGGAATACACTTC |
| AlloApt-P2-9nt-2 | GTCAAATGTCGCTCGtAGTTATGCAAGGGGTCTTGCCGAA |
| AlloApt-P2-8nt-1 | TTCGGCAAGAAGGGATGATGCGGCAGTGGGCTTTGCATAACtCCTGGAATACACTTC |
| AlloApt-P2-8nt-2 | GTCAAATGTCGCTCGtGTTATGCAAGGGGTCTTGCCGAA |
| AlloApt-P2-7nt-1 | TTCGGCAAGAAGGGATGATGCGGCAGTGGGCTTTGCATAAtCCTGGAATACACTTC |
| AlloApt-P2-7nt-2 | GTCAAATGTCGCTCGtTTATGCAAGGGGTCTTGCCGAA |
| AlloApt-P2-6nt-1 | TTCGGCAAGAAGGGATGATGCGGCAGTGGGCTTTGCATAtCCTGGAATACACTTC |
| AlloApt-P2-6nt-2 | GTCAAATGTCGCTCGtTATGCAAGGGGTCTTGCCGAA |
| AlloApt-P2-5nt-1 | TTCGGCAAGAAGGGATGATGCGGCAGTGGGCTTTGCATtCCTGGAATACACTTC |
| AlloApt-P2-5nt-2 | GTCAAATGTCGCTCGtATGCAAGGGGTCTTGCCGAA |
| AlloApt-P2-4nt-1 | TTCGGCAAGAAGGGATGATGCGGCAGTGGGCTTTGCAtCCTGGAATACACTTC |
| AlloApt-P2-4nt-2 | GTCAAATGTCGCTCGtTGCAAGGGGTCTTGCCGAA |
| AlloApt-P2-3nt-1 | TTCGGCAAGAAGGGATGATGCGGCAGTGGGCTTTGCtCCTGGAATACACTTC |
| AlloApt-P2-3nt-2 | GTCAAATGTCGCTCGtGCAAGGGGTCTTGCCGAA |

| | |
|---|---|
| AlloApt-P2-2nt-1 | TTCGGCAAGAAGGGATGATGCGGCAGTGGGCTTTGtCCTGGAATACACTTC |
| AlloApt-P2-2nt-2 | GTCAAATGTCGCTCGtCAAGGGGTCTTGCCGAA |
| AlloApt-P2-1nt-1 | TTCGGCAAGAAGGGATGATGCGGCAGTGGGCTTTtCCTGGAATACACTTC |
| AlloApt-P2-1nt-2 | GTCAAATGTCGCTCGtAAGGGGTCTTGCCGAA |
| AlloApt-P2-0nt-1 | TTCGGCAAGAAGGGATGATGCGGCAGTGGGCTTtCCTGGAATACACTTC |
| AlloApt-P2-0nt-2 | GTCAAATGTCGCTCGtAGGGGTCTTGCCGAA |
| Modulator (ω=15bp) | GAAGTGTATTCCAGGCGAGCGACATTTGAC |
| Modulator (ω=13bp) | AGTGTATTCCAGGCGAGCGACATTTG |
| Modulator (ω=11bp) | TGTATTCCAGGCGAGCGACATT |
| Modulator (ω=9bp) | TATTCCAGGCGAGCGACA |
| Modulator (ω=7bp) | TTCCAGGCGAGCGA |
| Modulator (ω=5bp) | CCAGGCGAGC |
| AlloApt-rev-1 | TTCGGCAAGAAGGGATGATGCGGCAGTGGGCTTTGCATAACTC |
| AlloApt-rev-2 | ctagagtgGAGTTATGCAAGGGGTCTTGCCGAA |
| Displacement-15nt | CCCCTTGCATAACTCcactctagTAGGTCCT |
| Recovery-15nt | AGGACCTActagagtgGAGTTATGCAAGGGG |
| Displacement-14nt | CCCTTGCATAACTCcactctagTAGGTCCT |
| Recovery-14nt | AGGACCTActagagtgGAGTTATGCAAGGG |
| Displacement-13nt | CCTTGCATAACTCcactctagTAGGTCCT |
| Recovery-13nt | AGGACCTActagagtgGAGTTATGCAAGG |
| Displacement-12nt | CTTGCATAACTCcactctagTAGGTCCT |

| | |
|---|---|
| Recovery-12nt | AGGACCTActagagtgGAGTTATGCAAG |
| Displacement-11nt | TTGCATAACTCcactctagTAGGTCCT |
| Recovery-11nt | AGGACCTActagagtgGAGTTATGCAA |
| Displacement-10nt | TGCATAACTCcactctagTAGGTCCT |
| Recovery-10nt | AGGACCTActagagtgGAGTTATGCA |
| Displacement-9nt | GCATAACTCcactctagTAGGTCCT |
| Recovery-9nt | AGGACCTActagagtgGAGTTATGC |
| Displacement-8nt | CATAACTCcactctagTAGGTCCT |
| Recovery-8nt | AGGACCTActagagtgGAGTTATG |
| Displacement-7nt | ATAACTCcactctagTAGGTCCT |
| Recovery-7nt | AGGACCTActagagtgGAGTTAT |
| Displacement-6nt | TAACTCcactctagTAGGTCCT |
| Recovery-6nt | AGGACCTActagagtgGAGTTA |
| Displacement-5nt | AACTCcactctagTAGGTCCT |
| Recovery-5nt | AGGACCTActagagtgGAGTT |
| Displacement-4nt | ACTCcactctagTAGGTCCT |
| Recovery-4nt | AGGACCTActagagtgGAGT |
| Displacement-3nt | CTCcactctagTAGGTCCT |
| Recovery-3nt | AGGACCTActagagtgGAG |
| Displacement-2nt | TCcactctagTAGGTCCT |
| Recovery-2nt | AGGACCTActagagtgGA |
| Displacement-1nt | CcactctagTAGGTCCT |
| Recovery-1nt | AGGACCTActagagtgG |
| Displacement-0nt | cactctagTAGGTCCT |
| Recovery-0nt | AGGACCTActagagtg |
| AND-apt-1 | TTCGGCAAGAAGGGATGATGCGGCAGTGGGCTTTGtCCTGGAATACACTTC |

| | |
|---|---|
| AND-apt-2 | GTCAAATGTCGCTCGtCAAGGGGTCTTGCCGAA |
| AND-input1 | GAAGTGTATTCCAGGctctctctctctct |
| AND-input2 | agagagagagagagCGAGCGACATaTGAC |
| OR-apt-1 | TTCGGCAAGAAGGGATGATGCGGCAGTGGGCTTTGtCCTGGAATACACTTC |
| OR-apt-2 | GTCAAATGTCGCTCGtCAAGGGGTCTTGCCGAA |
| OR-input1 | GAAGTGTATTCCAGGCGAGCGACATTTGAC |
| OR-input2 | GAAGTGTATTCCAGGctagagtgTTTTcactctagCGAGCGACATTTGAC |
| XOR-apt-1 | TTCGGCAAGAAGGGATGATGCGGCAGTGGGCTTTGtCCTGGAATACACTTC |
| XOR-apt-2 | GTCAAATGTCGCTCGtCAAGGGGTCTTGCCGAA |
| XOR-input1a | GAAGTGTATTCCAGGtctagagtgacgtgtccgtgt |
| XOR-input1b | gtcactctagtCGAGCGACATTTGAC |
| XOR-input2a | GAAGTGTATTCCAGGtacacggacacgtcactctag |
| XOR-input2b | gtgtccgtgttCGAGCGACATTTGAC |
| NAND-apt-1 | TTCGGCAAGAAGGGATGATGCGGCAGTGGGCTTTGCATAACTCCCTGGAAT |
| NAND-apt-2 | GTCGCTCGGAGTTATGCAAGGGGTCTTGCCGAA |
| NAND-input1 | ATTCCAGGGAGTctctctctctctct |
| NAND-input2 | agagagagagagagTATGCAAAGCC |
| NOR-apt-1 | TTCGGCAAGAAGGGATGATGCGGCAGTGGGCTTTGCATAACTCCCTGGAAT |
| NOR-apt-2 | GTCGCTCGGAGTTATGCAAGGGGTCTTGCCGAA |
| NOR-input1 | ATTCCAGGGAGTTATGCAAAGCCCA |
| NOR-input2 | ACCCCTTGCATAACTCCGAGCGAC |
| XNOR-apt-1 | TTCGGCAAGAAGGGATGATGCGGCAGTGGGCTTTGCATAACTCCCTGGAAT |
| XNOR-apt-2 | GTCGCTCGGAGTTATGCAAGGGGTCTTGCCGAA |
| XNOR-input1 | ATTCCAGGGAGTTATGCAaccgtcaaaaaaaaaaaaatgacggtAAGCCCA |
| XNOR-input2 | CCTaccgtcatttttttttttttgacggtTGCATAACTCCGAGCGAC |
| Ortho-apt-1-1 | TTCGGCAAGAAGGGATGATGCGGCAGTGGGCTTcatcAGGCTGTTCG |
| Ortho-apt-1-2 | CACCTCACTCgatgAGGGGTCTTGCCGAA |
| Ortho-M-1 | CGAACAGCCTGAGTGAGGTG |
| Ortho-apt-2-1 | TTCGGCAAGAAGGGATGATGCGGCAGTGGGCTTcatcTGCCAATCAG |
| Ortho-apt-2-2 | AGCGTTCCGAgatgAGGGGTCTTGCCGAA |
| Ortho-M-2 | CTGATTGGCATCGGAACGCT |
| Ortho-apt-3-1 | TTCGGCAAGAAGGGATGATGCGGCAGTGGGCTTcatcACCATAGAAT |
| Ortho-apt-3-2 | ACCAGCTCCAgatgAGGGGTCTTGCCGAA |
| Ortho-M-3 | ATTCTATGGTTGGAGCTGGT |
| Ortho-apt-4-1 | TTCGGCAAGAAGGGATGATGCGGCAGTGGGCTTcatcTAGACCAGAA |
| Ortho-apt-4-2 | GGATCACCCTgatgAGGGGTCTTGCCGAA |
| Ortho-M-4 | TTCTGGTCTAAGGGTGATCC |
| Ortho-apt-5-1 | TTCGGCAAGAAGGGATGATGCGGCAGTGGGCTTcatcTCGCCACTAA |
| Ortho-apt-5-2 | TTGATCCTGGgatgAGGGGTCTTGCCGAA |
| Ortho-M-5 | TTAGTGGCGACCAGGATCAA |
| Ortho-apt-6-1 | TTCGGCAAGAAGGGATGATGCGGCAGTGGGCTTcatcATTCCCGGAT |

| | |
|---|---|
| Ortho-apt-6-2 | CAGCTTCCTCgatgAGGGGTCTTGCCGAA |
| Ortho-M-6 | ATCCGGGAATGAGGAAGCTG |
| Ortho-apt-7-1 | TTCGGCAAGAAGGGATGATGCGGCAGTGGGCTTcatcATGACCCTGA |
| Ortho-apt-7-2 | TAACCACATAgatgAGGGGTCTTGCCGAA |
| Ortho-M-7 | TCAGGGTCATTATGTGGTTA |
| Ortho-apt-8-1 | TTCGGCAAGAAGGGATGATGCGGCAGTGGGCTTcatcCCTAAGATGA |
| Ortho-apt-8-2 | TTGCCATGCCgatgAGGGGTCTTGCCGAA |
| Ortho-M-8 | TCATCTTAGGGGCATGGCAA |
| Ortho-apt-9-1 | TTCGGCAAGAAGGGATGATGCGGCAGTGGGCTTcatcGTACCTTCAC |
| Ortho-apt-9-2 | GTCACAGTGGgatgAGGGGTCTTGCCGAA |
| Ortho-M-9 | GTGAAGGTACCCACTGTGAC |
| Ortho-apt-10-1 | TTCGGCAAGAAGGGATGATGCGGCAGTGGGCTTcatcGTCCGCGCAT |
| Ortho-apt-10-2 | CATGTGTCATgatgAGGGGTCTTGCCGAA |
| Ortho-M-10 | ATGCGCGGACATGACACATG |
| BUF-apt-1 | TTCGGCAAGAAGGGATGATGCGGCAGTGGGCTTcatcTGCCAATCAG |
| BUF-apt-2 | AGCGTTCCGAgatgAGGGGTCTTGCCGAA |
| BUF-apt-3 | GTCAAATGTCAGGGATGATGCGGCAGTGGGCTTcatcTAGACCAGAA |
| BUF-apt-4 | GGATCACCCTgatgAGGGGGACATTTGAC |
| BUF-apt-5 | CCTTGACTGTAGGGATGATGCGGCAGTGGGCTTcatcATGACCCTGA |
| BUF-apt-6 | TAACCACATAgatgAGGGGACAGTCAAGG |
| BUF-input-1 | CTGATTGGCATCGGAACGCT |
| BUF-input-2 | TTCTGGTCTAAGGGTGATCC |
| BUF-input-3 | TCAGGGTCATTATGTGGTTA |
| Major-apt-1 | GTCAAATGTCGCTCGtGAAGGGATGATGCGGCAGTGGGCTTTGtCACAAGCAGTTCTGC |
| Major-apt-2 | CCGCCGAGGATAAACtCAAGGGGTCtCCTGGAATACACTTC |
| Major-apt-3 | CCGCCGAGGATAAACtGAAGGGATGATGCGGCAGTGGGCTTTGtTTGCCCTGAGCAACG |
| Major-apt-4 | CCTTGACTCTTTGCAtCAAGGGGTCtCACAAGCAGTTCTGC |
| Major-apt-5 | CCTTGACTCTTTGCAtGAAGGGATGATGCGGCAGTGGGCTTTGtCCTGGAATACACTTC |
| Major-apt-6 | GTCAAATGTCGCTCGttCAAGGGGTCttTTGCCCTGAGCAACG |
| Major-input-1 | GAAGTGTATTCCAGGCGAGCGACATTTGAC |
| Major-input-2 | GCAGAACTGCTTGTGGTTTATCCTCGGCGG |
| Major-input-3 | CGTTGCTCAGGGCAATGCAAAGAGTCAAGG |

#### 10-HB origami sequences:

| Strand name | Sequence (5' to 3') |
|---|---|
| 10HB-core1 | TTTTGAGGCGAGTAGCGGATTGACCGTAGACTTGTGCAACCG |
| 10HB-core2 | AGATGAAGTTATATTAAATTTAGAACGGGTATTAAGGAATCA |
| 10HB-core3 | TGAGGGGCGTTTTTCTGGAGGTGTCCAGCCATATAATCATAC |
| 10HB-core4 | CAGCCAGTAAAGTTGCACTCAATCCGCCTTTTAAAAGCCTCA |
| 10HB-core5 | AAAGCGCCCCGTAACTGTTGCCCTGCGGCTTAGAGATGACCC |
| 10HB-core6 | GCGATCGCCGCCAGGTCATAAACATCCCAAGCATATCATAGT |
| 10HB-core7 | AGGGGGAGACGCAGACGGCATCAGATGCAAATTGTACGTTAG |
| 10HB-core8 | GTTTTCCAGACTTTGCCCCCTGCATCAGGAGCTCGCTTTCAA |
| 10HB-core9 | GCTTTCACGTACAGTGTGCACTCTGTGGCGCGTCCAAAGGAA |
| 10HB-core10 | ATATCAAAAAGCATTTCACGGTCGGAGCTAAAGATAAACAAA |
| 10HB-core11 | CAGTTGAGCAACAGCGGCGAACGTGGCGTTTTTTGCAGTCTC |
| 10HB-core12 | GAGCACTGAAGATACGGGCGCTAGGGCGTCAGGGCTTTGATG |
| 10HB-core13 | CATTTGACCTAAAATAACCACCACACCCAAAGAACTCAGTGC |
| 10HB-core14 | CAACTCGATTTTTGCGTACTATGGTTGCCAACATGATAAATC |
| 10HB-core15 | AAGTTTGCCCTTCTTCGTTAGAATCAGACAACAGTCGTCGCT |
| 10HB-core16 | CAGAAGGCGACCAGGGGATTTTAGACAGCATATGCAGCGATA |
| 10HB-core17 | GTTGGTGTCCTCATGCAGCACCGTCGGTAGTTTGAGGCATCA |
| 10HB-core18 | ATGGCAAGTCTGAATTTTTATAATCAGTAAACACCATTTATC |
| 10HB-core19 | TCTGAATGGAAAAATCACGCAAATTAACTGAAATACCGGCTT |
| 10HB-core20 | CGCGGTCACGACGACAGTCAAATCACCATCAATATGCCAGTT |
| 10HB-core21 | TGAAGGGCTTTCCGAATGTGTAGGTAAAGATTCAACGCACTC |
| 10HB-core22 | AAAAAATCATTCGCAATTTTTAGAACCCTCATATAACCAGGC |
| 10HB-core23 | TCATTTGGTGCGGGTACAACGCCTGTAGCATTCCTGGGAAGG |
| 10HB-core24 | AAAAAGATGTGCTGAACCCATGTACCGTAACACTGTGGCGAA |
| 10HB-core25 | GAGAGATCAGTCACAGAGCCACCACCCTCATTTTCCGCCAGG |
| 10HB-core26 | GGCGAAAGAGGTGGACCGCCACCCTCAGAACCGCCGTGCCAA |
| 10HB-core27 | AAAATCTACCCTCAATAGCCCGGAATAGGTGTATCACCTCAA |
| 10HB-core28 | ACCGCCTAAGGAATAGTACCAGGCGGATAAGTGCCAAATCAA |
| 10HB-core29 | ACCAGCAAACAACTAGACTCCTCAAGAGAAGGATTTCTTTAG |
| 10HB-core30 | TGATAGCGGATTTATCGGAACCTATTATTCTGAAAGATAATA |
| 10HB-core31 | AGACAATTATTAAATTTGAATTACCTTTTTTAAACAATTCGA |
| 10HB-core32 | GATAGAAAGTAACACAAACATCAAGAAAACAAAATATTTTAA |
| 10HB-core33 | AGTCACAAGCGGAAAATTATTCATTTCAATTACCTAAACCAC |
| 10HB-core34 | CTCAATCTTCATCACTGATTGCTTTGAATACCAAGTCAGATG |
| 10HB-core35 | TGCAACAAATGGAATAACAGTACCTTTTACATCGGTTATACT |
| 10HB-core36 | GAGCACATAGATGGATAAATTAATGCCGGAGAGGGAGGTCAC |
| 10HB-core37 | CAAGAATGCCAACGAACGGAACGTGCCGATGGGATTAGCTAT |
| 10HB-core38 | TTCTTTGTCTGACCAACTATAGAGCCAGCAAAATCGGAGGGA |
| 10HB-core39 | ATTCTGCAGTAGCACACCAACCTAAAACCGCGACCCGGAATC |
| 10HB-core40 | ACGGTGTATTAGCAGTTTCCATTAAACGGGTCAATGGGTAAT |
| 10HB-core41 | ATAATGCATCGGTTGGCTACAGAGGCTTACAGATGCAAAATA |

| | |
|---|---|
| 10HB-core42 | GGGTGCCGGGAGAAACCCTCAGCAGCGACTTCATCCAACACT |
| 10HB-core43 | TCCACACAAAGTTTTGAGGCTTGCAGGGCCAAATCCAGATAC |
| 10HB-core44 | GTCATAGGTATGGGGACAACAACCATCGTGCCCTGGATTCAT |
| 10HB-core45 | AGTTGAGTGAGAATGAATTTCTTAAACAGAGATGGTAATAAA |
| 10HB-core46 | CCAGCACTTTTTCAGGAGCCTTTAATTGTGCGATTGCTCATT |
| 10HB-core47 | TAAAGCATTAAAGCAGGTCAGACGATTGATTGAGTTAACTGA |
| 10HB-core48 | TGAACCACAGTAAGCCACCACCAGAGCCGCAATAGGAGAATA |
| 10HB-core49 | CAAAGGGTGGTAATCCGCCACCCTCAGAAGATAGCTTTTTGT |
| 10HB-core50 | GCATTTTCCCGTATCCACCACCGGAACCAATAATATAAACAG |
| 10HB-core51 | AATCGCCAATAACCGCGTTTGCCATCTTCCTTATTCTAACGA |
| 10HB-core52 | TCGCAAATGGGGCGCTCATCTTTGACCCAACGGAGTAAACAG |
| 10HB-core53 | TACCAGTCCTTAGAAGCGCGTTTTCATCTAAAGGTATTTTGC |
| 10HB-core54 | TAGAAAAGCTGAGATCAGTAGCGACAGAATAAGTTACTTGCG |
| 10HB-core55 | AATAAGGGAGAGACCCGGAAACGTCACCTACCAGCTCCGGTA |
| 10HB-core56 | TTCATCTATTAGTATATCCAGAACAATACATATCATAACGTC |
| 10HB-core57 | AGGCAAGCCGGAGACAGTATCGGCCTCAGCCGTTCGGGGTCA |
| 10HB-core58 | GAGCATACCTGAGTGCACCGCTTCTGGTGCGGCCTGTTGCGG |
| 10HB-core59 | TGTAATAGGATAAACATTCAGGCTGCGCTGTACATTGGGTAA |
| 10HB-core60 | TAGCGTATACAAACCCTCTTCGCTATTAACTTAAAGGTGTGT |
| 10HB-core61 | TAAATGACAATAGGCAAGGCGATTAAGTGATAGCTCCTGCAG |
| 10HB-core62 | CAGTTTCCACCCTCGACGTTGTAAAACGGTCCCGGAGCGCAG |
| 10HB-core63 | TTGCGAAGTTTAGTAGCCGCCACGGGAACACCGGAGCCAGAA |
| 10HB-core64 | CAAATAATATAAGTATCAATATCTGGTCCCAGCAGTTAGAGC |
| 10HB-core65 | TGAATTTTTTGCTCTGAGGAAGGTTATCCGGTCAGAAGGGAA |
| 10HB-core66 | ATACAGGGAGGCTGAATAGATTAGAGCCATACCGAGTGTAGC |
| 10HB-core67 | CTTGAGTGCCTATTGAAGTATTAGACTTTTTAATGTTAATGC |
| 10HB-core68 | AATATATAATTTCATCCTTTGCCCGAACAGAATACAGCACGT |
| 10HB-core69 | ATTAATTGATGAAATTATCATTTTGCGGTTCTGGCCTAAACA |
| 10HB-core70 | GCTTAGAAGAGGCGTTATCATCATATTCTTGGCAGACGCCAG |
| 10HB-core71 | AAAATCAGATTCGCATATAATCCTGATTCCTACATCCGAGTA |
| 10HB-core72 | AGGTTGGTATACAGGGGTTAGAACCTACTTACCGCGCAATAC |
| 10HB-core73 | ATTCTACCTAGCTGGCGCATCGTAACCGGCAGCCTTCCCACG |
| 10HB-core74 | TGCCACTTAGCCGGACTGGATAGCGTCCGAAAGACATTCCCA |
| 10HB-core75 | TTTTTCAACTGACCCAAAAGAAGTTTTGAACCAGACTAAAGT |
| 10HB-core76 | ACGAGGGCAGGCGCTTACCAGACGACGATACCTTTGCTGAAT |
| 10HB-core77 | TTTGCGGCAAGAACAATTACGAGGCATACTAACTCAAGCCTG |
| 10HB-core78 | ATATATTGCTCATTGAATACCACATTCAGTCGGGATCACAAT |
| 10HB-core79 | GTTGCGCAACGAGTCAACATTATTACAGACGCGCGAATCATG |
| 10HB-core80 | CTTGCTTTAATCATGTTGGGAAGAAAAATTTTTCTTCCTCAC |
| 10HB-core81 | AAAAGGCTCAGAGAGTCAGAGGGTAATTTCACCGCGTTTCTG |
| 10HB-core82 | CAGGAGGAGAGCAAAAGCGCATTAGACGTGGTTTGGGTGCCG |
| 10HB-core83 | AGCCGCCCCCTTTTGAAAATAGCAGCCTCGAAATCCACTACG |
| 10HB-core84 | CGCCACCAGAAGGACCAATCCAAATAAGGATAGGGCCAACGT |

| | |
|---|---|
| 10HB-core85 | CACCGGAAGAACTGCCTAATTTGCCAGTTAAGAGAGGCAGAG |
| 10HB-core86 | TAGCCCCCAAACGTTTATCCTGAATCTTCCAGACGAATTGAG |
| 10HB-core87 | TAGCGTCGAAACGCCTTAAATCAAGATTGCGCCTGAAATTCT |
| 10HB-core88 | TAGCAGCCAATAGAGGCGTTTTAGCGAATAATATCTAATTAC |
| 10HB-core89 | TACCATTGGCGACAAAATCAGATATAGATAATCGGTGTGATA |
| 10HB-core90 | AATACACCTGATAATTGAATCCCCCTCAAGAAGCAATACATT |
| 10HB-core91 | AGGTAAATATTGACGGAATTATGTAAATGCTGATGAGTTAAT |
| 10HB-core92 | AGTAAAATTTAATTCGAGCTTCAAAGCGCCAGAGGCATAAGG |
| 10HB-core93 | GCGAGAGCAACAGGTCAGGATTAGAGAGTAAAAACAACGGTG |
| 10HB-core94 | ATCATAAATAAGAGGTCATATGAGTGAGGTAAGAGAAGAGTA |
| 10HB-core95 | ATAACGCGCGCTCACTGCCCGCTTTCCAACTAATGAACGTAA |
| 10HB-core96 | CAGTTGAGCTGCATTAATGAATCGGCCAGTAGAAAACGAGAA |
| 10HB-core97 | ACGAACTGCGTATTGGGCGCCAGGGTGGTCTACGTTTTAATT |
| 10HB-core98 | ATACCAGCGGGCAACAGCTGATTGCCCTGAAACTGTTAAGGC |
| 10HB-core99 | ACACCCTGTTGCAGCAAGCGGTCCACGCGGAGAATTAAGCCC |
| 10HB-core100 | ACATAAAAATCCTGTTTGATGGTGGTTCTTACAGACTATCTT |
| 10HB-core101 | TTCAGAATTTACCCTGACTATTATAGTCAATGCTTATTTGTA |
| 10HB-core102 | TTAACGTTAAATCAAAAGAATAGCCCGAAAACGATCGAACAA |
| 10HB-core103 | CCATATTAGTTTGGAACAAAGCCAGTAATACAAAAACGGAAT |
| 10HB-core104 | GCGTCTTCAAAAGGTAAAGTAATTCTGTACCAACGACGCAGT |
| 10HB-core105 | ACCCAGCCATGTTCAGCTAATGCAGAACAGTTGCTGGCAACA |
| 10HB-core106 | GGAGGTTATAAGTCCTGAACAAGAAAAACCTCCCGTATTTTG |
| 10HB-core107 | TTCTAAGGAGCATGTAGAAACCAATCAAAGGCTTAGCCAAAG |
| 10HB-core108 | TTACCGCGCCCAATCATTCCAAATGGTTCGTTGTACAGCCAT |
| 10HB-core109 | GTCATAAAAAAAGATTAAGAGGAAGCCCAATACTGTGCTCCA |
| 10HB-core110 | TCATCGCTAAAACACGAGCTGAAAAGGTCCATTAGAAGCGGA |
| 10HB-core111 | TTGCATCATATTCAATTGTGTCGAAATCGAAAGAGGCAAAAG |
| 10HB-edge-L1 | ACAACCCGTCGGATTCAA |
| 10HB-edge-L2 | ATTCCGTGGGAACAAACG |
| 10HB-edge-L3 | AGAACGTCAGCGTGGTAC |
| 10HB-edge-L4 | GAGCTGGTCTGGTCAGCA |
| 10HB-edge-L5 | TGGTCAATAACCTGTTCG |
| 10HB-edge-L6 | CGTAGCTATATTTTCATT |
| 10HB-edge-L7 | CCAGCGATTATACCAATC |
| 10HB-edge-L8 | TAGCGCGAAACAAAGTAC |
| 10HB-edge-L9 | AACGAGAATGACCATAAG |
| 10HB-edge-L10 | CAAATCAAAAATCAGGTC |
| 10HB-edge-R1 | AGAAATTGCGTAGATTTTCAGGTT |
| 10HB-edge-R2 | AAATTATTTGCACGTAAAACAGAA |
| 10HB-edge-R3 | TCAAACTATCGGCCTTGCTGGTAA |
| 10HB-edge-R4 | ATAACATCACTTGCCTGAGTAGAA |

| | |
|---|---|
| 10HB-edge-R5 | AGAAAACTTTTTCAAATATATTTT |
| 10HB-edge-R6 | CAAATCCAATCGCAAGACAAAGAA |
| 10HB-edge-R7 | ACCGTCACCGACTTGAGCCATTTG |
| 10HB-edge-R8 | GGAAATTATTCATTAAAGGTGAAT |
| 10HB-edge-R9 | CAAGCCGTTTTTATTTTCATCGTA |
| 10HB-edge-R10 | ACCAAGTACCGCACTCATCGAGAA |
| 10HB-edge-L1-tttt | ACAACCCGTCGGATTCTTTT |
| 10HB-edge-L2-tttt | TTTTTCCGTGGGAACAAACG |
| 10HB-edge-L3-tttt | AGAACGTCAGCGTGGTTTTT |
| 10HB-edge-L4-tttt | TTTTGCTGGTCTGGTCAGCA |
| 10HB-edge-L5-tttt | TGGTCAATAACCTGTTTTTT |
| 10HB-edge-L6-tttt | TTTTTAGCTATATTTTCATT |
| 10HB-edge-L7-tttt | CCAGCGATTATACCAATTTT |
| 10HB-edge-L8-tttt | TTTTGCGCGAAACAAAGTAC |
| 10HB-edge-L9-tttt | AACGAGAATGACCATATTTT |
| 10HB-edge-L10-tttt | TTTTAATCAAAAATCAGGTC |
| 10HB-edge-R1-tttt | TTTTAAAGAAATTGCGTAGATTTTCAGGTT |
| 10HB-edge-R2-tttt | AAATTATTTGCACGTAAAACAGAAATTTTT |
| 10HB-edge-R3-tttt | TTTTACTCAAACTATCGGCCTTGCTGGTAA |
| 10HB-edge-R4-tttt | ATAACATCACTTGCCTGAGTAGAAGATTTT |
| 10HB-edge-R5-tttt | TTTTCGAGAAAACTTTTTCAAATATATTTT |
| 10HB-edge-R6-tttt | CAAATCCAATCGCAAGACAAAGAACGTTTT |
| 10HB-edge-R7-tttt | TTTTTCACCGTCACCGACTTGAGCCATTTG |

| | |
|---|---|
| 10HB-edge-R8-tttt | GGAAATTATTCATTAAAGGTGAATTATTTT |
| 10HB-edge-R9-tttt | TTTTAGCAAGCCGTTTTTATTTTCATCGTA |
| 10HB-edge-R10-tttt | ACCAAGTACCGCACTCATCGAGAACATTTT |
| 10HB-edge-L1-sticky4nt | ACAACCCGTCGGATTCAAAG |
| 10HB-edge-L2-sticky4nt | AAATTCCGTGGGAACAAACG |
| 10HB-edge-L3-sticky4nt | AGAACGTCAGCGTGGTACTC |
| 10HB-edge-L4-sticky4nt | AAGAGCTGGTCTGGTCAGCA |
| 10HB-edge-L5-sticky4nt | TGGTCAATAACCTGTTCGAG |
| 10HB-edge-L6-sticky4nt | AACGTAGCTATATTTTCATT |
| 10HB-edge-L7-sticky4nt | CCAGCGATTATACCAATCAC |
| 10HB-edge-L8-sticky4nt | ATTAGCGCGAAACAAAGTAC |
| 10HB-edge-L9-sticky4nt | AACGAGAATGACCATAAGCA |
| 10HB-edge-L10-sticky4nt | AACAAATCAAAAATCAGGTC |
| 10HB-edge-R1-sticky4nt | AAATTGCGTAGATTTTCAGGTT |
| 10HB-edge-R2-sticky4nt | AAATTATTTGCACGTAAAACAG |
| 10HB-edge-R3-sticky4nt | AAACTATCGGCCTTGCTGGTAA |
| 10HB-edge-R4-sticky4nt | ATAACATCACTTGCCTGAGTAG |
| 10HB-edge-R5-sticky4nt | AAAACTTTTTCAAATATATTTT |
| 10HB-edge-R6-sticky4nt | CAAATCCAATCGCAAGACAAAG |
| 10HB-edge-R7-sticky4nt | CGTCACCGACTTGAGCCATTTG |
| 10HB-edge-R8-sticky4nt | GGAAATTATTCATTAAAGGTGA |

| | |
|---|---|
| 10HB-edge-R9-sticky4nt | AGCCGTTTTTATTTTCATCGTA |
| 10HB-edge-R10-sticky4nt | ACCAAGTACCGCACTCATCGAG |
| helix-9-1-unmodified | TGTTACTACGAAGGTTAACATCCAATAAACAGTTGTTCAAAT |
| helix-9-2-unmodified | GAACCGATGAGGAAAAATTAAGCAATAATATGCAACCGGAAG |
| helix-9-3-unmodified | TACAGACTAGCAACGTACCAAAAACATTCTTAATTAATTGCT |
| helix-9-4-unmodified | ATCTTGAGATCGTCGCCACAGACAGCCCAAGTGTAACATTAA |
| helix-9-5-unmodified | CAAAGCTCGGTCGCTGTCGTCTTTCCAGTATCCGCAACCTGT |
| helix-9-6-unmodified | ACACCAGCGACAATATTTTGCTAAACAAAATTCGTGGGAGAG |
| helix-9-7-unmodified | TCAACTTTCGAGGTAGAAAGGAACAACTGTGAGCCTTTCACC |
| helix-9-8-unmodified | GCTAATATCCAAAACGTTGAAAATCTCCACCGGGGCTGGCCC |
| helix-9-9-unmodified | AATAATATTGAGGCCAGAATGGAAAGCGGGGTCGACCCCAGC |
| helix-9-10-unmodified | ACCGAAGACCAGAACGTCATACATGGCTGATGGCCGGCAAAA |
| helix-9-11-unmodified | AGTTACCCTCAGAAAAGTTTTAACGGGGGTGGACTTTGAGTG |
| helix-9-12-unmodified | ACCCAAAACCAGAGAATGGAAACAGTACTAATTTAATATAAA |
| helix-9-13-unmodified | ATGTTAGCTTATTATTGCTTCTGTAAATAGGGCTTACGACAA |
| helix-9-14-unmodified | TATAAAAAGACTGTATCCTTGAAAACATGTTATACTTTATCA |
| helix-9-15-unmodified | TCACAATACCGTAAAGAGTCAATAGTGAGGAATCACCATCCT |
| helix-9-16-unmodified | ACAAAAGAGCAAGGTACCTTTTTAACCTCCGACCGCTGTCTT |
| helix-9-17-unmodified | ATCGCGTTGTTTAGAACGAGGCGCAGACGGTAAAATACGTAA |
| helix-9-18-unmodified | CAAACTCGCTTTTGAACTTTGAAAGAGGTGAGGACTAAAGAC |
| helix-9-19-unmodified | CCTTTTGCCCTCGTATAGGCTGGCTGACAAGACAGCATCGGA |

| | |
|---|---|
| helix-9-20-unmodified | TTGCGTTCAAAAGGCGGATATTCATTACAGTTAAAGGCCGCT |
| helix-9-21-unmodified | CGTGCCAGATTTAGCAGTGAATAAGGCTCCCACGCATAACCG |
| helix-9-22-unmodified | GCGGTTTAACGGAAAGTAAATTGGGCTTGCTTGATACCGATA |
| helix-9-23-unmodified | AGTGAGATCAGGACTGTGAATTACCTTATATCGGTTTATCAG |
| helix-9-24-unmodified | TGAGAGAGAACAAAGATAACCCACAAGAGCCTTGATATTCAA |
| helix-9-25-unmodified | AGGCGAAAACAGGGGAAACAATGAAATAGCCGCCAGCATTGA |
| helix-9-26-unmodified | TCCCTTACAAAAATTAAGAAAAGTAAGCGCCACCACCCTCAG |
| helix-9-27-unmodified | TTGTTCCATTTATCAACCGAGGAAACGCGCCTCCCTCAGAGC |
| helix-9-28-unmodified | GTACCGATCCAGAGGCATGATTAAGACTTTCATAATCAAAAT |
| helix-9-29-unmodified | TAAACAATACAATTAGAAAATACATACAGGCATTTTCGGTCA |
| helix-9-30-unmodified | ACAATAGTTGAAGCAAAGACACCACGGAATCAAGTTTGCCTT |
| helix-9-31-unmodified | AATTTACAACGCGAAAATTCATATGGTTAATGAAACCATCGA |
| helix-9-32-unmodified | TCCTTATAGCAAGCTTCAACCGATTGAGACCAGTAGCACCAT |
| helix-0-1-unmodified | AAAGAGTCTGTCCACGCTCATGGAAATAGTTTGGAGAGAAAC |
| helix-0-2-unmodified | AATCCTGAGAAGTGATGGATTATTTACACTGATTATTACAAA |
| helix-0-3-unmodified | GGAGGCCGATTAAATAATAAAAGGGACAAACAAAGGAGCAAA |
| helix-0-4-unmodified | ATAACGTGCTTTCCGACCTGAAAGCGTAGTTATTATAATTAC |
| helix-0-5-unmodified | GCCGCTACAGGGCGAATGGCTATTAGTCTACAAACAGTTAAT |
| helix-0-6-unmodified | GGTCACGCTGCGCGCATCGCCATTAAAAGTCAATACATGAAA |
| helix-0-7-unmodified | GAAAGCGAAAGGAGAAACAGAGGTGAGGTAAAATAAGGATTA |
| helix-0-8-unmodified | TTGACGGGGAAAGCTGCCACGCTGAGAGAGTTGGCGTCGAGA |

| | |
|---|---|
| helix-0-9-unmodified | TGCGGCGGGCCGTTCACCTTGCTAACCTCGGATGAACCGTAC |
| helix-0-10-unmodified | TGTCACTGCGCGCCCGCCATGTTTACCAACGGCCAACCCTCA |
| helix-0-11-unmodified | CCAGCGGTGCCGGTCTCCGTGGTGAAGGTGGGTAAAGGGATA |
| helix-0-12-unmodified | TCAGCAAATCGTTAAAACAGCGGATCAACGCCAGCAGTTTCG |
| helix-0-13-unmodified | AGGTTTCTTTGCTCCAGTTGGGCGGTTGAACTGTTTTATTTC |
| helix-0-14-unmodified | TATGAGCCGGGTCAAAAAAGCCGCACAGGCCGGAATTTTAAA |
| helix-0-15-unmodified | TTGCAGGCGCTTTCAAACGATGCTGATTGGAAGATAAGGGTG |
| helix-0-16-unmodified | CAACCAGCTTACGGTCGTCTCGTCGCTGTGCATCTGATATTC |
| helix-0-17-unmodified | AATAACGTAGGTCTCGTTAAATAAGAATGAGGCCATTTGACG |
| helix-0-18-unmodified | ATCGCGCTTAAGACAGCCTGTTTAGTATGAACGGTATTCACC |
| helix-0-19-unmodified | AGAAGATAATTTTCATAAAGCCAACGCTGCGGGAGCAACAGA |
| helix-0-20-unmodified | ATTTAACGTGAGTGATATTTAACAACGCTTTGACGGTGGCAC |
| helix-0-21-unmodified | GCCCCCTAACAGTGCGGAGTCCACTATTGCCGCGCCGCGAAC |
| helix-0-22-unmodified | GTATTAAAGTGTACCGAAAAACCGTCTACTGGCAAACGAACC |
| helix-0-23-unmodified | GCGGGGTACCGTTCTCACCCAAATCAAGAGAAAGGTATTAAC |
| helix-0-24-unmodified | GGGTTGAATCCTCACTAAATCGGAACCCCCCCGATCAAATGA |
| helix-0-25-unmodified | TCAGGAGTAATAATGCGTGCCTGTTCTTTGCTGCGAACAATC |
| helix-0-26-unmodified | GAACCGCAGCGGAGGATCCCCGGGTACCACGATCCAATTTGT |
| helix-0-27-unmodified | GCAAGCCATTTTCTCTGTTTCCTGTGTGCGGGTTACTCACGG |
| helix-0-28-unmodified | TCACCAGACGATCTAACATACGAGCCGGTTACACTTTTCTGC |
| helix-0-29-unmodified | AACGCAACTTTTGCTATTTTGCGGATGGCTGGTAACGACATA |

| | |
|---|---|
| helix-0-30-unmodified | TGCAATGAAGCTAATGTAGCTCAACATGGGGCGCGTTAGTGA |
| helix-0-31-unmodified | AGAAAGGGCAAAGACTGGAAGTTTCATTCATCAGCCGGCAAA |
| helix-0-32-unmodified | AACCGTTTAATAGTGAACGAGTAGATTTGGTGCCACCGGCCA |
| helix-9-1-apt | TGTTACTACGAAGGTTAACATCCAATAAACAGTTGTTCAAATtAAGAAGGGATGATGCGGCAGTGGGCTTcatcAGGCTGTTCG |
| helix-9-2-apt | GAACCGATGAGGAAAAATTAAGCAATAATATGCAACCGGAAGtAAGAAGGGATGATGCGGCAGTGGGCTTcatcAGGCTGTTCG |
| helix-9-3-apt | TACAGACTAGCAACGTACCAAAAACATTCTTAATTAATTGCTtAAGAAGGGATGATGCGGCAGTGGGCTTcatcAGGCTGTTCG |
| helix-9-4-apt | ATCTTGAGATCGTCGCCACAGACAGCCCAAGTGTAACATTAAtAAGAAGGGATGATGCGGCAGTGGGCTTcatcAGGCTGTTCG |
| helix-9-5-apt | CAAAGCTCGGTCGCTGTCGTCTTTCCAGTATCCGCAACCTGTtAAGAAGGGATGATGCGGCAGTGGGCTTcatcAGGCTGTTCG |
| helix-9-6-apt | ACACCAGCGACAATATTTTGCTAAACAAAATTCGTGGGAGAGtAAGAAGGGATGATGCGGCAGTGGGCTTcatcAGGCTGTTCG |
| helix-9-7-apt | TCAACTTTCGAGGTAGAAAGGAACAACTGTGAGCCTTTCACCtAAGAAGGGATGATGCGGCAGTGGGCTTcatcAGGCTGTTCG |
| helix-9-8-apt | GCTAATATCCAAAACGTTGAAAATCTCCACCGGGGCTGGCCCtAAGAAGGGATGATGCGGCAGTGGGCTTcatcAGGCTGTTCG |
| helix-9-9-apt | AATAATATTGAGGCCAGAATGGAAAGCGGGGTCGACCCCAGCtAAGAAGGGATGATGCGGCAGTGGGCTTcatcAGGCTGTTCG |
| helix-9-10-apt | ACCGAAGACCAGAACGTCATACATGGCTGATGGCCGGCAAAAtAAGAAGGGATGATGCGGCAGTGGGCTTcatcAGGCTGTTCG |
| helix-9-11-apt | AGTTACCCTCAGAAAAGTTTTAACGGGGGTGGACTTTGAGTGtAAGAAGGGATGATGCGGCAGTGGGCTTcatcAGGCTGTTCG |
| helix-9-12-apt | ACCCAAAACCAGAGAATGGAAACAGTACTAATTTAATATAAAtAAGAAGGGATGATGCGGCAGTGGGCTTcatcAGGCTGTTCG |
| helix-9-13-apt | ATGTTAGCTTATTATTGCTTCTGTAAATAGGGCTTACGACAAtAAGAAGGGATGATGCGGCAGTGGGCTTcatcAGGCTGTTCG |
| helix-9-14-apt | TATAAAAAGACTGTATCCTTGAAAACATGTTATACTTTATCAtAAGAAGGGATGATGCGGCAGTGGGCTTcatcAGGCTGTTCG |
| helix-9-15-apt | TCACAATACCGTAAAGAGTCAATAGTGAGGAATCACCATCCTtAAGAAGGGATGATGCGGCAGTGGGCTTcatcAGGCTGTTCG |
| helix-9-16-apt | ACAAAAGAGCAAGGTACCTTTTTAACCTCCGACCGCTGTCTTtAAGAAGGGATGATGCGGCAGTGGGCTTcatcAGGCTGTTCG |
| helix-9-17-apt | CACCTCACTCgatgAGGGGTCTTtATCGCGTTGTTTAGAACGAGGCGCAGACGGTAAAATACGTAA |
| helix-9-18-apt | CACCTCACTCgatgAGGGGTCTTtCAAACTCGCTTTTGAACTTTGAAAGAGGTGAGGACTAAAGAC |

| | |
|---|---|
| helix-9-19-apt | CACCTCACTCgatgAGGGGTCTTtCCTTTTGCCCTCGTATAGGCTGGCTGACAAGACAGCATCGGA |
| helix-9-20-apt | CACCTCACTCgatgAGGGGTCTTtTTGCGTTCAAAAGGCGGATATTCATTACAGTTAAAGGCCGCT |
| helix-9-21-apt | CACCTCACTCgatgAGGGGTCTTtCGTGCCAGATTTAGCAGTGAATAAGGCTCCCACGCATAACCG |
| helix-9-22-apt | CACCTCACTCgatgAGGGGTCTTtGCGGTTTAACGGAAAGTAAATTGGGCTTGCTTGATACCGATA |
| helix-9-23-apt | CACCTCACTCgatgAGGGGTCTTtAGTGAGATCAGGACTGTGAATTACCTTATATCGGTTTATCAG |
| helix-9-24-apt | CACCTCACTCgatgAGGGGTCTTtTGAGAGAGAACAAAGATAACCCACAAGAGCCTTGATATTCAA |
| helix-9-25-apt | CACCTCACTCgatgAGGGGTCTTtAGGCGAAAACAGGGGAAACAATGAAATAGCCGCCAGCATTGA |
| helix-9-26-apt | CACCTCACTCgatgAGGGGTCTTtTCCCTTACAAAAATTAAGAAAAGTAAGCGCCACCACCCTCAG |
| helix-9-27-apt | CACCTCACTCgatgAGGGGTCTTtTTGTTCCATTTATCAACCGAGGAAACGCGCCTCCCTCAGAGC |
| helix-9-28-apt | CACCTCACTCgatgAGGGGTCTTtGTACCGATCCAGAGGCATGATTAAGACTTTCATAATCAAAAT |
| helix-9-29-apt | CACCTCACTCgatgAGGGGTCTTtTAAACAATACAATTAGAAAATACATACAGGCATTTTCGGTCA |
| helix-9-30-apt | CACCTCACTCgatgAGGGGTCTTtACAATAGTTGAAGCAAAGACACCACGGAATCAAGTTTGCCTT |
| helix-9-31-apt | CACCTCACTCgatgAGGGGTCTTtAATTTACAACGCGAAAATTCATATGGTTAATGAAACCATCGA |
| helix-9-32-apt | CACCTCACTCgatgAGGGGTCTTtTCCTTATAGCAAGCTTCAACCGATTGAGACCAGTAGCACCAT |
| helix-0-1-apt | AAAGAGTCTGTCCACGCTCATGGAAATAGTTTGGAGAGAAACtAAGAAGGGATGATGCGGCAGTGGGCTTcatcAGGCTGTTCG |
| helix-0-2-apt | AATCCTGAGAAGTGATGGATTATTTACACTGATTATTACAAAtAAGAAGGGATGATGCGGCAGTGGGCTTcatcAGGCTGTTCG |
| helix-0-3-apt | GGAGGCCGATTAAATAATAAAAGGGACAAACAAAGGAGCAAAtAAGAAGGGATGATGCGGCAGTGGGCTTcatcAGGCTGTTCG |
| helix-0-4-apt | ATAACGTGCTTTCCGACCTGAAAGCGTAGTTATTATAATTACtAAGAAGGGATGATGCGGCAGTGGGCTTcatcAGGCTGTTCG |
| helix-0-5-apt | GCCGCTACAGGGCGAATGGCTATTAGTCTACAAACAGTTAATtAAGAAGGGATGATGCGGCAGTGGGCTTcatcAGGCTGTTCG |
| helix-0-6-apt | GGTCACGCTGCGCGCATCGCCATTAAAAGTCAATACATGAAAtAAGAAGGGATGATGCGGCAGTGGGCTTcatcAGGCTGTTCG |
| helix-0-7-apt | GAAAGCGAAAGGAGAAACAGAGGTGAGGTAAAATAAGGATTAtAAGAAGGGATGATGCGGCAGTGGGCTTcatcAGGCTGTTCG |

| | |
|---|---|
| helix-0-8-apt | TTGACGGGGAAAGCTGCCACGCTGAGAGAGTTGGCGTCGAGAtAAGAAGGGATGATG CGGCAGTGGGCTTcatcAGGCTGTTCG |
| helix-0-9-apt | TGCGGCGGGCCGTTCACCTTGCTAACCTCGGATGAACCGTACtAAGAAGGGATGATG CGGCAGTGGGCTTcatcAGGCTGTTCG |
| helix-0-10-apt | TGTCACTGCGCGCCCGCCATGTTTACCAACGGCCAACCCTCAtAAGAAGGGATGATG CGGCAGTGGGCTTcatcAGGCTGTTCG |
| helix-0-11-apt | CCAGCGGTGCCGGTCTCCGTGGTGAAGGTGGGTAAAGGGATAtAAGAAGGGATGATG CGGCAGTGGGCTTcatcAGGCTGTTCG |
| helix-0-12-apt | TCAGCAAATCGTTAAAACAGCGGATCAACGCCAGCAGTTTCGtAAGAAGGGATGATG CGGCAGTGGGCTTcatcAGGCTGTTCG |
| helix-0-13-apt | AGGTTTCTTTGCTCCAGTTGGGCGGTTGAACTGTTTTATTTCtAAGAAGGGATGATG CGGCAGTGGGCTTcatcAGGCTGTTCG |
| helix-0-14-apt | TATGAGCCGGGTCAAAAAAGCCGCACAGGCCGGAATTTTAAAtAAGAAGGGATGATG CGGCAGTGGGCTTcatcAGGCTGTTCG |
| helix-0-15-apt | TTGCAGGCGCTTTCAAACGATGCTGATTGGAAGATAAGGGTGtAAGAAGGGATGATG CGGCAGTGGGCTTcatcAGGCTGTTCG |
| helix-0-16-apt | CAACCAGCTTACGGTCGTCTCGTCGCTGTGCATCTGATATTCtAAGAAGGGATGATG CGGCAGTGGGCTTcatcAGGCTGTTCG |
| helix-0-17-apt | CACCTCACTCgatgAGGGGTCTTtAATAACGTAGGTCTCGTTAAATAAGAATGAGGC CATTTGACG |
| helix-0-18-apt | CACCTCACTCgatgAGGGGTCTTtATCGCGCTTAAGACAGCCTGTTTAGTATGAACG GTATTCACC |
| helix-0-19-apt | CACCTCACTCgatgAGGGGTCTTtAGAAGATAATTTTCATAAAGCCAACGCTGCGGG AGCAACAGA |
| helix-0-20-apt | CACCTCACTCgatgAGGGGTCTTtATTTAACGTGAGTGATATTTAACAACGCTTTGA CGGTGGCAC |
| helix-0-21-apt | CACCTCACTCgatgAGGGGTCTTtGCCCCCTAACAGTGCGGAGTCCACTATTGCCGC GCCGCGAAC |
| helix-0-22-apt | CACCTCACTCgatgAGGGGTCTTtGTATTAAAGTGTACCGAAAAACCGTCTACTGGC AAACGAACC |
| helix-0-23-apt | CACCTCACTCgatgAGGGGTCTTtGCGGGGTACCGTTCTCACCCAAATCAAGAGAAA GGTATTAAC |
| helix-0-24-apt | CACCTCACTCgatgAGGGGTCTTtGGGTTGAATCCTCACTAAATCGGAACCCCCCCG ATCAAATGA |
| helix-0-25-apt | CACCTCACTCgatgAGGGGTCTTtTCAGGAGTAATAATGCGTGCCTGTTCTTTGCTG CGAACAATC |
| helix-0-26-apt | CACCTCACTCgatgAGGGGTCTTtGAACCGCAGCGGAGGATCCCCGGGTACCACGAT CCAATTTGT |
| helix-0-27-apt | CACCTCACTCgatgAGGGGTCTTtGCAAGCCATTTTCTCTGTTTCCTGTGTGCGGGT TACTCACGG |
| helix-0-28-apt | CACCTCACTCgatgAGGGGTCTTtTCACCAGACGATCTAACATACGAGCCGGTTACA CTTTTCTGC |

| | |
|---|---|
| helix-0-29-apt | CACCTCACTCgatgAGGGGTCTTtAACGCAACTTTTGCTATTTTGCGGATGGCTGGTAACGACATA |
| helix-0-30-apt | CACCTCACTCgatgAGGGGTCTTtTGCAATGAAGCTAATGTAGCTCAACATGGGGCGCGTTAGTGA |
| helix-0-31-apt | CACCTCACTCgatgAGGGGTCTTtAGAAAGGGCAAAGACTGGAAGTTTCATTCATCAGCCGGCAAA |
| helix-0-32-apt | CACCTCACTCgatgAGGGGTCTTtAACCGTTTAATAGTGAACGAGTAGATTTGGTGCCACCGGCCA |
| 10-HB modulator | CGAACAGCCTGAGTGAGGTG |
| helix-9-1-probebinding | TGTTACTACGAAGGTTAACATCCAATAAACAGTTGTTCAAATctcactcaatcactaccact |
| helix-9-2-probebinding | GAACCGATGAGGAAAAATTAAGCAATAATATGCAACCGGAAGctcactcaatcactaccact |
| helix-9-3-probebinding | TACAGACTAGCAACGTACCAAAAACATTCTTAATTAATTGCTctcactcaatcactaccact |
| helix-9-4-probebinding | ATCTTGAGATCGTCGCCACAGACAGCCCAAGTGTAACATTAActcactcaatcactaccact |
| helix-9-5-probebinding | CAAAGCTCGGTCGCTGTCGTCTTTCCAGTATCCGCAACCTGTctcactcaatcactaccact |
| helix-9-6-probebinding | ACACCAGCGACAATATTTTGCTAAACAAAATTCGTGGGAGAGctcactcaatcactaccact |
| helix-9-7-probebinding | TCAACTTTCGAGGTAGAAAGGAACAACTGTGAGCCTTTCACCctcactcaatcactaccact |
| helix-9-8-probebinding | GCTAATATCCAAAACGTTGAAAATCTCCACCGGGGCTGGCCCctcactcaatcactaccact |
| helix-9-9-probebinding | AATAATATTGAGGCCAGAATGGAAAGCGGGGTCGACCCCAGCctcactcaatcactaccact |
| helix-9-10-probebinding | ACCGAAGACCAGAACGTCATACATGGCTGATGGCCGGCAAAActcactcaatcactaccact |
| helix-9-11-probebinding | AGTTACCCTCAGAAAAGTTTTAACGGGGGTGGACTTTGAGTGctcactcaatcactaccact |
| helix-9-12-probebinding | ACCCAAAACCAGAGAATGGAAACAGTACTAATTTAATATAAActcactcaatcactaccact |
| helix-9-13-probebinding | ATGTTAGCTTATTATTGCTTCTGTAAATAGGGCTTACGACAActcactcaatcactaccact |
| helix-9-14-probebinding | TATAAAAAGACTGTATCCTTGAAAACATGTTATACTTTATCActcactcaatcactaccact |
| helix-9-15-probebinding | TCACAATACCGTAAAGAGTCAATAGTGAGGAATCACCATCCTctcactcaatcactaccact |

| | |
|---|---|
| helix-9-16-probebinding | ACAAAAGAGCAAGGTACCTTTTTAACCTCCGACCGCTGTCTTctcactcaatcactaccact |
| helix-0-1-probebinding | AAAGAGTCTGTCCACGCTCATGGAAATAGTTTGGAGAGAAACctcactcaatcactaccact |
| helix-0-2-probebinding | AATCCTGAGAAGTGATGGATTATTTACACTGATTATTACAAActcactcaatcactaccact |
| helix-0-3-probebinding | GGAGGCCGATTAAATAATAAAAGGGACAAACAAAGGAGCAAActcactcaatcactaccact |
| helix-0-4-probebinding | ATAACGTGCTTTCCGACCTGAAAGCGTAGTTATTATAATTACctcactcaatcactaccact |
| helix-0-5-probebinding | GCCGCTACAGGGCGAATGGCTATTAGTCTACAAACAGTTAATctcactcaatcactaccact |
| helix-0-6-probebinding | GGTCACGCTGCGCGCATCGCCATTAAAAGTCAATACATGAAActcactcaatcactaccact |
| helix-0-7-probebinding | GAAAGCGAAAGGAGAAACAGAGGTGAGGTAAAATAAGGATTActcactcaatcactaccact |
| helix-0-8-probebinding | TTGACGGGGAAAGCTGCCACGCTGAGAGAGTTGGCGTCGAGActcactcaatcactaccact |
| helix-0-9-probebinding | TGCGGCGGGCCGTTCACCTTGCTAACCTCGGATGAACCGTACctcactcaatcactaccact |
| helix-0-10-probebinding | TGTCACTGCGCGCCCGCCATGTTTACCAACGGCCAACCCTCActcactcaatcactaccact |
| helix-0-11-probebinding | CCAGCGGTGCCGGTCTCCGTGGTGAAGGTGGGTAAAGGGATActcactcaatcactaccact |
| helix-0-12-probebinding | TCAGCAAATCGTTAAAACAGCGGATCAACGCCAGCAGTTTCGctcactcaatcactaccact |
| helix-0-13-probebinding | AGGTTTCTTTGCTCCAGTTGGGCGGTTGAACTGTTTTATTTCctcactcaatcactaccact |
| helix-0-14-probebinding | TATGAGCCGGGTCAAAAAAGCCGCACAGGCCGGAATTTTAAActcactcaatcactaccact |
| helix-0-15-probebinding | TTGCAGGCGCTTTCAAACGATGCTGATTGGAAGATAAGGGTGctcactcaatcactaccact |
| helix-0-16-probebinding | CAACCAGCTTACGGTCGTCTCGTCGCTGTGCATCTGATATTCctcactcaatcactaccact |
| FAM-probe | agtggtagtgattgagtgag |
| helix-9-1-RNA1 | TGTTACTACGAAGGTTAACATCCAATAAACAGTTGTTCAAATtAAGAAGGGATGATGCGGCAGTGGGCTTcatcGGGTCGCCAT |
| helix-9-2-RNA1 | GAACCGATGAGGAAAAATTAAGCAATAATATGCAACCGGAAGtAAGAAGGGATGATGCGGCAGTGGGCTTcatcTAACCTGGTT |
| helix-9-3-RNA1 | TACAGACTAGCAACGTACCAAAAACATTCTTAATTAATTGCTtAAGAAGGGATGATGCGGCAGTGGGCTTcatcCTCAGCATAA |

| | |
|---|---|
| helix-9-4-RNA1 | ATCTTGAGATCGTCGCCACAGACAGCCCAAGTGTAACATTAAtAAGAAGGGATGATGCGGCAGTGGGCTTcatcCCATAATTAG |
| helix-9-5-RNA1 | CAAAGCTCGGTCGCTGTCGTCTTTCCAGTATCCGCAACCTGTtAAGAAGGGATGATGCGGCAGTGGGCTTcatcATCTCCTTCA |
| helix-9-6-RNA1 | ACACCAGCGACAATATTTTGCTAAACAAAATTCGTGGGAGAGtAAGAAGGGATGATGCGGCAGTGGGCTTcatcCCCCTTGAGC |
| helix-9-7-RNA1 | TCAACTTTCGAGGTAGAAAGGAACAACTGTGAGCCTTTCACCtAAGAAGGGATGATGCGGCAGTGGGCTTcatcCTTTGATGTA |
| helix-9-8-RNA1 | GCTAATATCCAAAACGTTGAAAATCTCCACCGGGGCTGGCCCtAAGAAGGGATGATGCGGCAGTGGGCTTcatcACAGTCATAG |
| helix-9-9-RNA1 | AATAATATTGAGGCCAGAATGGAAAGCGGGGTCGACCCCAGCtAAGAAGGGATGATGCGGCAGTGGGCTTcatcTGACTGGTCA |
| helix-9-10-RNA1 | ACCGAAGACCAGAACGTCATACATGGCTGATGGCCGGCAAAAtAAGAAGGGATGATGCGGCAGTGGGCTTcatcTTGAGAGATC |
| helix-9-11-RNA1 | AGTTACCCTCAGAAAAGTTTTAACGGGGGTGGACTTTGAGTGtAAGAAGGGATGATGCGGCAGTGGGCTTcatcATTATATCTT |
| helix-9-12-RNA1 | ACCCAAAACCAGAGAATGGAAACAGTACTAATTTAATATAAAtAAGAAGGGATGATGCGGCAGTGGGCTTcatcGACCAAGGAA |
| helix-9-13-RNA1 | ATGTTAGCTTATTATTGCTTCTGTAAATAGGGCTTACGACAAtAAGAAGGGATGATGCGGCAGTGGGCTTcatcGCAAGCTTGC |
| helix-9-14-RNA1 | TATAAAAAGACTGTATCCTTGAAAACATGTTATACTTTATCAtAAGAAGGGATGATGCGGCAGTGGGCTTcatcTCTGGCTTAT |
| helix-9-15-RNA1 | TCACAATACCGTAAAGAGTCAATAGTGAGGAATCACCATCCTtAAGAAGGGATGATGCGGCAGTGGGCTTcatcTCCTACAACA |
| helix-9-16-RNA1 | ACAAAAGAGCAAGGTACCTTTTTAACCTCCGACCGCTGTCTTtAAGAAGGGATGATGCGGCAGTGGGCTTcatcGATTCAAATC |
| helix-9-17-RNA1 | CCAGGGCTGCgatgAGGGGTCTTtATCGCGTTGTTTAGAACGAGGCGCAGACGGTAAAATACGTAA |
| helix-9-18-RNA1 | ATCAAGGTCAgatgAGGGGTCTTtCAAACTCGCTTTTGAACTTTGAAAGAGGTGAGGACTAAAGAC |
| helix-9-19-RNA1 | TTTCCAAATCgatgAGGGGTCTTtCCTTTTGCCCTCGTATAGGCTGGCTGACAAGACAGCATCGGA |
| helix-9-20-RNA1 | TCAGTCCTGTgatgAGGGGTCTTtTTGCGTTCAAAAGGCGGATATTCATTACAGTTAAAGGCCGCT |
| helix-9-21-RNA1 | ATGGCCTCCCgatgAGGGGTCTTtCGTGCCAGATTTAGCAGTGAATAAGGCTCCCACGCATAACCG |
| helix-9-22-RNA1 | ATTTATAGCCgatgAGGGGTCTTtGCGGTTTAACGGAAAGTAAATTGGGCTTGCTTGATACCGATA |
| helix-9-23-RNA1 | CTATTCAGTGgatgAGGGGTCTTtAGTGAGATCAGGACTGTGAATTACCTTATATCGGTTTATCAG |
| helix-9-24-RNA1 | GATAAAATCTgatgAGGGGTCTTtTGAGAGAGAACAAAGATAACCCACAAGAGCCTTGATATTCAA |

| | |
|---|---|
| helix-9-25-RNA1 | TGTCCCCTGTgatgAGGGGTCTTtAGGCGAAAACAGGGGAAACAATGAAATAGCCGCCAGCATTGA |
| helix-9-26-RNA1 | CCAGTTAAAGgatgAGGGGTCTTtTCCCTTACAAAAATTAAGAAAAGTAAGCGCCACCACCCTCAG |
| helix-9-27-RNA1 | GCCAGTGTCAgatgAGGGGTCTTtTTGTTCCATTTATCAACCGAGGAAACGCGCCTCCCTCAGAGC |
| helix-9-28-RNA1 | TATACTGCCTgatgAGGGGTCTTtGTACCGATCCAGAGGCATGATTAAGACTTTCATAATCAAAAT |
| helix-9-29-RNA1 | CTTTTCACCAgatgAGGGGTCTTtTAAACAATACAATTAGAAAATACATACAGGCATTTTCGGTCA |
| helix-9-30-RNA1 | TCCAACAAAGgatgAGGGGTCTTtACAATAGTTGAAGCAAAGACACCACGGAATCAAGTTTGCCTT |
| helix-9-31-RNA1 | CAAGGGCATAgatgAGGGGTCTTtAATTTACAACGCGAAAATTCATATGGTTAATGAAACCATCGA |
| helix-9-32-RNA1 | ACACAAACATgatgAGGGGTCTTtTCCTTATAGCAAGCTTCAACCGATTGAGACCAGTAGCACCAT |
| helix-0-1-RNA1 | AAAGAGTCTGTCCACGCTCATGGAAATAGTTTGGAGAGAAACtAAGAAGGGATGATGCGGCAGTGGGCTTcatcGGGTCGCCAT |
| helix-0-2-RNA1 | AATCCTGAGAAGTGATGGATTATTTACACTGATTATTACAAAtAAGAAGGGATGATGCGGCAGTGGGCTTcatcTAACCTGGTT |
| helix-0-3-RNA1 | GGAGGCCGATTAAATAATAAAAGGGACAAACAAAGGAGCAAAtAAGAAGGGATGATGCGGCAGTGGGCTTcatcCTCAGCATAA |
| helix-0-4-RNA1 | ATAACGTGCTTTCCGACCTGAAAGCGTAGTTATTATAATTACtAAGAAGGGATGATGCGGCAGTGGGCTTcatcCCATAATTAG |
| helix-0-5-RNA1 | GCCGCTACAGGGCGAATGGCTATTAGTCTACAAACAGTTAATtAAGAAGGGATGATGCGGCAGTGGGCTTcatcATCTCCTTCA |
| helix-0-6-RNA1 | GGTCACGCTGCGCGCATCGCCATTAAAAGTCAATACATGAAAtAAGAAGGGATGATGCGGCAGTGGGCTTcatcCCCCTTGAGC |
| helix-0-7-RNA1 | GAAAGCGAAAGGAGAAACAGAGGTGAGGTAAAATAAGGATTAtAAGAAGGGATGATGCGGCAGTGGGCTTcatcCTTTGATGTA |
| helix-0-8-RNA1 | TTGACGGGGAAAGCTGCCACGCTGAGAGAGTTGGCGTCGAGAtAAGAAGGGATGATGCGGCAGTGGGCTTcatcACAGTCATAG |
| helix-0-9-RNA1 | TGCGGCGGGCCGTTCACCTTGCTAACCTCGGATGAACCGTACtAAGAAGGGATGATGCGGCAGTGGGCTTcatcTGACTGGTCA |
| helix-0-10-RNA1 | TGTCACTGCGCGCCCGCCATGTTTACCAACGGCCAACCCTCAtAAGAAGGGATGATGCGGCAGTGGGCTTcatcTTGAGAGATC |
| helix-0-11-RNA1 | CCAGCGGTGCCGGTCTCCGTGGTGAAGGTGGGTAAAGGGATAtAAGAAGGGATGATGCGGCAGTGGGCTTcatcATTATATCTT |
| helix-0-12-RNA1 | TCAGCAAATCGTTAAAACAGCGGATCAACGCCAGCAGTTTCGtAAGAAGGGATGATGCGGCAGTGGGCTTcatcGACCAAGGAA |
| helix-0-13-RNA1 | AGGTTTCTTTGCTCCAGTTGGGCGGTTGAACTGTTTTATTTCtAAGAAGGGATGATGCGGCAGTGGGCTTcatcGCAAGCTTGC |

| | |
|---|---|
| helix-0-14-RNA1 | TATGAGCCGGGTCAAAAAAGCCGCACAGGCCGGAATTTTAAAtAAGAAGGGATGATGCGGCAGTGGGCTTcatcTCTGGCTTAT |
| helix-0-15-RNA1 | TTGCAGGCGCTTTCAAACGATGCTGATTGGAAGATAAGGGTGtAAGAAGGGATGATGCGGCAGTGGGCTTcatcTCCTACAACA |
| helix-0-16-RNA1 | CAACCAGCTTACGGTCGTCTCGTCGCTGTGCATCTGATATTCtAAGAAGGGATGATGCGGCAGTGGGCTTcatcGATTCAAATC |
| helix-0-17-RNA1 | CCAGGGCTGCgatgAGGGGTCTTtAATAACGTAGGTCTCGTTAAATAAGAATGAGGCCATTTGACG |
| helix-0-18-RNA1 | ATCAAGGTCAgatgAGGGGTCTTtATCGCGCTTAAGACAGCCTGTTTAGTATGAACGGTATTCACC |
| helix-0-19-RNA1 | TTTCCAAATCgatgAGGGGTCTTtAGAAGATAATTTTCATAAAGCCAACGCTGCGGGAGCAACAGA |
| helix-0-20-RNA1 | TCAGTCCTGTgatgAGGGGTCTTtATTTAACGTGAGTGATATTTAACAACGCTTTGACGGTGGCAC |
| helix-0-21-RNA1 | ATGGCCTCCCgatgAGGGGTCTTtGCCCCCTAACAGTGCGGAGTCCACTATTGCCGCGCCGCGAAC |
| helix-0-22-RNA1 | ATTTATAGCCgatgAGGGGTCTTtGTATTAAAGTGTACCGAAAAACCGTCTACTGGCAAACGAACC |
| helix-0-23-RNA1 | CTATTCAGTGgatgAGGGGTCTTtGCGGGGTACCGTTCTCACCCAAATCAAGAGAAAGGTATTAAC |
| helix-0-24-RNA1 | GATAAAATCTgatgAGGGGTCTTtGGGTTGAATCCTCACTAAATCGGAACCCCCCCGATCAAATGA |
| helix-0-25-RNA1 | TGTCCCCTGTgatgAGGGGTCTTtTCAGGAGTAATAATGCGTGCCTGTTCTTTGCTGCGAACAATC |
| helix-0-26-RNA1 | CCAGTTAAAGgatgAGGGGTCTTtGAACCGCAGCGGAGGATCCCCGGGTACCACGATCCAATTTGT |
| helix-0-27-RNA1 | GCCAGTGTCAgatgAGGGGTCTTtGCAAGCCATTTTCTCTGTTTCCTGTGTGCGGGTTACTCACGG |
| helix-0-28-RNA1 | TATACTGCCTgatgAGGGGTCTTtTCACCAGACGATCTAACATACGAGCCGGTTACACTTTTCTGC |
| helix-0-29-RNA1 | CTTTTCACCAgatgAGGGGTCTTtAACGCAACTTTTGCTATTTTGCGGATGGCTGGTAACGACATA |
| helix-0-30-RNA1 | TCCAACAAAGgatgAGGGGTCTTtTGCAATGAAGCTAATGTAGCTCAACATGGGGCGCGTTAGTGA |
| helix-0-31-RNA1 | CAAGGGCATAgatgAGGGGTCTTtAGAAAGGGCAAAGACTGGAAGTTTCATTCATCAGCCGGCAAA |
| helix-0-32-RNA1 | ACACAAACATgatgAGGGGTCTTtAACCGTTTAATAGTGAACGAGTAGATTTGGTGCCACCGGCCA |
| helix-9-1-RNA2 | TGTTACTACGAAGGTTAACATCCAATAAACAGTTGTTCAAATtAAGAAGGGATGATGCGGCAGTGGGCTTcatcCCTTCCCCAT |
| helix-9-2-RNA2 | GAACCGATGAGGAAAAATTAAGCAATAATATGCAACCGGAAGtAAGAAGGGATGATGCGGCAGTGGGCTTcatcTAAAAGCAGC |

| | |
|---|---|
| helix-9-3-RNA2 | TACAGACTAGCAACGTACCAAAAACATTCTTAATTAATTGCTtAAGAAGGGATGATGCGGCAGTGGGCTTcatcCCATGTAGTT |
| helix-9-4-RNA2 | ATCTTGAGATCGTCGCCACAGACAGCCCAAGTGTAACATTAAtAAGAAGGGATGATGCGGCAGTGGGCTTcatcTCTCAGCCTT |
| helix-9-5-RNA2 | CAAAGCTCGGTCGCTGTCGTCTTTCCAGTATCCGCAACCTGTtAAGAAGGGATGATGCGGCAGTGGGCTTcatcTGGAGGGATC |
| helix-9-6-RNA2 | ACACCAGCGACAATATTTTGCTAAACAAAATTCGTGGGAGAGtAAGAAGGGATGATGCGGCAGTGGGCTTcatcTGGTGAAGAC |
| helix-9-7-RNA2 | TCAACTTTCGAGGTAGAAAGGAACAACTGTGAGCCTTTCACCtAAGAAGGGATGATGCGGCAGTGGGCTTcatcGGGCAGAGAT |
| helix-9-8-RNA2 | GCTAATATCCAAAACGTTGAAAATCTCCACCGGGGCTGGCCCtAAGAAGGGATGATGCGGCAGTGGGCTTcatcGGCTGTTGTC |
| helix-9-9-RNA2 | AATAATATTGAGGCCAGAATGGAAAGCGGGGTCGACCCCAGCtAAGAAGGGATGATGCGGCAGTGGGCTTcatcTGACCTTGGC |
| helix-9-10-RNA2 | ACCGAAGACCAGAACGTCATACATGGCTGATGGCCGGCAAAAtAAGAAGGGATGATGCGGCAGTGGGCTTcatcTGGCAGTGAT |
| helix-9-11-RNA2 | AGTTACCCTCAGAAAAGTTTTAACGGGGGTGGACTTTGAGTGtAAGAAGGGATGATGCGGCAGTGGGCTTcatcGGAGAGCCCC |
| helix-9-12-RNA2 | ACCCAAAACCAGAGAATGGAAACAGTACTAATTTAATATAAAtAAGAAGGGATGATGCGGCAGTGGGCTTcatcGCTCAGGGAT |
| helix-9-13-RNA2 | ATGTTAGCTTATTATTGCTTCTGTAAATAGGGCTTACGACAAtAAGAAGGGATGATGCGGCAGTGGGCTTcatcCCACCACTGA |
| helix-9-14-RNA2 | TATAAAAAGACTGTATCCTTGAAAACATGTTATACTTTATCAtAAGAAGGGATGATGCGGCAGTGGGCTTcatcGCTTCACCAC |
| helix-9-15-RNA2 | TCACAATACCGTAAAGAGTCAATAGTGAGGAATCACCATCCTtAAGAAGGGATGATGCGGCAGTGGGCTTcatcAGGAGACCAC |
| helix-9-16-RNA2 | ACAAAAGAGCAAGGTACCTTTTTAACCTCCGACCGCTGTCTTtAAGAAGGGATGATGCGGCAGTGGGCTTcatcTGAGGGCAAT |
| helix-9-17-RNA2 | CCGACCTTCAgatgAGGGGTCTTtATCGCGTTGTTTAGAACGAGGCGCAGACGGTAAAATACGTAA |
| helix-9-18-RNA2 | TTACCAGAGTgatgAGGGGTCTTtCAAACTCGCTTTTGAACTTTGAAAGAGGTGAGGACTAAAGAC |
| helix-9-19-RNA2 | AACATGTAAAgatgAGGGGTCTTtCCTTTTGCCCTCGTATAGGCTGGCTGACAAGACAGCATCGGA |
| helix-9-20-RNA2 | AGCTTCCCGTgatgAGGGGTCTTtTTGCGTTCAAAAGGCGGATATTCATTACAGTTAAAGGCCGCT |
| helix-9-21-RNA2 | CACTTGATTTgatgAGGGGTCTTtCGTGCCAGATTTAGCAGTGAATAAGGCTCCCACGCATAACCG |
| helix-9-22-RNA2 | TTCTCCATGGgatgAGGGGTCTTtGCGGTTTAACGGAAAGTAAATTGGGCTTGCTTGATACCGATA |
| helix-9-23-RNA2 | TCAGCAGAGGgatgAGGGGTCTTtAGTGAGATCAGGACTGTGAATTACCTTATATCGGTTTATCAG |

| | |
|---|---|
| helix-9-24-RNA2 | ATGATCTTGAgatgAGGGGTCTTtTGAGAGAGAACAAAGATAACCCACAAGAGCCTTGATATTCAA |
| helix-9-25-RNA2 | TTGTCATGGAgatgAGGGGTCTTtAGGCGAAAACAGGGGAAACAATGAAATAGCCGCCAGCATTGA |
| helix-9-26-RNA2 | GTCTTCTGGGgatgAGGGGTCTTtTCCCTTACAAAAATTAAGAAAAGTAAGCGCCACCACCCTCAG |
| helix-9-27-RNA2 | ATGATGTTCTgatgAGGGGTCTTtTTGTTCCATTTATCAACCGAGGAAACGCGCCTCCCTCAGAGC |
| helix-9-28-RNA2 | TTCCCGTTCAgatgAGGGGTCTTtGTACCGATCCAGAGGCATGATTAAGACTTTCATAATCAAAAT |
| helix-9-29-RNA2 | CAGGTCAGGTgatgAGGGGTCTTtTAAACAATACAATTAGAAAATACATACAGGCATTTTCGGTCA |
| helix-9-30-RNA2 | TCCGACGCCTgatgAGGGGTCTTtACAATAGTTGAAGCAAAGACACCACGGAATCAAGTTTGCCTT |
| helix-9-31-RNA2 | TTGAAGTCAGgatgAGGGGTCTTtAATTTACAACGCGAAAATTCATATGGTTAATGAAACCATCGA |
| helix-9-32-RNA2 | AAGTGGTCGTgatgAGGGGTCTTtTCCTTATAGCAAGCTTCAACCGATTGAGACCAGTAGCACCAT |
| helix-0-1-RNA2 | AAAGAGTCTGTCCACGCTCATGGAAATAGTTTGGAGAGAAACtAAGAAGGGATGATGCGGCAGTGGGCTTcatcCCTTCCCCAT |
| helix-0-2-RNA2 | AATCCTGAGAAGTGATGGATTATTTACACTGATTATTACAAAtAAGAAGGGATGATGCGGCAGTGGGCTTcatcTAAAAGCAGC |
| helix-0-3-RNA2 | GGAGGCCGATTAAATAATAAAAGGGACAAACAAAGGAGCAAAtAAGAAGGGATGATGCGGCAGTGGGCTTcatcCCATGTAGTT |
| helix-0-4-RNA2 | ATAACGTGCTTTCCGACCTGAAAGCGTAGTTATTATAATTACtAAGAAGGGATGATGCGGCAGTGGGCTTcatcTCTCAGCCTT |
| helix-0-5-RNA2 | GCCGCTACAGGGCGAATGGCTATTAGTCTACAAACAGTTAATtAAGAAGGGATGATGCGGCAGTGGGCTTcatcTGGAGGGATC |
| helix-0-6-RNA2 | GGTCACGCTGCGCGCATCGCCATTAAAAGTCAATACATGAAAtAAGAAGGGATGATGCGGCAGTGGGCTTcatcTGGTGAAGAC |
| helix-0-7-RNA2 | GAAAGCGAAAGGAGAAACAGAGGTGAGGTAAAATAAGGATTAtAAGAAGGGATGATGCGGCAGTGGGCTTcatcGGGCAGAGAT |
| helix-0-8-RNA2 | TTGACGGGGAAAGCTGCCACGCTGAGAGAGTTGGCGTCGAGAtAAGAAGGGATGATGCGGCAGTGGGCTTcatcGGCTGTTGTC |
| helix-0-9-RNA2 | TGCGGCGGGCCGTTCACCTTGCTAACCTCGGATGAACCGTACtAAGAAGGGATGATGCGGCAGTGGGCTTcatcTGACCTTGGC |
| helix-0-10-RNA2 | TGTCACTGCGCGCCCGCCATGTTTACCAACGGCCAACCCTCAtAAGAAGGGATGATGCGGCAGTGGGCTTcatcTGGCAGTGAT |
| helix-0-11-RNA2 | CCAGCGGTGCCGGTCTCCGTGGTGAAGGTGGGTAAAGGGATAtAAGAAGGGATGATGCGGCAGTGGGCTTcatcGGAGAGCCCC |
| helix-0-12-RNA2 | TCAGCAAATCGTTAAAACAGCGGATCAACGCCAGCAGTTTCGtAAGAAGGGATGATGCGGCAGTGGGCTTcatcGCTCAGGGAT |

| | |
|---|---|
| helix-0-13-RNA2 | AGGTTTCTTTGCTCCAGTTGGGCGGTTGAACTGTTTTATTTCtAAGAAGGGATGATGCGGCAGTGGGCTTcatcCCACCACTGA |
| helix-0-14-RNA2 | TATGAGCCGGGTCAAAAAAGCCGCACAGGCCGGAATTTTAAAtAAGAAGGGATGATGCGGCAGTGGGCTTcatcGCTTCACCAC |
| helix-0-15-RNA2 | TTGCAGGCGCTTTCAAACGATGCTGATTGGAAGATAAGGGTGtAAGAAGGGATGATGCGGCAGTGGGCTTcatcAGGAGACCAC |
| helix-0-16-RNA2 | CAACCAGCTTACGGTCGTCTCGTCGCTGTGCATCTGATATTCtAAGAAGGGATGATGCGGCAGTGGGCTTcatcTGAGGGCAAT |
| helix-0-17-RNA2 | CCGACCTTCAgatgAGGGGTCTTtAATAACGTAGGTCTCGTTAAATAAGAATGAGGCCATTTGACG |
| helix-0-18-RNA2 | TTACCAGAGTgatgAGGGGTCTTtATCGCGCTTAAGACAGCCTGTTTAGTATGAACGGTATTCACC |
| helix-0-19-RNA2 | AACATGTAAAgatgAGGGGTCTTtAGAAGATAATTTTCATAAAGCCAACGCTGCGGGAGCAACAGA |
| helix-0-20-RNA2 | AGCTTCCCGTgatgAGGGGTCTTtATTTAACGTGAGTGATATTTAACAACGCTTTGACGGTGGCAC |
| helix-0-21-RNA2 | CACTTGATTTgatgAGGGGTCTTtGCCCCCTAACAGTGCGGAGTCCACTATTGCCGCGCCGCGAAC |
| helix-0-22-RNA2 | TTCTCCATGGgatgAGGGGTCTTtGTATTAAAGTGTACCGAAAAACCGTCTACTGGCAAACGAACC |
| helix-0-23-RNA2 | TCAGCAGAGGgatgAGGGGTCTTtGCGGGGTACCGTTCTCACCCAAATCAAGAGAAAGGTATTAAC |
| helix-0-24-RNA2 | ATGATCTTGAgatgAGGGGTCTTtGGGTTGAATCCTCACTAAATCGGAACCCCCCGATCAAATGA |
| helix-0-25-RNA2 | TTGTCATGGAgatgAGGGGTCTTtTCAGGAGTAATAATGCGTGCCTGTTCTTTGCTGCGAACAATC |
| helix-0-26-RNA2 | GTCTTCTGGGgatgAGGGGTCTTtGAACCGCAGCGGAGGATCCCCGGGTACCACGATCCAATTTGT |
| helix-0-27-RNA2 | ATGATGTTCTgatgAGGGGTCTTtGCAAGCCATTTTCTCTGTTTCCTGTGTGCGGGTTACTCACGG |
| helix-0-28-RNA2 | TTCCCGTTCAgatgAGGGGTCTTtTCACCAGACGATCTAACATACGAGCCGGTTACACTTTTCTGC |
| helix-0-29-RNA2 | CAGGTCAGGTgatgAGGGGTCTTtAACGCAACTTTTGCTATTTTGCGGATGGCTGGTAACGACATA |
| helix-0-30-RNA2 | TCCGACGCCTgatgAGGGGTCTTtTGCAATGAAGCTAATGTAGCTCAACATGGGGCGCGTTAGTGA |
| helix-0-31-RNA2 | TTGAAGTCAGgatgAGGGGTCTTtAGAAAGGGCAAAGACTGGAAGTTTCATTCATCAGCCGGCAAA |
| helix-0-32-RNA2 | AAGTGGTCGTgatgAGGGGTCTTtAACCGTTTAATAGTGAACGAGTAGATTTGGTGCCACCGGCCA |
| helix-9-1-RNA3 | TGTTACTACGAAGGTTAACATCCAATAAACAGTTGTTCAAATtAAGAAGGGATGATGCGGCAGTGGGCTTcatcTCACAGACAT |

| | |
|---|---|
| helix-9-2-RNA3 | GAACCGATGAGGAAAAATTAAGCAATAATATGCAACCGGAAGtAAGAAGGGATGATGCGGCAGTGGGCTTcatcTTTTAATATC |
| helix-9-3-RNA3 | TACAGACTAGCAACGTACCAAAAACATTCTTAATTAATTGCTtAAGAAGGGATGATGCGGCAGTGGGCTTcatcTTTTGGGTGT |
| helix-9-4-RNA3 | ATCTTGAGATCGTCGCCACAGACAGCCCAAGTGTAACATTAAtAAGAAGGGATGATGCGGCAGTGGGCTTcatcTGGCACAAAG |
| helix-9-5-RNA3 | CAAAGCTCGGTCGCTGTCGTCTTTCCAGTATCCGCAACCTGTtAAGAAGGGATGATGCGGCAGTGGGCTTcatcTATGAAATAT |
| helix-9-6-RNA3 | ACACCAGCGACAATATTTTGCTAAACAAAATTCGTGGGAGAGtAAGAAGGGATGATGCGGCAGTGGGCTTcatcCTGTTTGAAC |
| helix-9-7-RNA3 | TCAACTTTCGAGGTAGAAAGGAACAACTGTGAGCCTTTCACCtAAGAAGGGATGATGCGGCAGTGGGCTTcatcTGCCCCTCTT |
| helix-9-8-RNA3 | GCTAATATCCAAAACGTTGAAAATCTCCACCGGGGCTGGCCCtAAGAAGGGATGATGCGGCAGTGGGCTTcatcCATCCTCAAA |
| helix-9-9-RNA3 | AATAATATTGAGGCCAGAATGGAAAGCGGGGTCGACCCCAGCtAAGAAGGGATGATGCGGCAGTGGGCTTcatcTAACATTGTG |
| helix-9-10-RNA3 | ACCGAAGACCAGAACGTCATACATGGCTGATGGCCGGCAAAAtAAGAAGGGATGATGCGGCAGTGGGCTTcatcGGCCATTATT |
| helix-9-11-RNA3 | AGTTACCCTCAGAAAAGTTTTAACGGGGGTGGACTTTGAGTGtAAGAAGGGATGATGCGGCAGTGGGCTTcatcTGATGAAGAA |
| helix-9-12-RNA3 | ACCCAAAACCAGAGAATGGAAACAGTACTAATTTAATATAAAtAAGAAGGGATGATGCGGCAGTGGGCTTcatcTCATGTCCAA |
| helix-9-13-RNA3 | ATGTTAGCTTATTATTGCTTCTGTAAATAGGGCTTACGACAAtAAGAAGGGATGATGCGGCAGTGGGCTTcatcCATCTATTAC |
| helix-9-14-RNA3 | TATAAAAAGACTGTATCCTTGAAAACATGTTATACTTTATCAtAAGAAGGGATGATGCGGCAGTGGGCTTcatcACTTCTCCAA |
| helix-9-15-RNA3 | TCACAATACCGTAAAGAGTCAATAGTGAGGAATCACCATCCTtAAGAAGGGATGATGCGGCAGTGGGCTTcatcGAGGTCGGTA |
| helix-9-16-RNA3 | ACAAAAGAGCAAGGTACCTTTTTAACCTCCGACCGCTGTCTTtAAGAAGGGATGATGCGGCAGTGGGCTTcatcTAGTTATGTC |
| helix-9-17-RNA3 | GTATGCAGTGgatgAGGGGTCTTtATCGCGTTGTTTAGAACGAGGCGCAGACGGTAAAATACGTAA |
| helix-9-18-RNA3 | AAGACTTCAAgatgAGGGGTCTTtCAAACTCGCTTTTGAACTTTGAAAGAGGTGAGGACTAAAGAC |
| helix-9-19-RNA3 | TTCTCACATGgatgAGGGGTCTTtCCTTTTGCCCTCGTATAGGCTGGCTGACAAGACAGCATCGGA |
| helix-9-20-RNA3 | TAGTAATTACgatgAGGGGTCTTtTTGCGTTCAAAAGGCGGATATTCATTACAGTTAAAGGCCGCT |
| helix-9-21-RNA3 | TTGATATTCCgatgAGGGGTCTTtCGTGCCAGATTTAGCAGTGAATAAGGCTCCCACGCATAACCG |
| helix-9-22-RNA3 | GTTGGATCTCgatgAGGGGTCTTtGCGGTTTAACGGAAAGTAAATTGGGCTTGCTTGATACCGATA |

| | |
|---|---|
| helix-9-23-RNA3 | CAAATAACGTgatgAGGGGTCTTtAGTGAGATCAGGACTGTGAATTACCTTATATCGGTTTATCAG |
| helix-9-24-RNA3 | TCACTGTATTgatgAGGGGTCTTtTGAGAGAGAACAAAGATAACCCACAAGAGCCTTGATATTCAA |
| helix-9-25-RNA3 | ACAACACCTCgatgAGGGGTCTTtAGGCGAAAACAGGGGAAACAATGAAATAGCCGCCAGCATTGA |
| helix-9-26-RNA3 | TTGGTGTTCGgatgAGGGGTCTTtTCCCTTACAAAAATTAAGAAAAGTAAGCGCCACCACCCTCAG |
| helix-9-27-RNA3 | TTGCCATAGGgatgAGGGGTCTTtTTGTTCCATTTATCAACCGAGGAAACGCGCCTCCCTCAGAGC |
| helix-9-28-RNA3 | ACGGTGTATTgatgAGGGGTCTTtGTACCGATCCAGAGGCATGATTAAGACTTTCATAATCAAAAT |
| helix-9-29-RNA3 | GTTTCCAGACgatgAGGGGTCTTtTAAACAATACAATTAGAAAATACATACAGGCATTTTCGGTCA |
| helix-9-30-RNA3 | TTTACTGGCAgatgAGGGGTCTTtACAATAGTTGAAGCAAAGACACCACGGAATCAAGTTTGCCTT |
| helix-9-31-RNA3 | ACATCATTAAgatgAGGGGTCTTtAATTTACAACGCGAAAATTCATATGGTTAATGAAACCATCGA |
| helix-9-32-RNA3 | TTGGCATGAAgatgAGGGGTCTTtTCCTTATAGCAAGCTTCAACCGATTGAGACCAGTAGCACCAT |
| helix-0-1-RNA3 | AAAGAGTCTGTCCACGCTCATGGAAATAGTTTGGAGAGAAACtAAGAAGGGATGATGCGGCAGTGGGCTTcatcTCACAGACAT |
| helix-0-2-RNA3 | AATCCTGAGAAGTGATGGATTATTTACACTGATTATTACAAAtAAGAAGGGATGATGCGGCAGTGGGCTTcatcTTTTAATATC |
| helix-0-3-RNA3 | GGAGGCCGATTAAATAATAAAAGGGACAAACAAAGGAGCAAAtAAGAAGGGATGATGCGGCAGTGGGCTTcatcTTTTGGGTGT |
| helix-0-4-RNA3 | ATAACGTGCTTTCCGACCTGAAAGCGTAGTTATTATAATTACtAAGAAGGGATGATGCGGCAGTGGGCTTcatcTGGCACAAAG |
| helix-0-5-RNA3 | GCCGCTACAGGGCGAATGGCTATTAGTCTACAAACAGTTAATtAAGAAGGGATGATGCGGCAGTGGGCTTcatcTATGAAATAT |
| helix-0-6-RNA3 | GGTCACGCTGCGCGCATCGCCATTAAAAGTCAATACATGAAAtAAGAAGGGATGATGCGGCAGTGGGCTTcatcCTGTTTGAAC |
| helix-0-7-RNA3 | GAAAGCGAAAGGAGAAACAGAGGTGAGGTAAAATAAGGATTAtAAGAAGGGATGATGCGGCAGTGGGCTTcatcTGCCCCTCTT |
| helix-0-8-RNA3 | TTGACGGGGAAAGCTGCCACGCTGAGAGAGTTGGCGTCGAGAtAAGAAGGGATGATGCGGCAGTGGGCTTcatcCATCCTCAAA |
| helix-0-9-RNA3 | TGCGGCGGGCCGTTCACCTTGCTAACCTCGGATGAACCGTACtAAGAAGGGATGATGCGGCAGTGGGCTTcatcTAACATTGTG |
| helix-0-10-RNA3 | TGTCACTGCGCGCCCGCCATGTTTACCAACGGCCAACCCTCAtAAGAAGGGATGATGCGGCAGTGGGCTTcatcGGCCATTATT |
| helix-0-11-RNA3 | CCAGCGGTGCCGGTCTCCGTGGTGAAGGTGGGTAAAGGGATAtAAGAAGGGATGATGCGGCAGTGGGCTTcatcTGATGAAGAA |

| | |
|---|---|
| helix-0-12-RNA3 | TCAGCAAATCGTTAAAACAGCGGATCAACGCCAGCAGTTTCGtAAGAAGGGATGATGCGGCAGTGGGCTTcatcTCATGTCCAA |
| helix-0-13-RNA3 | AGGTTTCTTTGCTCCAGTTGGGCGGTTGAACTGTTTTATTTCtAAGAAGGGATGATGCGGCAGTGGGCTTcatcCATCTATTAC |
| helix-0-14-RNA3 | TATGAGCCGGGTCAAAAAAGCCGCACAGGCCGGAATTTTAAAtAAGAAGGGATGATGCGGCAGTGGGCTTcatcACTTCTCCAA |
| helix-0-15-RNA3 | TTGCAGGCGCTTTCAAACGATGCTGATTGGAAGATAAGGGTGtAAGAAGGGATGATGCGGCAGTGGGCTTcatcGAGGTCGGTA |
| helix-0-16-RNA3 | CAACCAGCTTACGGTCGTCTCGTCGCTGTGCATCTGATATTCtAAGAAGGGATGATGCGGCAGTGGGCTTcatcTAGTTATGTC |
| helix-0-17-RNA3 | GTATGCAGTGgatgAGGGGTCTTtAATAACGTAGGTCTCGTTAAATAAGAATGAGGCCATTTGACG |
| helix-0-18-RNA3 | AAGACTTCAAgatgAGGGGTCTTtATCGCGCTTAAGACAGCCTGTTTAGTATGAACGGTATTCACC |
| helix-0-19-RNA3 | TTCTCACATGgatgAGGGGTCTTtAGAAGATAATTTTCATAAAGCCAACGCTGCGGGAGCAACAGA |
| helix-0-20-RNA3 | TAGTAATTACgatgAGGGGTCTTtATTTAACGTGAGTGATATTTAACAACGCTTTGACGGTGGCAC |
| helix-0-21-RNA3 | TTGATATTCCgatgAGGGGTCTTtGCCCCCTAACAGTGCGGAGTCCACTATTGCCGCGCCGCGAAC |
| helix-0-22-RNA3 | GTTGGATCTCgatgAGGGGTCTTtGTATTAAAGTGTACCGAAAAACCGTCTACTGGCAAACGAACC |
| helix-0-23-RNA3 | CAAATAACGTgatgAGGGGTCTTtGCGGGGTACCGTTCTCACCCAAATCAAGAGAAAGGTATTAAC |
| helix-0-24-RNA3 | TCACTGTATTgatgAGGGGTCTTtGGGTTGAATCCTCACTAAATCGGAACCCCCCCGATCAAATGA |
| helix-0-25-RNA3 | ACAACACCTCgatgAGGGGTCTTtTCAGGAGTAATAATGCGTGCCTGTTCTTTGCTGCGAACAATC |
| helix-0-26-RNA3 | TTGGTGTTCGgatgAGGGGTCTTtGAACCGCAGCGGAGGATCCCCGGGTACCACGATCCAATTTGT |
| helix-0-27-RNA3 | TTGCCATAGGgatgAGGGGTCTTtGCAAGCCATTTTCTCTGTTTCCTGTGTGCGGGTTACTCACGG |
| helix-0-28-RNA3 | ACGGTGTATTgatgAGGGGTCTTtTCACCAGACGATCTAACATACGAGCCGGTTACACTTTTCTGC |
| helix-0-29-RNA3 | GTTTCCAGACgatgAGGGGTCTTtAACGCAACTTTTGCTATTTTGCGGATGGCTGGTAACGACATA |
| helix-0-30-RNA3 | TTTACTGGCAgatgAGGGGTCTTtTGCAATGAAGCTAATGTAGCTCAACATGGGGCGCGTTAGTGA |
| helix-0-31-RNA3 | ACATCATTAAgatgAGGGGTCTTtAGAAAGGGCAAAGACTGGAAGTTTCATTCATCAGCCGGCAAA |
| helix-0-32-RNA3 | TTGGCATGAAgatgAGGGGTCTTtAACCGTTTAATAGTGAACGAGTAGATTTGGTGCCACCGGCCA |

| | |
|---|---|
| Short RNA modulator | GAAGUGUAUUCCAGGCGAGCGACAUUUGAC |
| RNA-1 | AUGGCGACCCGCAGCCCUGGCGUCGUGAUUAGUGAUGAUGAACCAGGUUAUGACCUU<br>GAUUUAUUUUGCAUACCUAAUCAUUAUGCUGAGGAUUUGGAAAGGGUGUUUAUUCCU<br>CAUGGACUAAUUAUGGACAGGACUGAACGUCUUGCUCGAGAUGUGAUGAAGGAGAUG<br>GGAGGCCAUCACAUUGUAGCCCUCUGUGUGCUCAAGGGGGGCUAUAAAUUCUUUGCU<br>GACCUGCUGGAUUACAUCAAAGCACUGAAUAGAAAUAGUGAUAGAUCCAUUCCUAUG<br>ACUGUAGAUUUUAUCAGACUGAAGAGCUAUUGUAAUGACCAGUCAACAGGGGACAUA<br>AAAGUAAUUGGUGGAGAUGAUCUCUCAACUUUAACUGGAAAGAAUGUCUUGAUUGUG<br>GAAGAUAUAAUUGACACUGGCAAAACAAUGCAGACUUUGCUUUCCUUGGUCAGGCAG<br>UAUAAUCCAAAGAUGGUCAAGGUCGCAAGCUUGCUGGUGAAAAGGACCCCACGAAGU<br>GUUGGAUAUAAGCCAGACUUUGUUGGAUUUGAAAUUCCAGACAAGUUUGUUGUAGGA<br>UAUGCCCUUGACUAUAAUGAAUACUUCAGGGAUUUGAAUCAUGUUUGUGUCAUUAGU<br>GAAACUGGAAAAGCAAAAUACAAAGCCUAA |
| RNA-2 | AUGGGGAAGGUGAAGGUCGGAGUCAACGGAUUUGGUCGUAUUGGGCGCCUGGUCACC<br>AGGGCUGCUUUUAACUCUGGUAAAGUGGAUAUUGUUGCCAUCAAUGACCCCUUCAUU<br>GACCUCAACUACAUGGUUUACAUGUUCCAAUAUGAUUCCACCCAUGGCAAAUUCCAU<br>GGCACCGUCAAGGCUGAGAACGGGAAGCUUGUCAUCAAUGGAAAUCCCAUCACCAUC<br>UUCCAGGAGCGAGAUCCCUCCAAAAUCAAGUGGGGCGAUGCUGGCGCUGAGUACGUC<br>GUGGAGUCCACUGGCGUCUUCACCACCAUGGAGAAGGCUGGGGCUCAUUUGCAGGGG<br>GGAGCCAAAAGGGUCAUCAUCUCUGCCCCCUCUGCUGAUGCCCCCAUGUUCGUCAUG<br>GGUGUGAACCAUGAGAAGUAUGACAACAGCCUCAAGAUCAUCAGCAAUGCCUCCUGC<br>ACCACCAACUGCUUAGCACCCCUGGCCAAGGUCAUCCAUGACAACUUUGGUAUCGUG<br>GAAGGACUCAUGACCACAGUCCAUGCCAUCACUGCCACCCAGAAGACUGUGGAUGGC<br>CCCUCCGGGAAACUGUGGCGUGAUGGCCGCGGGGCUCUCCAGAACAUCAUCCCUGCC<br>UCUACUGGCGCUGCCAAGGCUGUGGGCAAGGUCAUCCCUGAGCUGAACGGGAAGCUC<br>ACUGGCAUGGCCUUCCGUGUCCCCACUGCCAACGUGUCAGUGGUGGACCUGACCUGC<br>CGUCUAGAAAAACCUGCCAAAUAUGAUGACAUCAAGAAGGUGGUGAAGCAGGCGUCG<br>GAGGGCCCCCUCAAGGGCAUCCUGGGCUACACUGAGCACCAGGUGGUCUCCUCUGAC<br>UUCAACAGCGACACCCACUCCUCCACCUUUGACGCUGGGGCUGGCAUUGCCCUCAAC<br>GACCACUUUGUCAAGCUCAUUUCCUGGU |
| RNA-3 | AUGUCUGUGACACUGCAUACAGAUGUAGGUGAUAUUAAAAUUGAAGUCUUCUGUGAG<br>AGGACACCCAAAACAUGUGAGAAUUUCUUGGCUCUUUGUGCCAGUAAUUACUACAAU<br>GGCUGUAUAUUUCAUAGGAAUAUCAAGGGUUUCAUGGUUCAAACAGGAGAUCCAACA<br>GGAACUGGAAAGAGGGGCAACGUUAUUUGGGGCAAGAAGUUUGAGGAUGAAUACAGU<br>GAAAUUCUUAAGCACAAUGUUAGAGGUGUUGUAUCUAUGGCUAAUAAUGGCCCGAAC<br>ACCAAUGGAUCUCAGUUCUUCAUCACCUAUGGCAAACAGCCACAUUUGGACAUGAAA<br>UACACCGUAUUUGGAAAGGUAAUAGAUGGUCUGGAAACUCUAGAUGAGUUGGAGAAG<br>UUGCCAGUAAAUGAGAAGACAUACCGACCUCUUAAUGAUGUACACAUUAAGGACAUA<br>ACUAUUCAUGCCAACCCAUUUGCUCAGUAG |

#### References.

[1] Zadeh JN, Steenberg CD, Bois JS, Wolfe BR, Pierce MB, Khan AR, Dirks RM, Pierce NA. NUPACK: analysis and design of nucleic acid systems. J Comput Chem. 2011 Jan 15;32(1):170-3.

[2] Fornace ME, Huang J, Newman CT, Porubsky NJ, Pierce MB, Pierce NA. NUPACK: analysis and design of nucleic acid structures, devices, and systems. ChemRxiv, 2022.

[3] Douglas SM, Marblestone AH, Teerapittayanon S, Vazquez A, Church GM, Shih WM. Rapid prototyping of 3D DNA-origami shapes with caDNAno. Nucleic Acids Res. 2009 Aug;37(15):5001-6.

[4] Doty D, Lee BL, and Stérin T. scadnano: a browser-based, scriptable tool for designing DNA nanostructures. In 26th International Conference on DNA Computing and Molecular Programming (DNA 26). Leibniz International Proceedings in Informatics (LIPIcs), 2020 Sep;174(9):1-17.
